# Longitudinal and multimodal auditing of tumor adaptation to CDK4/6 inhibitors in HR+ metastatic breast cancers

**DOI:** 10.1101/2023.09.27.557464

**Authors:** Allison L. Creason, Jay Egger, Cameron Watson, Shamilene Sivagnanam, Koei Chin, Kevin MacPherson, Jia-Ren Lin, Yu-An Chen, Brett E. Johnson, Heidi S. Feiler, Danielle Galipeau, Nicholas E. Navin, Emek Demir, Young Hwan Chang, Christopher L. Corless, Zahi I. Mitri, Peter K. Sorger, George V. Thomas, Lisa M. Coussens, Andrew C. Adey, Joe W. Gray, Gordon B. Mills, Jeremy Goecks

## Abstract

CDK4/6 inhibitors (CDK4/6i) have transformed the treatment of hormone receptor-positive (HR+), HER2-negative (HR+) breast cancers as they are effective across all clinicopathological, age, and ethnicity subgroups for metastatic HR+ breast cancer. In metastatic ER+ breast cancer, CDK4/6i lead to strong and consistent improvement in survival across different lines of therapy. To improve understanding of how metastatic HR+ breast cancers become refractory to CDK4/6i, we have created a multimodal and longitudinal tumor atlas to investigate therapeutic adaptations in malignant cells and in the tumor immune microenvironment. This atlas is part of the NCI Cancer Moonshot Human Tumor Atlas Network and includes seven pairs of pre- and on-progression biopsies from five metastatic HR+ breast cancer patients treated with CDK4/6i. Biopsies were profiled with bulk genomics, transcriptomics, and proteomics as well as single-cell ATAC-seq and multiplex tissue imaging for spatial, single-cell resolution. These molecular datasets were then linked with detailed clinical metadata to create an atlas for understanding tumor adaptations during therapy. Analysis of our atlas datasets suggests a diverse set of tumor adaptations to CDK4/6i therapy. Malignant cells may adapt to therapy via mTORC1 activation, cell cycle bypass, and increased replication stress. The tumor immune microenvironment displayed evidence of both immune activation and immune suppression during therapy. Together, our metastatic ER+ breast cancer atlas represents a rich multimodal resource to better understand HR+ breast cancer tumor therapeutic adaptations to CDK4/6i therapy.

## Introduction

Hormone receptor-positive (HR+) breast cancer is the predominant subtype of breast cancer and is defined by its malignant cells that have receptors for the hormones estrogen or progesterone. HR+ breast cancer accounts for 70% of all breast cancers and has a 5-year survival rate of 31% upon metastasis. Patients with metastatic HR+ breast cancer receive a multitude of different endocrine, targeted, and chemotherapies, deployed over many years(Waks and Winer 2019). Among the most common therapies used in metastatic HR+ breast cancers are cyclin dependent kinase 4/6 inhibitors (CDK4/6i), which have transformed care of this disease (Dean et al. 2010; Pernas et al. 2018). CDK4/6i are targeted molecular cytostatic therapies that inhibit cell cycle progression and proliferation in tumors by impeding activity of the CDK4/6-CCND1 axis. As front-line therapy for metastatic HR+ breast cancer, the combination of CDK4/6i and endocrine therapy (ET) nearly double progression-free survival rates compared to ET alone (Janni et al. 2017; Finn et al. 2016; Goetz et al. 2015).

Like other targeted molecular therapies, overcoming resistance to CDK4/6i is a key ongoing challenge for the cancer research community. HR+ breast cancers may be intrinsically resistant to CDK4/6i or may acquire resistance during therapy. Given the clinical significance of CDK4/6i in treating HR+ breast cancer, it is critical to understand mechanisms underlying resistance to these agents. Examination of clinical specimens via bulk DNA-sequencing (Wander et al. 2020), bulk RNA-sequencing (Zhu et al. 2022), and single-cell RNA-sequencing (Griffiths et al. 2021) have identified genomic alterations and dysregulated oncogenic signaling as contributors to CDK4/6i resistance. Common aberrations include loss-of-function alterations of RB1 (Herrera-Abreu et al. 2016; Condorelli et al. 2018; O’Leary et al. 2018; Wander et al. 2020), and amplification or overexpression of CDK6 (Yang et al. 2017) and CCNE1/2 (Caldon et al. 2012; Herrera-Abreu et al. 2016; Wander et al. 2020); these aberrations mitigate efficacy of CDK4/6i by alleviating repression of cell cycle activity and proliferation. Moreover, these studies have further implicated mTOR (Xu et al. 2021; Knudsen et al. 2019; Romano et al. 2018), JNK/MAPK (Griffiths et al. 2021), and Hippo (Z. Li et al. 2018) pathway activation in CDK4/6i resistance. Collectively, upregulation of these pathways drives proliferation by promoting CDK4/6-independent cell cycle progression, as well as protumoral tumor immune microenvironment (TIME) dynamics (Scirocchi et al. 2022; Zhu et al. 2022), including inflammatory programs that may also contribute to CDK4/6i resistance.

In previous work, we presented an omic and multidimensional spatial (OMS) case study of a metastatic HR+ breast cancer patient treated with CDK4/6i and other therapies (Johnson et al. 2022). That study was created by linking detailed, longitudinal clinical metadata to clinical and exploratory multiscale molecular data analyses from four serial biopsies. Here we build on that study to create a data atlas that includes a cohort of 5 patients and 7 pre- and post-treatment biopsy pairs from metastatic HR+ breast cancer patients receiving CDK4/6i and/or ET. This atlas includes the founding patient in our original case study, and this atlas was created as part of the NCI Cancer Moonshot Human Tumor Atlas Network (HTAN) (Mitri et al. 2018; Rozenblatt-Rosen et al. 2020). Each biopsy in our atlas was profiled using a comprehensive suite of bulk and single-cell molecular assays, including several spatial omics assays, and linked with clinical metadata to inform analysis of the molecular datasets. Our atlas provides a rich resource for examining molecular changes within the tumor ecosystem, including genome, transcriptome, and proteome following exposure to CDK4/6i therapy, thus enabling investigation of therapeutic adaptations in both malignant cells and the TIME. Malignant cell adaptations observed during therapy include genetic and epigenetic changes and increases in cell cycle activity. TIME changes observed include chronic inflammation and/or T cell suppression, and stromal tumor microenvironment barriers. This detailed atlas of metastatic breast cancers on CDK4/6i therapy is a unique resource for identifying candidate tumor adaptations to therapy and corresponding therapeutic opportunities to combat resistance. All atlas raw data and derived datasets are available from the HTAN data portal at https://data.humantumoratlas.org/.

## Results

### A Longitudinal and Multimodal Atlas of HR+ Metastatic Breast Cancers Treated with CDK4/6i

An atlas of metastatic HR+ breast cancers treated with a CDK4/6i was created by assaying paired biopsies with six complementary molecular assays (**Figure 1a**) and connecting the data generated from these assays with detailed clinical metadata. Biopsy pairs were obtained from the same patient with one biopsy taken prior to starting CDK4/6i therapy, and a second biopsy taken after the CDK4/6i therapy had been discontinued due to disease progression. Biopsy pairs were obtained from a cohort of five female patients under the IRB-approved observational study “Molecular Mechanisms of Tumor Evolution and Resistance to Therapy” as part of HTAN (Mitri et al. 2018; Rozenblatt-Rosen et al. 2020). Patients received one of two CDK4/6i therapies, palbociclib or abemaciclib, and an ET. ET used with the CDK4/6i were either an aromatase inhibitor or fulvestrant. Patients remained on therapy until progression occurred based on physician determination.

**Figure 1:**
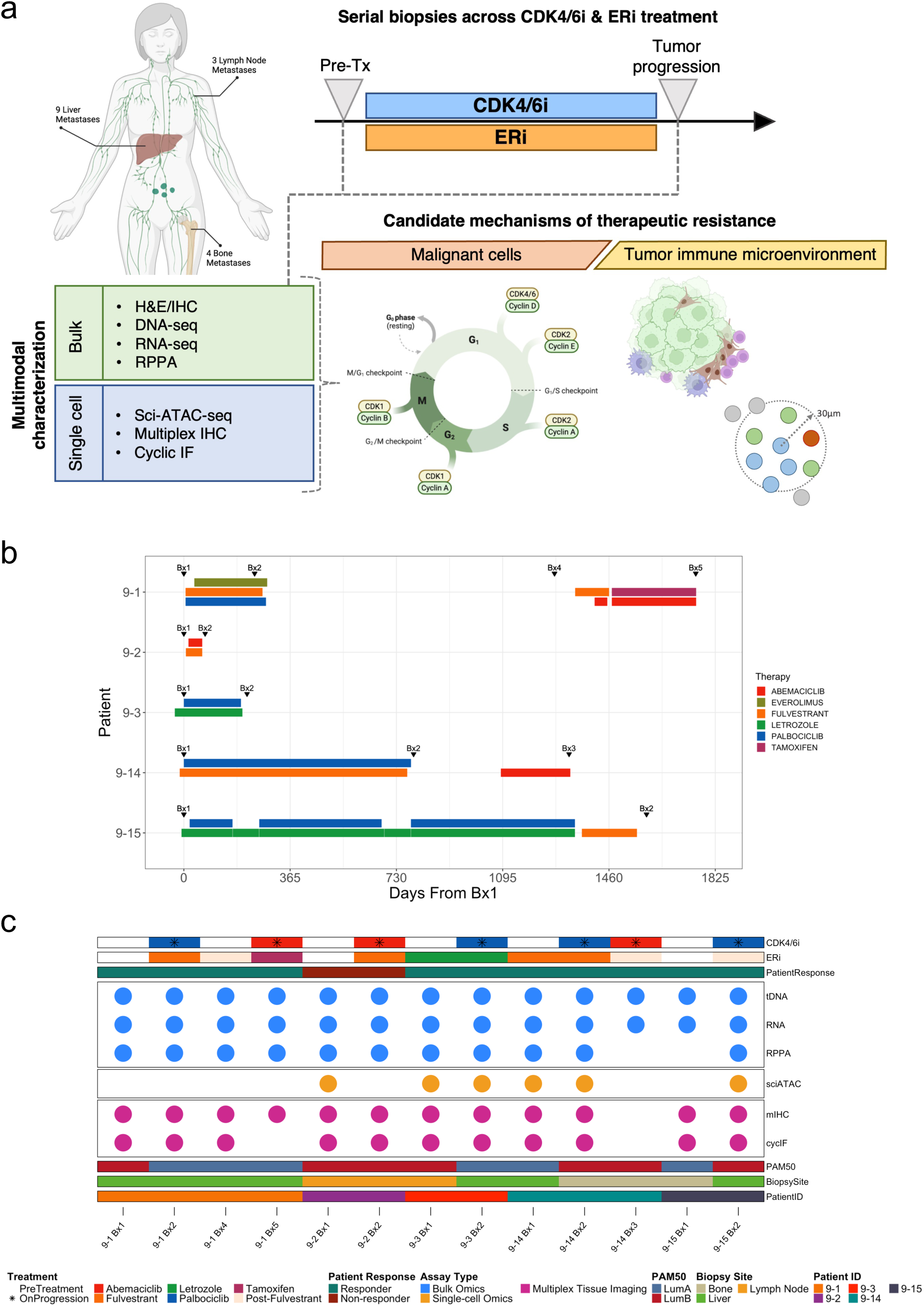
A longitudinal and multimodal atlas of five metastatic HR+ breast cancers on CDK4/6i + ET to understand how tumors adapt to this therapy combination. a) Overview of the atlas study design and data collection. Two biopsies were collected from each tumor: (1) before treatment and (2) after treatment was stopped due to disease progression. Disease progression indicates resistance to therapy. Three bulk molecular assays and three single-cell assays were applied to each biopsy, and data from these biopsies was used to identify malignant cell and TIME adaptations to therapy. Assays applied to each biopsy include bulk genomics (tDNA), transcriptomics (RNA), and proteomics (RPPA) as well as single-cell epigenomics (sciATAC) and multiplexed tissue imaging (mIHC, cycIF) for single-cell spatial proteomics. b) Patient treatment regimen and biopsy timelines. Timelines began when the pre-treatment biopsy was taken (labeled as day 0). Horizontal bars represent the duration of treatment with CDK4/6i, an estrogen receptor inhibitor (ERi), and other relevant therapies. Several biopsy pairs were collected from the same metastatic patient because the patient was treated with different CDK4/6i therapies. In total, 7 biopsy pairs were collected from 5 patients. Only therapies given between paired pre-treatment and on-progression biopsies are shown in the figure. Additional patient treatment regimens are available through the HTAN Data Portal (https://data.humantumoratlas.org/). c) Summary of assays completed on each biopsy. Clinical metadata includes CDK4/6i and ERi used for therapy, patient response, PAM50 score, biopsy site, and HTAN patient ID.

Duration of CDK4/6i treatment ranged from 1.5 months to 3.5 years (**Figure 1b**). Four of five patients responded to therapy before developing acquired resistance. The fifth patient, 9-2, showed no clinical response and therapy was stopped after 1.5 months (**Figure 1b, 1c**). Two patients, 9-1 and 9-14, received the CDK4/6i palbociclib (9-1P, 9-14P) as frontline therapy and then in later lines of therapy received abemaciclib monotherapy (9-1A, 9-14A) per its approved indication. Patient 9-1 received abemaciclib 1,129 days after palbociclib was stopped, and abemaciclib was taken for 348 days before resistance developed. Patient 9-14 received abemaciclib 309 days after palbociclib was stopped, and abemaciclib was taken for 238 days until the tumor became refractory. Additional pre- and on-progression biopsies were acquired for those patients, with a total of thirteen biopsies acquired from all five patients (**Figure 1b**).

Tumor tissue was collected from various biopsy sites, including seven liver, three bone, and three lymph node biopsies. Six complementary molecular/proteomic assays were used to generate data for each biopsy (**Figure 1c**). These included high depth targeted DNA sequencing (tDNA-seq), bulk transcriptome sequencing (RNA-seq), reverse phase protein array (RPPA) (Tibes et al. 2006), single-cell combinatorial indexing assay for transposase-accessible chromatin sequencing (sciATAC-seq) (Mulqueen et al. 2021), cyclic immunofluorescence (cycIF) (J.-R. Lin et al. 2018; Eng et al. 2020), and multiplex immunohistochemistry (mIHC) (Banik et al. 2020). The methods section provides details for each assay used in this study.

### Malignant cell adaptations to CDK4/6i therapy

In breast cancer, malignant epithelial cells take advantage of changes in their genomic, transcriptional, and proteomic activity to promote proliferation, differentiation, and survival. Analysis of the molecular datasets in our atlas suggests several malignant cell adaptations to CDK4/6i therapy across multiple modalities.

Genomic alterations identified within the cohort are consistent with previous reports observed before and after CDK4/6i treatment (Wander et al. 2020). Prior to therapy, the most common genomic alterations in our metastatic cohort were loss of cyclin-dependent kinase inhibitor 2A (*CDKN2A*), amplification of cyclin D1 (*CCND1*), activating mutations in estrogen receptor 1 (*ESR1*), and loss of cyclin-dependent kinase inhibitor 1A (*CDKN1A*) (**Figure 2a**). Only three of five patients in our cohort were found to have genomic alterations previously associated with resistance to CDK4/6i (Wander et al. 2020; Cristofanilli et al. 2018; Park et al. 2023), and no patients in our cohort shared genomic alterations associated with resistance. Resistance-associated genomic alterations observed in our cohort include acquired aberrations in *RB1* (9-15 Bx2), *ESR1* (9-15 Bx2), and *PIK3CA* (9-1 Bx2). A single copy *RB1* loss was present in the pre-treatment biopsy for the non-responding patient 9-2, though typically only biallelic *RB1* loss-of-function alterations have been associated with intrinsic CDK4/6i resistance (Herrera-Abreu et al. 2016; Condorelli et al. 2018; O’Leary et al. 2018; Wander et al. 2020) (**Figure 1b, 1c, 2a; Extended Data** Figure 1). Some tumors in our cohort (9-3, 9-14) did not acquire any notable genomic alterations following therapy. Given the limited number of genomic alterations present in our cohort, the use of additional assays is essential to fully reveal mechanisms of therapeutic resistance.

**Figure 2:**
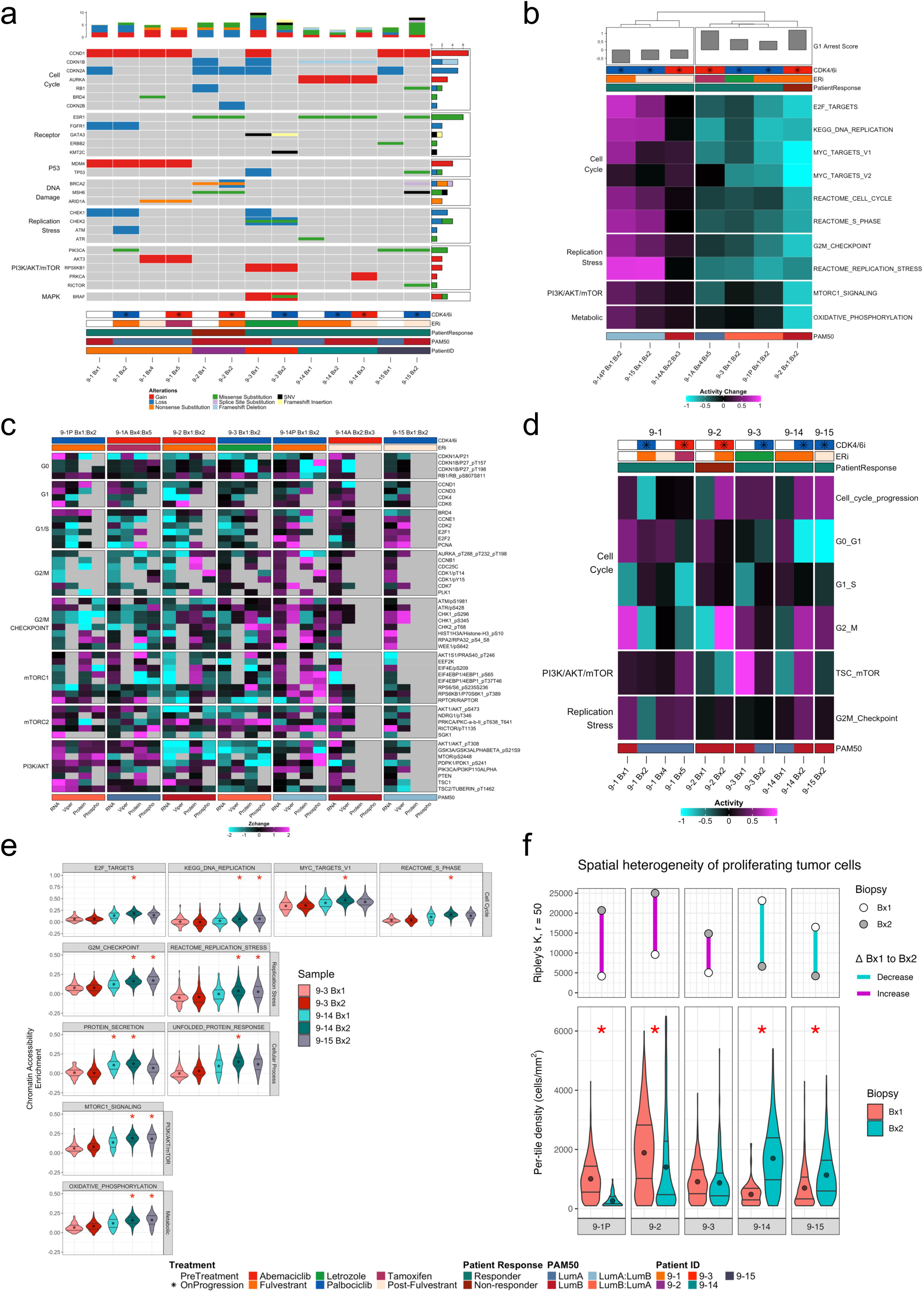
Multimodal profiling reveals malignant cell adaptations to CDK4/6i. **a)** Oncoplot of genomic alterations from high depth targeted DNA-seq. Oncoplot includes alterations from 24 genes that have been previously associated with resistance to CDK4/6i and ET and altered in at least one patient biopsy using a targeted sequencing panel of around 200 genes. Genes (rows) are grouped by pathway and function and ordered by frequency. A comprehensive set of all genomic alterations found across patient samples can be seen in Extended Data Figure 1. **b)** Transcriptional gene set variation analysis (GSVA) of malignant cell pathways during CDK4/6i therapy and progression. Heatmap describes the change in pathway enrichment during each line of CDK4/6i therapy across HTAN cases. Columns are annotated with transcriptional G1 arrest scores representing the extent to which CDK4/6i prevents transcription for cell cycle progression. Pathways (rows) in which the average change in activity between G1-positive and G1-negative tumors is greater than 0.5 were selected for the plot. Biopsy pairs (columns) are hierarchically clustered to show similar patterns of activity. **c)** Integrated heatmap of malignant cell RNA and protein changes during CDK4/6i therapy and progression. Heatmap describes the relative change of malignant cell markers during therapy using four measurements of regulation: RNA expression (RNA), regulator activity (Viper), protein abundance (Protein), and phosphoprotein (Phospho) levels. Genes and proteins included in the heatmap were selected and grouped based on GSVA results from Figure 2b to highlight specific malignant cell markers of regulation and activation across bulk assays. Each value is computed by first scaling and mean centering the data relative to a background HR+ cohort and for each pair subtracting the pre-treatment biopsy from the on-progression biopsy. **d**) Malignant cell proteomic pathway signaling during CDK4/6i therapy and progression. Heatmap describes the change in protein pathway scores during each line of CDK4/6i therapy across HTAN cases. Pathways included in the heatmap were selected and grouped based on GSVA results from Figure 2b to show concordance across RNA and protein assays. **e)** Violin plots showing distributions of tumor cell chromatin accessibility enrichment scores for malignant cell pathways from sciATAC-seq. Pathways were selected based on GSVA results from Figure 2b to show concordance in transcriptional and epigenetic regulation across assays. For all violin plots, the mean value is shown as a point, and the first (25%) and third quartiles (75%) are represented as vertical lines within each distribution. Red asterisks represent statistically significant samples when compared to all other samples (FDR < 0.01, one-sided Mann-Whitney test). **f)** Quantification of spatial heterogeneity of proliferating tumor cells during CDK4/6i therapy and progression using cycIF. Tumor cells were defined as positive for at least one cytokeratin marker, and were considered proliferative if positive for Ki67, PCNA, or pHH3. Top: Degree of clustering, calculated as univariate Ripley’s K, for proliferative tumor cells in each biopsy within a neighborhood radius of 50 micrometers. Increasing degree of clustering suggests higher co-localization of proliferative tumor cells relative to a sample-specific permuted random point process. Ripley’s K values across the cohort were all positive, suggesting that proliferative tumor cells cluster together significantly when compared to a random point process; however, the magnitude of the degree of clustering varied widely between 4172 and 24998 across the cohort. Bottom: Violin plots showing the distribution of proliferative tumor cell density across 0.01mm2 tiles within each biopsy. Mean tile densities of proliferating tumor cells varied across the cohort, with a minimum mean density of 145 cells/mm2, and a maximum of 1646 cells/mm2. For all violin plots, the mean value is shown as a point, and the first (25%) and third quartiles (75%) are represented as vertical lines within each distribution. Red asterisks represent statistical significance between patient biopsy pairs (p < 0.01, two-sided Mann-Whitney test).

Given CDK4/6 inhibitors are cytostatic therapies, we investigated the effect of therapy on tumor proliferation and cell cycle activity using a full suite of multimodal assays (**Figure 1c**) and observed a distinct split within our cohort. We hypothesized tumors where the CDK4/6i inhibitor is successfully controlling proliferative growth would stall at the G1-S checkpoint and display an inability to progress through the cell cycle. Using G1 arrest score, a transcriptional metric to quantify stalling at the G1-S checkpoint (Hafner et al. 2019), we observed three tumors (9-1, 9-2, and 9-3) had an increased score consistent with stalling in G1 phase. In contrast, two tumors (9-14 and 9-15) had decreased G1 arrest score suggesting progression through the G1-S checkpoint despite CDK4/6 inhibition. In the tumors with decreased G1 arrest scores, transcriptional activity in pathways related to cell cycle, mTORC1 signaling, and replication stress was highly elevated following progression (**Figure 2b**). It is not possible to clearly separate the effects of endocrine therapy and CDK4/6i for these patients because the therapies were given in combination. However, there were no substantial changes in clinical IHC receptor levels (**Supplemental Table 1)** or transcriptional hormone signaling post-therapy (**Extended Data** Figure 2) compared to pre-therapy. These data suggest the lack of proliferative control for some tumors may relate to hormone-independent adaptations to CDK4/6.

To contextualize and validate the multiomic observations from our cohort, we analyzed an external cohort of 23 metastatic HR+ breast cancers with paired bulk RNA-seq samples treated with CDK4/6i(Park et al. 2023). Like our cohort, two groups with distinct transcriptional changes were observed. In the external cohort, 13 of 23 tumor pairs showed decreased G1 arrest scores accompanied with significant activation of cell cycle progression, mTORC1, and replication stress upon progression with CDK4/6i (FDR < 0.05; two-sided Mann-Whitney test) (**Extended Data** Figure 3**; Supplemental Table 2**). Based on the observation that tumors displayed either minimal cell cycle activity or markedly increased cell activity after therapy, we performed an integrative analysis that combined our atlas with the external cohort. Hierarchical clustering of the joint cohort revealed tumors clustered with either increasing (termed Malignant Cell “MC-high”) or decreasing (termed Malignant Cell “MC-low”) proliferative and cell cycle pathway activity during CDK4/6i therapy and progression (**Extended Data** Figure 4). Taken together, these transcriptional observations suggest some tumors maintain control of proliferative growth despite progression on therapy whereas other tumors may circumvent the cytostatic effects of CDK4/6i, leading to progression.

Further investigation of the tumors from 9-14 and 9-15 revealed multiple adaptations potentially associated with increased proliferation and consequently therapeutic resistance. Gene and protein activity of *CDK2* and *E2F1* sharply increased between the pre-treatment and on-progression biopsies, indicating continued proliferation despite CDK4/6 inhibition. In 9-14P—the pair of biopsies taken before and after palbociclib treatment—this increase was accompanied by elevated *CCND1* activity, cyclin D3 protein abundance, and p27 phosphorylation (T157), whereas 9-15 had increased *CCNE1* activity (**Figure 2c; Extended Data** Figure 5). Taken together and consistent with other studies (Herrera-Abreu et al. 2016; Al-Qasem et al. 2022), observations identify tumors that bypass cyclin D-CDK4/6 signaling to progress through the G1-S checkpoint using non-canonical induction of cyclin D via CDK2 or hyperactivation of the cyclin E-CDK2 complex, to promote cell cycle progression.

Concordant with the transcriptomic results, proteomic pathway activity (RPPA) revealed an increase in proliferation for 9-14P, and high levels of cell cycle-related protein abundance for 9-15 (**Figure 2d**). Chromatin accessibility (sciATAC-seq) identified cell cycle pathway modules that were significantly elevated in malignant epithelial cells of the highly proliferative tumors (FDR < 0.01, one-sided Mann-Whitney test) (**Figure 2e; Extended Data** Figures 6 and 7). Spatial single-cell cyclic immunofluorescence (cycIF) revealed two biopsy pairs with negative G1 arrest scores (9-14P, 9-15) that contained significantly increased density of proliferative malignant epithelial cells and decreased spatial clustering after therapy (p < 0.01, two-sided Mann-Whitney test) (**Figure 2f**). In contrast, tumors 9-1P, 9-2, and 9-3 were decreased in proliferative density and increased in spatial clustering of proliferative malignant cells.

In tumors from 9-14 and 9-15, high proliferative activity was accompanied by increased activity of PI3K/AKT/MTOR, a pathway frequently dysregulated in HR+ breast cancers and associated with CDK4/6i resistance (Yoshida et al. 2019; Knudsen et al. 2019; Occhipinti et al. 2020; Michaloglou et al. 2018; Herrera-Abreu et al. 2016). Activation of mTORC1 and its downstream substrates results in sustained cellular growth, delayed apoptosis, and continued cellular proliferation, including induction of cyclin D expression to promote cell cycle entry (Zhilin Zou et al. 2020; Ciołczyk-Wierzbicka et al. 2020). This may partially explain acquired resistance in 9-14P in which mTORC1 activation coincided with an increase in *CCND1* expression and cyclin D3 protein abundance (**Figure 2c**). The mTORC1 pathway was especially elevated in 9-14P across all modalities, displaying increased phosphorylation of mTORC1 substrates (pS6 S235/S236; p4EBP1 S65) and enrichment of both chromatin accessibility and transcription of mTORC1 signaling (**Figure 2b, 2d, 2e**).

Lastly, replication stress was observed in the highly proliferative tumors from 9-14 and 9-15, potentially as a result of genotoxic effects induced by aberrant cell cycle entry or prolonged cell cycle arrest (Fallah et al. 2021; Crozier et al. 2022). Both 9-14 and 9-15 had increased transcriptional and chromatin accessibility enrichment of the pathways REACTOME_ACTIVATION_OF_ATR_IN_RESPONSE_TO_REPLICATION_STRESS (Jassal et al. 2020) and G2M_CHECKPOINT (Liberzon et al. 2015). Activation of these pathways was accompanied by elevated Replication Protein A 32 (RPA32) phosphorylation, an indicator of replication stress (**Figures 2b, 2c, 2e**). The role of replication stress as a mechanism of resistance is less established but hypothesized to allow cells with impaired G1/S checkpoint to escape cell cycle arrest by relying on the G2/M checkpoint machinery to surveil and repair DNA (Fallah et al. 2021; Scheidemann and Shajahan-Haq 2021).

In summary, multiple but not all tumors in this cohort displayed increased cell cycle activity and proliferation after progressing on CDK4/6i therapy that could not be accounted for by acquired genomic alterations. Cell cycle and proliferative activity were concordant across bulk RNA-seq, single cell and bulk proteomics, and scATAC-seq, indicating that progression on CDK4/6i may be due to direct resistance to the cytostatic effects of CDK4/6i. For highly proliferative tumors, upregulation of cell cycle genes including *CDK2*, *CCNE1*, and *CCND1* may bypass the CDK4/6 complex to alleviate G1 stalling and promote cell cycle progression. Hyperactivation of the PI3K/AKT/mTOR pathway was observed to be a potential mechanism of resistance. Replication stress was observed as a byproduct of tumor proliferation or as a mechanism to circumvent G1-S stalling.

### Immune Responses on CDK4/6i Therapy

Previous studies have reported CDK4/6i stimulation of immunoregulatory pathways that promote anti-tumor immunity by recruiting cytotoxic T cells following antigen presentation, increasing cytokine and interferon signaling, and suppressing regulatory T cells (Spranger 2016; Goel et al. 2017; Scirocchi et al. 2022; Zhu et al. 2022). Longer PFS in HR+ breast cancer patients treated with combination palbociclib and letrozole has also been associated with expression of immunomodulatory genes (Zhu et al. 2022). These findings motivated us to quantify leukocyte contexture induced by CDK4/6i therapy within this metastatic HR+ breast cancer cohort. We used bulk RNA-seq and RPPA to quantify global transcriptional and proteomic changes and multiplexed immunohistochemistry (mIHC) (Banik et al. 2020) for spatial single-cell leukocyte profiling revealing both T cell activation and suppression were observed in most tumors. This section discusses observations supporting immune activation within the tumor-immune microenvironment (TIME) in our atlas, and the next section discusses observations associated with immune suppression.

To characterize the transcriptional changes within the TIME associated with CDK4/6i therapy in metastatic HR+ breast cancer, we used the same approach from the previous section. A joint cohort of 30 CDK4/6i-treated breast cancers was created by combining our seven pre-and post-CDK4/6i RNA-seq datasets with the same external RNA-seq dataset of 23 CDK4/6i treated metastatic breast cancers(Park et al. 2023), and this joint cohort was used for transcriptional analyses. To quantify potential tissue-specific effects from different metastatic tumor sites, we first evaluated immune-related transcriptional signatures between tissue types and found only minor differences in activity (**Extended Data** Figure 8a**, 8b**). Next, hierarchical clustering was performed on the 30 total RNA-seq pairs from the joint cohort using activity levels of immune-related gene sets from the MSigDB hallmarks collection (Liberzon et al. 2015). Clustering revealed two distinct subgroups of patients having either increasing (“TIME-high”) or decreasing (“TIME-low”) activity of immune-related pathways including INTERFERON_ALPHA_RESPONSE, INTERFERON_GAMMA_RESPONSE, and IL6_JAK_STAT3_SIGNALING (**Extended Data** Figure 4). Comparison of these TIME-delineated clusters to the malignant cell (MC) pathway activity clusters described in the previous section showed no clear associations. Some tumors exhibited both MC-high and TIME-high activity or MC-low and TIME-low activity, but other tumors showed activation of either malignant cell or immune-related pathways but not both. Several tumors in our atlas had high activity in one set of pathways but not in the other, including 9-14P (MC-high/TIME-low) and 9-1P, 9-1A, and 9-3 (MC-low/TIME-high) (**Extended Data** Figure 4).

Five tumors from our cohort (9-1P, 9-1A, 9-3, 9-14A, and 9-15) exhibited TIME-high activity. These tumors displayed an increase in T cell inflamed GEP score, a transcriptional prognostic and putative immunotherapy biomarker used to assess the degree of T cell inflammation within the TIME(Ayers et al. 2017) as well as increased enrichment of multiple immune cell signatures(Tamborero et al. 2018) (**Figure 3a; Extended Data** Figures 4**, 9a, and 9b**). Bulk proteomic activity from RPPA were consistent with RNA-seq: three biopsy pairs (9-1P, 9-1A, 9-15) contained elevated activity of the JAK/STAT proteomic signature along with elevated abundance of multiple JAK/STAT proteins and phospho-proteins (**Figure 3b; Extended Data** Figure 10**; Supplemental Figure S1**). The association between INTERFERON_ALPHA_RESPONSE, INTERFERON_GAMMA_RESPONSE, and IL6_JAK_STAT3_SIGNALING was expected as interferons activate the JAK/STAT pathway (Villarino et al. 2015). The two tumors in the TIME-low cluster, including the non-responder tumor 9-2, displayed neither transcriptional nor proteomic immune activation after therapy.

**Figure 3:**
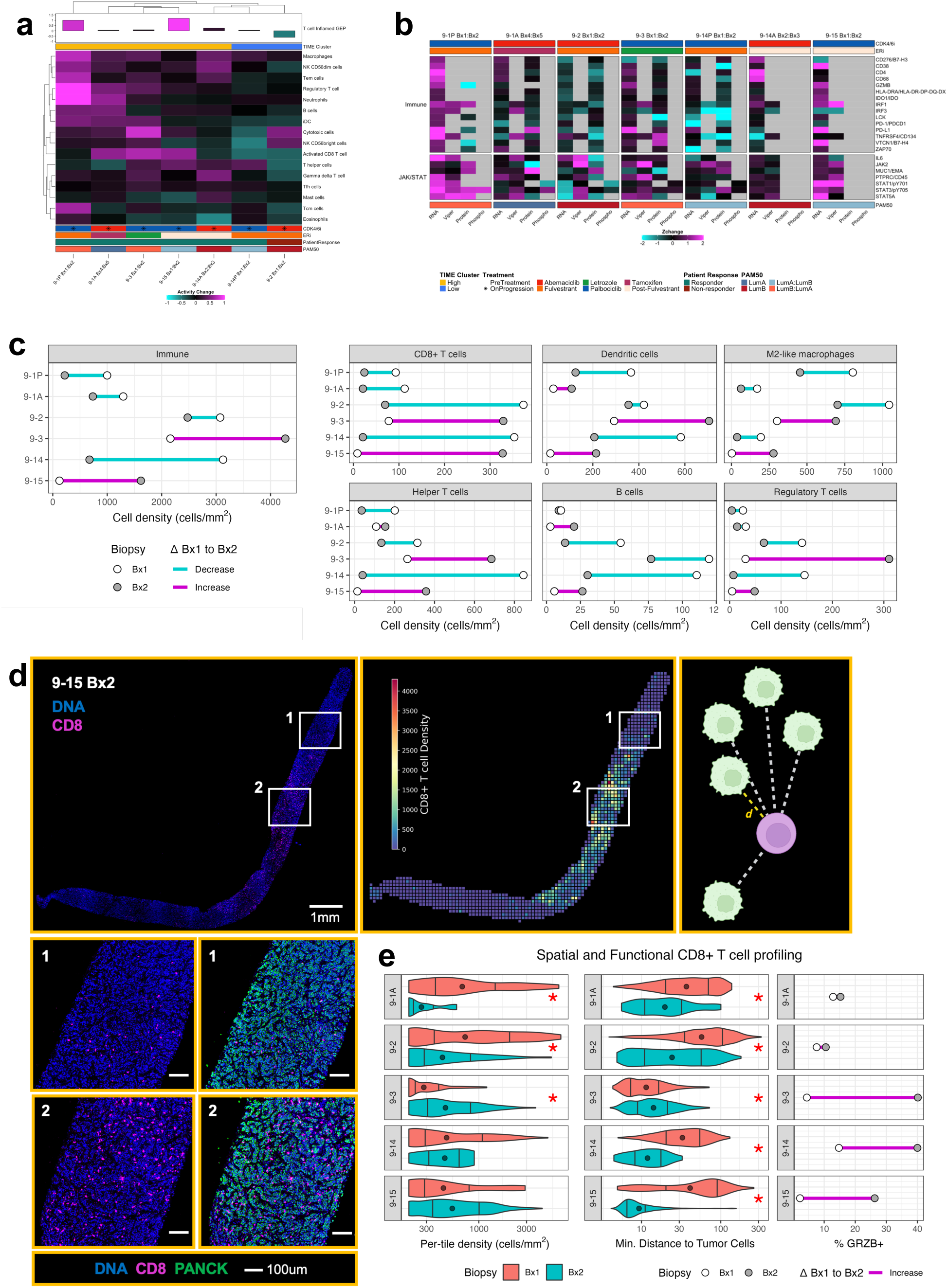
Multi-omics and mIHC reveal heterogeneous changes to the tumor-immune microenvironment suggestive of mechanisms of therapeutic resistance. **a)** Transcriptional activity of immune cell types during CDK4/6i therapy and progression. Heatmap describes the change in activity for sixteen immune cell types quantified with GSVA. Biopsy pairs (columns) are hierarchically clustered to show similar patterns of activity and annotated with change in T cell inflamed GEP score before and after therapy. **b)** Integrated heatmap of TIME RNA and protein changes during CDK4/6i therapy and progression. Heatmap describes the relative change of TIME markers during therapy using four measurements of regulation: RNA expression (RNA), regulator activity (VIPER), protein abundance (Protein), and phosphoprotein (Phospho) levels. Genes and proteins included in the heatmap were selected and grouped based on GSVA results from Figure 3a to highlight TIME markers of immune regulation and signaling across bulk assays. Each value is computed by first scaling and mean centering the data relative to a background HR+ cohort and for each pair subtracting the pre-treatment biopsy from the on-progression biopsy. **c)** Cell densities reported for all CD45^+^ immune cells and individual immune cell type populations across pre-treatment (Bx1, white) and on-progression (Bx2, dark grey) biopsies for all biopsy pairs using mIHC. Line color connecting the pre-treatment and on-progression biopsy points represents direction of change from Bx1 to Bx2. Across all biopsies, the density of all CD45^+^ immune cells varied between 122 cells/mm^2^ and 4270 cells/mm^2^. The individual immune lineages varied in density across the cohort, ranging between 0 cells/mm^2^ and 1050 cells/mm^2^. **d)** Psuedocolored mIHC whole slide image and select regions of interest of 9-15 Bx2 showing spatial heterogeneity of CD8 marker (magenta). For both the whole slide image and the regions of interest, the DNA channel is colored blue. The rightmost column of the regions of interest shows the pan-cytokeratin (PANCK) marker in green. On the right, analytical methods for quantifying spatial organization and heterogeneity of cell types using mIHC, including the utilization of a fishnet grid to divide the biopsy into 0.01mm2 tiles. Each tile is colored by the density of CD8^+^ T cells within the tile. Secondly, minimum distances are calculated between a target cell type (e.g., tumor cells) and a query cell type (e.g., CD8^+^ T cells) for every query cell in the sample. e) Spatial heterogeneity and cytotoxic function of CD8^+^ T cells during CDK4/6i therapy and progression. Left: Violin plots showing the distribution of CD8^+^ T cell density across 0.01mm^2^ tiles from each biopsy. Mean CD8^+^ T cell tile densities ranged between 293 cells/mm^2^ and 1348 cells/mm^2^ across the cohort. Middle: Violin plot showing the per-cell distribution of minimum distances to tumor cells for all CD8^+^ T cells in each biopsy. Across all biopsies in the cohort, minimum distances between CD8^+^ T cells and tumor cells varied between 12 μm and 76 μm. Right: Bar plots showing the percentage of CD8^+^ T cells positive for Granzyme B. The mean percentage of Granzyme B^+^ CD8^+^ T cells across the cohort was 17.4%, with a range between 2.2% and 40.13%. For all violin plots, the mean value is shown as a point, and the first (25%) and third quartiles (75%) are represented as vertical lines within each distribution. Red asterisks represent statistical significance between patient biopsy pairs (p < 0.01, two-sided Mann-Whitney test).

While global immune signaling activation in RNA-seq supported an increased immune response, the immune system’s ability to clear tumor is highly dependent on leukocyte composition and functional status within the TIME (Trujillo et al. 2018; Munhoz and Postow 2016). We used mIHC, a spatial single-cell proteomics profiling assay, to survey the types, functional status, and spatial organization of leukocytes in the TIME during CDK4/6i therapy. The four biopsy pairs in the TIME-high cluster displayed distinct changes in leukocyte infiltration after therapy: tumors 9-3 and 9-15 contained higher overall CD45^+^ cell density, whereas 9-1P and 9-1A had decreased CD45^+^ cell density after therapy (**Figure 3c**). In tumors 9-3 and 9-15, there was a large increase in density of CD45^+^ cell density encompassing lineages frequently associated with anti-tumoral effects, including CD8^+^ T cells, helper T cells, dendritic cells, and B cells (Kravtsov et al. 2022; Marciscano and Anandasabapathy 2021; Sautès-Fridman et al. 2019) (**Figure 3c; Supplemental Table 3**). 9-1A also exhibited increased density of helper T cells, B cells, and dendritic cells but these changes were minimal. Tumors from 9-3 and 9-15 contained increased helper T cell density after therapy, with helper T cells comprising over 15% of the CD45^+^ population in the 9-3 and 9-15 on-progression biopsies (**Figure 3c; Supplemental Figure S2**). Changes in dendritic cell abundance were highly heterogenous across biopsy pairs. Dendric cells play a critical role in antigen presentation and T cell recruitment (Marciscano and Anandasabapathy 2021); these increased in density for 9-3 and 9-15 (**Figure 3c**). Bulk RNA-seq results revealed increased enrichment of antigen presentation and chemokine pathways for the tumors in the TIME-high cluster, particularly for 9-15 (**Extended Data** Figure 2 and 4) (Wang et al. 2019; Prabhakaran et al. 2017). In 9-2 and 9-14, there was consistency across assays where RNA-seq and RPPA revealed decreased immune activity after treatment, and mIHC showed decreased CD45^+^ cell infiltration and densities of immunoreactive cell populations (**Figure 3b, 3c**).

Given the important role of CD8^+^ T cells in anti-tumor immunity, we next investigated their cytotoxic state and spatial organization. CD8^+^ T cells induce apoptosis by secreting granzyme B and perforin (Raskov et al. 2021). Growing evidence has also associated the spatial organization of CD8^+^ T cells with therapeutic response and survival (Fu et al. 2021). In our cohort, CD8^+^ T cell populations displayed overall increase in cytotoxicity as evident by the increase in granzyme B (GRZB)-positive cells and Cytotoxic cells signature, suggestive of an anti-tumor immune response elicited by CDK4/6i (**Figure 3a**). Despite this overall increase in cytotoxicity, the TIME exhibited heterogeneous changes in CD8^+^ T cell density and spatial proximity to malignant cells on progression. Concordant with the Activated CD8^+^ T cell pathway (Tamborero et al. 2018) activity from RNA-seq, 9-3 and 9-15 contained increased CD8^+^ cell density and relative proportion (**Figure 3c; Supplemental Figure S2**). To assess the spatial heterogeneity of leukocyte composition within a biopsy, we divided biopsy tissue images into tiles (0.01mm^2^; average 118 cells/tile). The 9-15 on-progression biopsy contained abundant CD8^+^ T cell infiltration that was highly co-localized with malignant tumor cells (**Figure 3d, 3e**). 9-3 exhibited an increase in overall CD8^+^ T cell density and maintained a high degree of co-localization between CD8^+^ T cells and tumor cells before and after therapy (**Figure 3e**). Although CD8^+^ T cells co-localized more frequently with tumor cells in 9-1A, 9-2, and 9-14, overall CD8^+^ T cell density decreased on-progression, indicating reduced infiltration (or retention) overall. The 9-2 pre-treatment biopsy, deemed a clinical non-responder to CDK4/6i, had a high degree of spatial exclusion of CD8^+^ T cells distal to malignant tumor cells, indicative of potential physical barriers arising from stroma or extracellular matrix, preventing further infiltration (**Figures 3e**).

In summary, while four biopsy pairs showed immune activation in bulk transcriptomics and proteomics, only tumors 9-3 and 9-15 had sustained active anti-tumor immune responses based on single-cell spatial proteomics. Single-cell spatial proteomics revealed that these two tumors had substantial increases in abundance of anti-tumor immune cell populations during therapy with a shift in cytotoxic cell spatial organization toward an anti-tumor state. Two biopsy pairs from patient 9-1 on different CDK4/6i therapies exhibited increased global immune signaling in RNA-seq and RPPA data. However, single-cell spatial proteomics revealed decreased (9-1P) or only minor increases (9-1A) in the abundance of immunoreactive cell types. 9-2 and 9-14 showed a reduction in immune activation consistent across bulk and single-cell spatial assays, and the pre-treatment biopsy for 9-2 exhibited TIME organization hypothesized to reduce cytotoxic cell efficacy.

### T cell Suppressive Changes Observed on CDK4/6i Therapy

Many biopsies in the cohort—including those with strong evidence of immune activation—displayed evidence of immunosuppression. Tumors evade immune surveillance through multiple mechanisms to prevent T cell infiltration and promote immunomodulatory processes that reduce their activation. In T cell-inflamed environments, chronic inflammation can develop from prolonged cytokine signaling favoring tumor immunosuppression and progression (Zaidi 2019; X. Zhang et al. 2017; Qadir et al. 2017). Further, recent studies have linked interferon and JAK/STAT signaling with CDK4/6i resistance (Kettner et al. 2019; De Angelis et al. 2021). We used RNA-seq, RPPA, and mIHC to explore multiple aspects of immune suppression, including macrophage polarization, the PD-1/PD-L1 axis, regulatory T cell recruitment, and dysfunction/exhaustion of effector T cells.

Anti-inflammatory signaling caused by prolonged interferon signaling can shift myelomonocytic cell (monotypes and macrophages) polarization to protumoral states (Poh and Ernst 2018; Boutilier and Elsawa 2021; Goswami, Bose, and Baral 2021; Zijuan Zou et al. 2023) resulting in tumor progression (Larionova et al. 2020; Noy and Pollard 2014; E. Y. Lin et al. 2006; Tu et al. 2021). High intra-tumoral abundance of myelomonocytic cells represents a poor prognostic indicator of overall survival and progression-free survival in breast cancer (Larionova et al. 2020; Tiainen et al. 2015; Ni et al. 2019; Medrek et al. 2012; Esbona et al. 2018). Using mIHC, we quantified the changes of protumoral CD163^+^ macrophages in six biopsy pairs from our cohort. Two of six biopsy pairs (9-3 and 9-15) exhibited a substantial increase in CD163^+^ macrophage density after therapy, while all other biopsy pairs contained decreased CD163^+^ macrophage density (**Figure 3c, 4a**). CD163^+^ macrophages resided in closer proximity to malignant cells in three biopsy pairs (9-2, 9-14, 9-15) after therapy. The 9-2 pre-treatment biopsy, which was resistant to therapy, displayed striking TIME spatial organization (**Figure 4b**). While CD8^+^ T cells were sequestered away from malignant cells in the 9-2 pre-treatment biopsy, CD163^+^ macrophages were concentrated both in stromal regions proximal to malignant cells and infiltrated throughout malignant-cell dense regions as highlighted by the bimodal distance distribution (**Figure 4a, 4b**).

**Figure 4:**
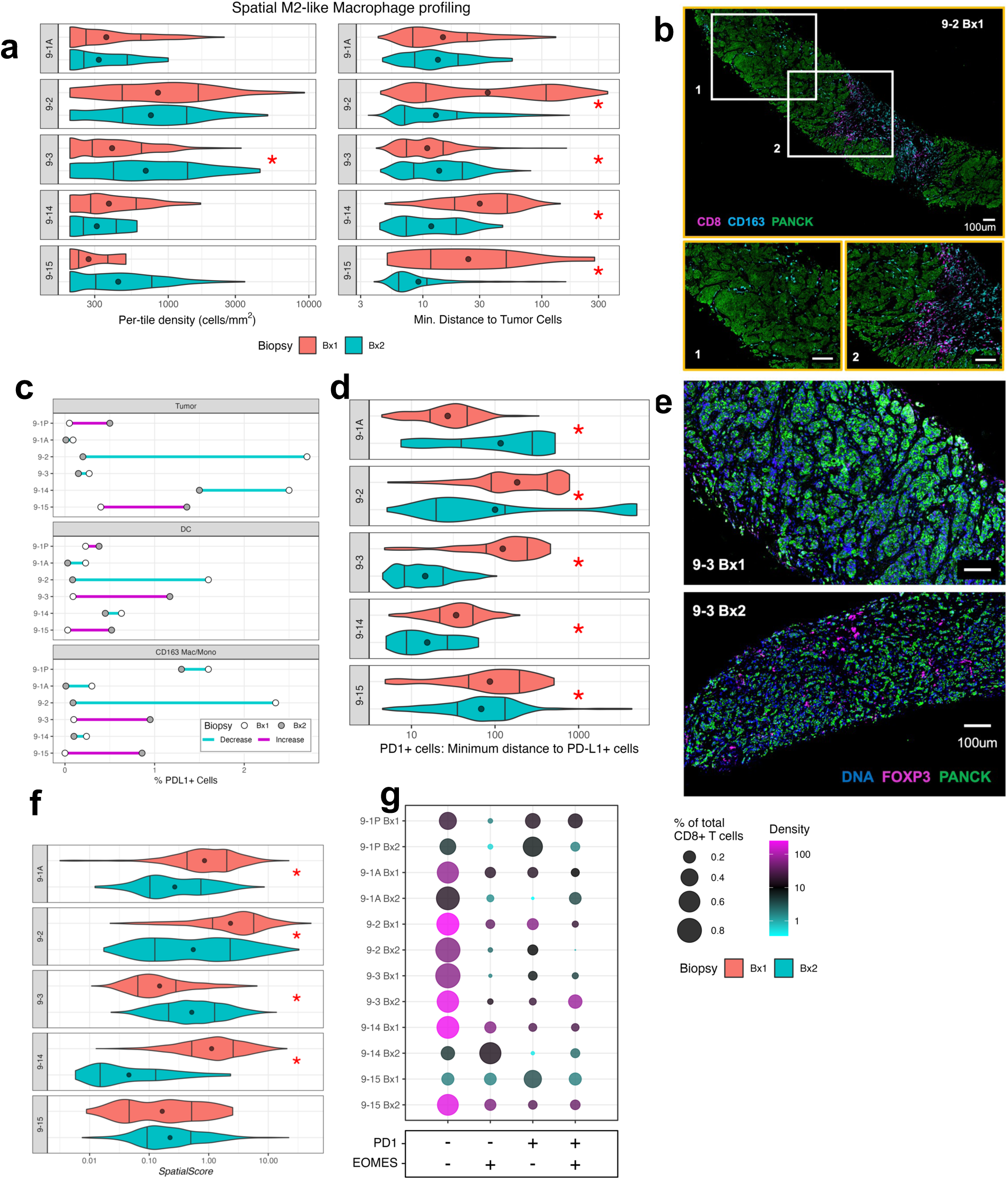
Immune suppression changes observed on CDK4/6i therapy. **a)** Spatial heterogeneity of CD163^+^ macrophages/monocytes during CDK4/6i therapy and progression. Left: Violin plots showing the distribution of CD163^+^ macrophages/monocytes density across 0.01mm^2^ tiles from each biopsy. Mean tile densities of CD163^+^ macrophages varied between 300 cells/mm^2^ and 1226 cells/mm^2^ across the cohort. Middle: Violin plot showing the per-cell distribution of minimum distances to tumor cells for all CD163^+^ macrophages/monocytes in each biopsy. Mean minimum distances between CD163^+^ macrophages and tumor cells varied between 12 μm and 69 μm. For all violin plots, the mean value is shown as a point, and the first (25%) and third quartiles (75%) are represented as vertical lines within each distribution. Red asterisks represent statistical significance between patient biopsy pairs (p < 0.01, two-sided Mann-Whitney test). **b)** Psuedocolored mIHC whole slide image and select regions of interest of 9-2 Bx1 showing spatial heterogeneity of CD8 marker (magenta) and CD163 marker (cyan). For both the whole slide image and the regions of interest, the PANCK channel is colored green. All scale bars for both the whole slide image and regions of interest are 100 μm. **c)** Percent of cells that are positive for PD-L1 reported for tumor cells, dendritic cells (DC), and CD163^+^ macrophages/monocytes across pre-treatment (Bx1, white) and on-progression (Bx2, dark grey) biopsies for all biopsy pairs using mIHC. Line color connecting the pre-treatment and on-progression biopsy points represents direction of change from Bx1 to Bx2. Across tumor cells and antigen presenting immune cell lineages, the percentage of PDL1^+^ cells varied between 0-3%. **d)** Minimum distances from PD-1^+^ cells to PD-L1^+^ cells during CDK4/6i therapy and progression reported using mIHC. For all biopsies across the cohort, the mean minimum distance between PD-1^+^ and PD-L1^+^ cells varied between 18 μm and 962 μm. For all violin plots, the mean value is shown as a point, and the first (25%) and third quartiles (75%) are represented as vertical lines within each distribution. Red asterisks represent statistical significance between patient biopsy pairs (p < 0.01, two-sided Mann-Whitney test). **e)** Psuedocolored mIHC images of select regions of interest from Bx1 (top) and Bx2 (bottom) of patient 9-3 showing abundance of FOXP3 (magenta), a marker for Regulatory T cells. For both images, the DNA channel is colored blue, and PANCK is green. Scale bars are 100 micrometers. **f)** SpatialScore during CDK4/6i therapy and progression. The SpatialScore is the ratio of the minimum distance between a helper T cell and a tumor cell to the minimum distance between a helper T cell and a Regulatory T cell, reported for all helper T cells in each biopsy. Larger SpatialScore values indicate more frequent co-location of helper T cells and Regulatory T cells. Across all biopsies, the mean SpatialScore across all helper T cells varied between 0.16 and 4.5. For all violin plots, the mean value is shown as a point, and the first (25%) and third quartiles (75%) are represented as vertical lines within each distribution. Red asterisks represent statistical significance between patient biopsy pairs (p < 0.01, two-sided Mann-Whitney test). **g)** Differentiation states of CD8^+^ T cells based on PD-1 and EOMES markers using mIHC. CD8^+^ T cells were classified as positive or negative for both PD-1 and EOMES and divided into groups based on the combination of these markers. Double negative (PD-1^-^/EOMES^-^) are representative of naïve CD8^+^ T cells. Double positive (PD-1^+^/EOMES^+^) are suggestive of CD8^+^ T cell dysfunction and exhaustion. PD-1^+^/EOMES^-^ and PD-1^-^/EOMES^+^ are representative of early and late effector CD8^+^ T cells respectively. Bubble color shows the density of each T cell subset within the biopsy, and bubble size shows the relative abundance of each subset within the CD8^+^ T cell population.

The PD-1/PD-L1 checkpoint axis is a key component of lymphocyte regulation and a strong indicator of immunosuppression in cancer where PD-L1 presence is negatively associated with PFS in HR+ breast cancers on CDK4/6i therapy (Zhu et al. 2022). PD-L1 is often expressed by CD163^+^ macrophages closely associated with malignant cells either via direct PD-L1 expression and by modulating malignant cell expression of PD-L1 (Pu and Ji 2022). Both 9-3 and 9-15 had sharp increases in bulk RNA-seq REACTOME_PD_1_SIGNALING pathway activity with moderate increases for 9-1P and 9-1A after therapy. Two pre-treatment biopsies (9-2 and 9-14) also displayed elevated activity for the same PD-1 Signaling pathway relative to the cohort, indicating the TIME for these tumors was in a T cell suppressive state prior to therapy (**Extended data Figure 2**) (Jassal et al. 2020). PD-L1 protein abundance in mIHC was low in our cohort at 0-3% of cells, which is expected for HR+/HER2-breast cancer (Núñez Abad et al. 2022). However, small increases in cells expressing PD-L1 post therapy were observed across malignant cells, dendritic cells, and macrophages for 9-3 and 9-15 (**Figure 4c**). All biopsy pairs, except 9-1A, showed a decrease in the mean minimum distance between PD-1^+^ and PD-L1^+^ cells post therapy (**Figure 4d**). Reduced distance between PD-1^+^/PD-L1^+^ cells allow for direct receptor-ligand interactions required to suppress T cell functional state.

Recruitment of regulatory T cells (FoxP3^+^), which function to suppress cytotoxic and helper T cells, can be promoted both by elevated CD163^+^ macrophages and PD-L1 expression (H. Zhang et al. 2023; Liu et al. 2011). Both T cell inflamed tumors (9-3 and 9-15) exhibited an increase in regulatory T cell density after therapy (**Figure 3c, 4e**). One mechanism regulatory T cells can employ to directly inhibit helper T cells is through contact-dependent signaling. We quantified the co-localization of regulatory and helper T cells using SpatialScore, a spatial biomarker associated with immunotherapy resistance in lymphoma (Phillips et al. 2021). In the two biopsy pairs with increasing in regulatory T cell density (9-3, 9-15), the SpatialScore significantly increased or remained unchanged in the on-progression biopsy, whereas the pre-treatment biopsies for 9-1A, 9-2, and 9-14 were elevated in SpatialScore and decreased after therapy (**Figure 4f**). Direct interaction of regulatory T cells can shift CD8^+^ T cells into a dysfunctional state (Schmidt, Oberle, and Krammer 2012; C. Li et al. 2020). We quantified CD8^+^ T cell differentiation states to explore the extent of CD8^+^ T cell dysfunction using single-cell positivity of PD-1 and Eomesodermin (EOMES), biomarkers typically upregulated in exhausted or terminally differentiated dysfunctional CD8^+^ T cell populations. In T cell infiltrated biopsies, the majority of the CD8^+^ populations were composed of T effector naïve cells (PD-1^-^/EOMES^-^), indicative of recent recruitment to the tumor, antigen inexperience, or overall lack of T cell priming (**Figure 4g**). 9-3 displayed a substantial increase in its population of exhausted T cells (PD-1^+^/EOMES^+^) after therapy (24.7%) (**Figure 4g**) (J. Li et al. 2018), thus indicating regulatory T cells may be impacting T cell priming capability while also pushing effector T cells toward exhaustion (Sawant et al. 2019; C. Li et al. 2020).

Our multimodal approach reveals a complex picture of both immune activation and immune suppression. Four of the six biopsy pairs analyzed with mIHC displayed substantial immune activation after CDK4/6i treatment, with two biopsy pairs having a highly T cell inflamed TIME. Immune-active tumors had increased interferon signaling, JAK/STAT activation, and increased T cell inflamed GEP scores, often in conjunction with increases in immunoreactive cell types such as cytotoxic T cells, helper T cells, and dendritic cells. Simultaneously, there was increased immunosuppressive environment after treatment in multiple tumors, including increases in CD163^+^ macrophages, PD-1/PD-L1 interactions, regulatory T cell recruitment, and T cell dysfunction. The concurrent immune activation and suppression indicates that many tumors become immunoreactive during therapy but eventually exhibit immunosuppression due to chronic inflammation. This arc from immunoreactive to immunosuppressive during therapy likely contributes to therapeutic resistance as T cell-inflamed tumors under prolonged antigen exposure led to ineffective tumor clearance (Trujillo et al. 2018; Spranger 2016).

## Discussion

CDK4/6i therapy has improved outcomes for HR+ breast cancer patients with metastatic disease, but therapeutic resistance is exceedingly common. Studies that highlight genomic and transcriptomic mechanisms of resistance to CDK4/6i identify mechanisms of resistance in only a subset of tumors (Wander et al. 2020; Cristofanilli et al. 2018; Finn et al. 2020; Park et al. 2023). To better understand molecular mechanisms of tumor adaptation and resistance to CDK4/6i, we created a longitudinal and multimodal atlas of metastatic HR+ breast cancer during CDK4/6i therapy. This atlas includes seven pairs of biopsies from five patients where each pair had a pre-CDK4/6i biopsy and on-progression biopsy from the same patient. Using six complementary bulk and single-cell molecular assays, we deeply characterized patient biopsies to build multi-dimensional atlases capturing the transition as tumors develop therapeutic resistance on CDK4/6i therapy. This atlas enabled quantification of tumor changes during therapy, and these changes suggest potential malignant cell and TIME mechanisms of resistance to CDK4/6i therapy. We validated our transcriptional results in an external cohort with pre- and post-treatment RNA-seq datasets. However, validation of our single-cell spatial results is not possible due to lack of suitable public spatial omics datasets.

Analysis of our atlas data suggests potential adaptations in metastatic HR+ breast cancer tumors and the TIME to CDK4/6i therapy (**Figure 5a**). In addition to genomic alterations acquired during therapy, malignant cell adaptations to therapy include mTORC1 pathway activation, cell-cycle bypass via increased cyclin and CDK activity, and increased replication stress. TIME adaptations included interferon and JAK/STAT signaling paired with T cell suppression and increased presence of PD-1/PD-L1 positive cells. Figures 5b and 5c summarize malignant cell and TIME adaptations on CDK4/6i therapy in each patient. At least two adaptations were identified for each tumor, and all adaptations were observed in at least two tumors. There was broad agreement across assays for support of therapeutic adaptations. While no two tumors shared the same adaptations to therapy in this limited dataset, individual adaptations were shared by subsets of tumors. Three of seven biopsy pairs, both of 9-14’s biopsy pairs and 9-15, showed marked upregulation of proliferation after therapy across many assays, indicating abrogation of the cytostatic effects of CDK4/6i therapy. Four of seven biopsy pairs showed evidence of T cell activation in the form of increased cytolytic activity, increases in overall immune cell and cytotoxic cell densities, and a decrease in regulatory T cells. However, all tumors also revealed evidence of a switch to an immunosuppressive environment, including increases in PD-1/PD-L1 activity and CD163^+^ macrophage infiltration. Malignant cell adaptations were not correlated with TIME adaptations. As the spatial omics field expands and public datasets become more readily available, validation of these observations and further characterization of the TIME will be important for future work.

**Figure 5:**
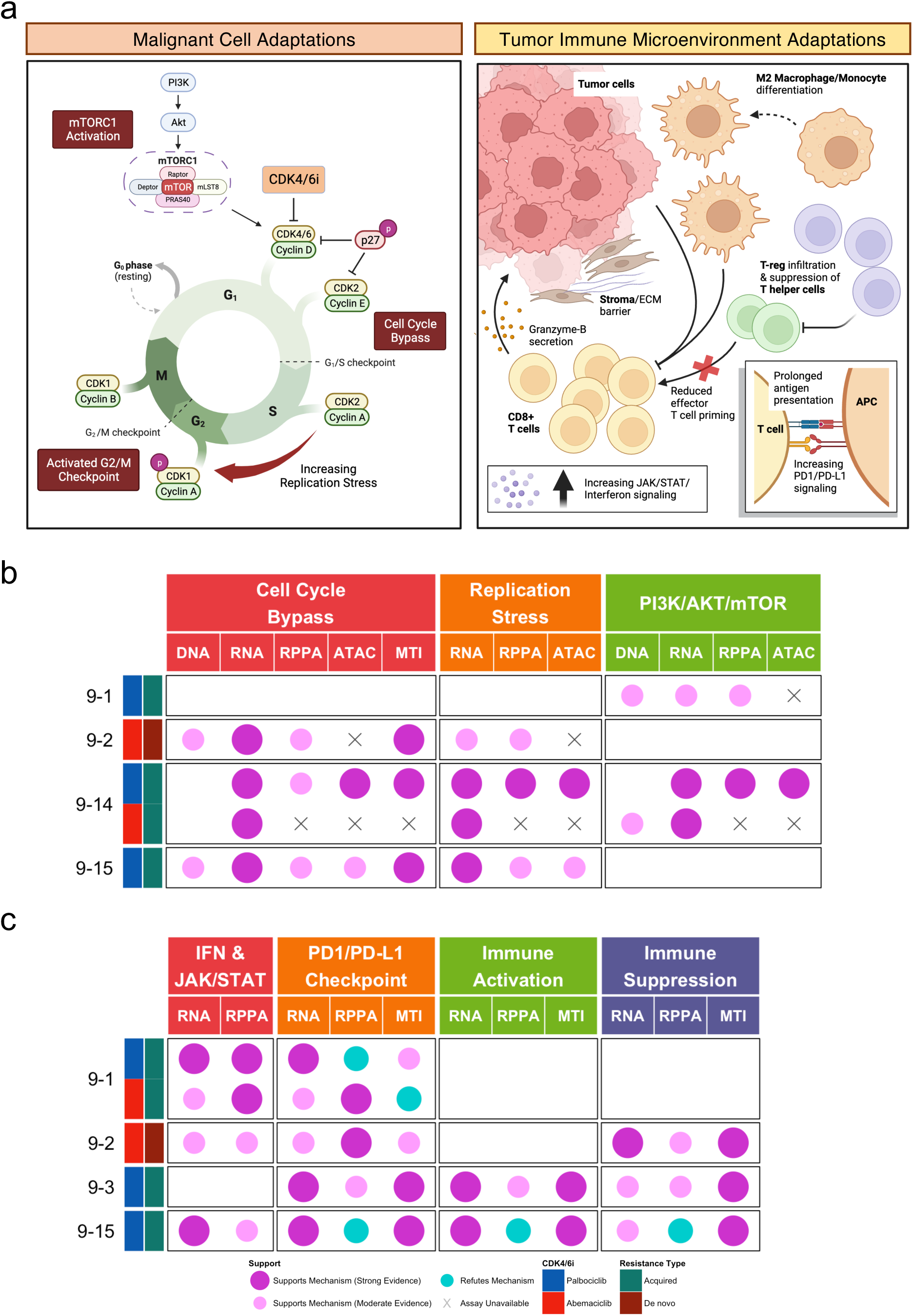
Malignant cell and tumor immune microenvironment adaptations to CDK4/6i. **a)** Diagram of malignant cell and tumor immune adaptations to CDK4/6i as suggested by multimodal profiling. Left: Upstream activation of mTORC1 may promote G1/S entry and cell cycle progression despite CDK4/6i. Activation of the G2/M checkpoint results in G2/M stalling and increased tolerance to replication stress, further promoting uncontrolled proliferation and tumor progression. Right: Evidence suggests a mix of increasing anti-tumor immunity and immunosuppression in resistant biopsies. Increased JAK/STAT and interferon signaling, increased antigen presentation, and increased CD8^+^ T cell infiltration and cytotoxicity are indicative of an active anti-tumor immune response; however, high abundance of CD163^+^ macrophage/monocyte populations, increased Regulatory T cell infiltration, increased immune checkpoint signaling may be indicative of elevated immunosuppression. Spatial organization may also contribute to reducing the efficacy of the immune response; whereby, cytotoxic cells are spatially stratified away from tumor cells due to dense stroma and extracellular matrix. **b)** Summary of patient-level malignant cell adaptations to CDK4/6i across targeted DNA-seq (DNA), bulk RNA-seq (RNA), RPPA, sciATAC-seq (ATAC), and multiplex tissue imaging (MTI). For each mechanism and patient, the strength of evidence either supporting (magenta) or refuting (cyan) the mechanism is indicated for each available assay. Patient-assay support is only indicated for mechanisms in which there is a sufficient amount of evidence across available assays for a given patient. Indicators of support are left blank for all assays within a given mechanism for tumors that displayed no evidence of that mechanism contributing to CDK4/6i adaptation. Observations for patient 9-2, a clinical non-responder, are based on their pre-treatment biopsy. Tumors (rows) are annotated by CDK4/6i therapy and clinical response. **c)** Summary of patient-level TIME adaptations to CDK4/6i across assays. Strength of evidence across assays is indicated in 5b.

Many of the adaptations observed during CDK4/6i therapy indicate potential therapeutic vulnerabilities that present opportunity for exploitation through combination therapy with CDK4/6i or via follow up therapy to CDK4/6i. MTOR inhibitors may be combined with CDK4/6i to combat increased mTORC1 activity (Michaloglou et al. 2018) as was done for patient 9-1. Inhibitors targeting other CDKs may address overactivation of the CDK2-CCNE1 complex in malignant cells. ATR or WEE1 inhibitors may be effective in tumors that show increased replication stress (Fallah et al. 2021; Ubhi and Brown 2019). Immunotherapies that block PD-1/PD-L1 checkpoint activity may dampen immune evasion by the tumor while on CDK4/6i therapy, and, indeed, prior evidence that CDK4/6i primes tumors to respond to PD-1/PD-L1 inhibitors (Deng et al. 2018). Lastly, clinical trials focused on novel targeted immunotherapies may serve to counter the increased CD163^+^ macrophage density by inhibition or reprogramming (Fendl et al. 2023). Importantly, these approaches could best be deployed in patients where the specific resistant pathway(s) is engaged as an adaptive response to the CDK4/6i.

Our study characterizes tumor adaptation to CDK4/6i at an exceptional depth making it possible to connect with and extend prior literature on molecular markers of resistance to CDK4/6i therapy. Importantly, the use of multiple bulk, single-cell, and spatial omics modalities enabled deep interrogation of adaptations over the course of CDK4/6i therapy that otherwise would not have been captured. Previous studies have found genomic aberrations occur on CDK4/6i therapy in RB1, PIK3CA, in other cyclins, and in genes driving activation of the PI3K pathways. Aberrations in these genes and pathways arose as well, albeit at a lower frequency than previously observed (Wander et al. 2020; O’Leary et al. 2018; Park et al. 2023; Cristofanilli et al. 2018). Results from bulk RNA-seq analysis are concordant with prior studies that found resistance to CDK4/6i often is associated with transcriptional activation of the mTORC1, E2F targets, replication stress, interferon signaling, and PD-1/PD-L1 pathways (Zhu et al. 2022; Park et al. 2023).

Through our use of single-cell spatial proteomics assays, we have characterized tumor changes on CDK4/6i in profound detail. Cell cycle activity and proliferation in malignant cells are associated with increased density and more spatial dispersion of proliferating malignant cells. Immune activation observed at the bulk transcriptomics and proteomics level was explored in depth to identify aspects of both immune activation and immune suppression in individual tumors. Tumors that displayed high degrees of immune activation frequently also presented signs of immunosuppression. This duality is consistent with previous observations regarding the effects of CDK4/6i on the TIME (Zhu et al. 2022; Teh and Aplin 2019), and with the progression of a T cell-inflamed TIME in which prolonged inflammation leads to immunosuppression and a dysfunctional anti-tumor immune response (Trujillo et al. 2018; Spranger 2016). The molecular atlas that we have created provides an exceptionally deep picture of how metastatic HR+ breast cancer tumors adapt and become resistant to CDK4/6i therapy.

Our study design prioritizes data depth over data breadth, and there are several limitations that arise due to this approach. Our small cohort size impedes the ability to capture the full extent of adaptations to CDK4/6i with statistical significance. In addition, the highly heterogenous nature of metastatic disease means biopsies may not represent the full complexity of disease within or across patients, opening the possibility for sampling bias and limiting statistical power. To mitigate some of these limitations, we statistically corroborated findings from our own bulk transcriptomics datasets using an external CDK4/6i-treated breast cancer cohort. This external validation of our transcriptomics results is encouraging. Unfortunately, validation using external cohorts was not possible for other assays due to the lack of suitable publicly available datasets. The variability in treatment regimen, drug class, and anatomical origin of biopsies make data interpretation challenging. Though the class of CDK4/6 therapies share targets, the specific mechanisms of action have known differences pharmacologically(Hafner et al. 2019). Further the use of other therapies in combination with CDK4/6i, including ET and mTORi, add to the clinical heterogeneity, though the observations in our cohort at least partially suggest tumor adaptations occurring independent of ET. Therapeutic adaptations specific to the TIME must also be contextualized with anatomical origin of biopsies because organs have different native immune cell populations.

Identifying biopsy timepoints at the transition from therapeutic response to resistance remains a major challenge in cancer research. While collection of a single timepoint prior to treatment is commonly used in clinical studies and trials, this approach provides a limited view for understanding the mechanisms of therapeutic adaptation. As we have demonstrated in our atlas, a longitudinal design using pre- and post-treatment biopsies provides a unique window into adaptations arising from therapy but is not well-suited for clearly separating therapeutic activity and adaptive resistance. A promising alternative strategy where paired biopsies are collected prior to treatment and shortly after therapy initiation would make it possible to differentiate the beneficial therapeutic effects from adaptive mechanisms of resistance. Neoadjuvant therapy and window-of-opportunity approaches, which are increasingly common in many cancers and when immunotherapies are used, provide opportunities for this longitudinal biopsy strategy.

Overall, this atlas demonstrates how high-depth molecular information captured across multiple scales and time can be used to further our understanding of how HR+ breast cancer tumors change because of CDK4/6i therapy. The highly individualized and heterogeneous nature of tumor adaptations in both malignant cells and the TIME observed across our tumor cohort highlights the need for detailed analysis of patient samples to optimize precision care. The data described here provide a rich resource capturing the molecular changes occurring in the tumor genome, transcriptome, and proteome and microenvironment during therapy.

### Data Availability

All data is available through the HTAN Data Portal as part of the HTAN OHSU Atlas (https://data.humantumoratlas.org/). Raw sequencing data has been deposited to dbGAP (Project phs002371.v1.p1). Images can be viewed at the Imaging Data Commons (IDC) https://portal.imaging.datacommons.cancer.gov/.

### Code Availability

No new tools were developed for the analyses in this study. Open-source workflows, R/python packages and software are described in the Methods. Code for creating the figures and panels presented in this work is available at: https://github.com/goeckslab/HTAN_HR_Figures.

## Acknowledgements

This project was carried out with major support from the Oregon Health & Science University (OHSU) SMMART Program, National Institutes of Health (NIH), National Cancer Institute (NCI) Human Tumor Atlas Network (HTAN) Research Center (U2CCA233280), and Prospect Creek Foundation.

## Author Information

### Author Contributions

JWG, GBM, JG conceived and designed the study. ZM enrolled and treated patients. JE and CW generated visualizations. JE, CW, ALC, and KM performed data and statistical analysis. JE, CW, and ALC created the figures. CW and ALC developed software and pipelines for primary data processing. JE, CW, ALC, and JG interpreted the data. SS, KC, and JRL generated data. BEJ, HSF, DG, and GVT managed sample collection, distribution, and curation. ALC managed data collection, cleaning, curation, and external sharing. ALC, BEJ, and HSF managed the project. JE, CW, ALC, and JG wrote the manuscript. NEN, ED, YHC, CLC, ZIM, PKS, GVT, LMC, ACA, JWG, GBM, and JG supervised the project. JWG, GBM, and JG acquired funding. All authors reviewed and revised the manuscript.

## Methods

### Human subjects

All biospecimens and data were collected under the single-center, observational study Molecular Mechanisms of Tumor Evolution and Resistance to Therapy (IRB#16113). The study was reviewed and approved by the Oregon Health & Science University (OHSU) Institutional Review Board (IRB). Eligibility criteria were determined by the enrolling physician and limited to participants at least 12 years old with localized, advanced, and/or metastatic cancer. All participants provided written informed consent to take part in the study. This study includes five female women, who at the time of consent were aged 70 (HTA9_1), 63 (HTA9_2), 57 (HTA9_3), 58 (HTA9_14), and 55 (HTA9_15).

### Clinical and exploratory workflows

All biospecimens were prospectively collected by clinical research coordinators at blood draws or biopsy procedures. The cold ischemia time for all biopsy tissue specimens was between two and nine minutes starting from removal of tissue from the patient. Biospecimens were either labeled for clinical testing per institutional guidelines or deidentified for research use. All biospecimens were tracked and managed using a custom implementation of the LabVantage laboratory information management system.

Clinical analytics were performed in CLIA-certified, CAP-accredited laboratories. Clinical metadata were acquired from the patient’s medical record through a combination of manual and automated data abstraction.

Biospecimen designated for research analytics underwent serial sectioning. Biospecimens preserved in formalin-fixed paraffin-embedded (FFPE) blocks were sectioned into five micron sections and distributed for H&E, multiplex immunohistochemistry, and cyclic immunofluorescence (https://dx.doi.org/10.17504/protocols.io.8epv56ppjg1b/v6). Biospecimens preserved in OCT blocks were sectioned into 5-40 micron sections and distributed for H&E, cyclic immunofluorescence, and single cell ATAC sequencing (https://dx.doi.org/10.17504/protocols.io.261ge677dl47/v4). Exploratory analytics were performed in academic laboratories or research core facilities. Assay availability across patient biospecimens varied due to insufficient tissue or failing quality control. Only biospecimens collected immediately prior to initiation of CDK4/6i therapy or after progression/therapy discontinuation were included in this manuscript. Data from additional patient biospecimens not included in this manuscript are available through the HTAN Data Portal (https://data.humantumoratlas.org/).

### Assay Methods

#### Targeted DNA sequencing

Targeted DNA-sequencing was performed at the CLIA-licensed/CAP-accredited OHSU Knight Diagnostic Laboratories using the GeneTrails Solid Tumor Panel assay. DNA was extracted from tumor-rich regions, macro-dissected from FFPE tissue. Library preparation was performed using custom QiaSeq chemistry (QIAGEN) with multiplexed PCR and sequencing was performed using an Illumina NextSeq 500/550. The DNA library was generated using 9,228 custom-designed primer extension assays covering 613,343 base pairs across 125 to 234 cancer-related genes. The panel is routinely sequenced at an average read depth of >2,000 for highly sensitive detection of SNVs, short in/dels, and copy number alterations. All variants included in this work were reported out clinically.

#### Transcriptome sequencing

RNA-sequencing was performed at the CLIA-licensed/CAP-accredited OHSU Knight Diagnostic Laboratories using the RNA Transcriptome assay. RNA was extracted from tumor-rich regions, micro-dissected from FFPE tissue. Library preparation was constructed with the TruSeq RNA Library Prep Kit and sequencing was performed on the Illumina NextSeq500/550.

#### Reverse phase protein arrays

Flash frozen tumor tissue collected from core needle biopsies was used with the Reverse Phase Protein Array (RPPA) assay (Tibes et al. 2006). Proteomic profiling of protein and phosphoprotein was performed at the MD Anderson Cancer Center Functional Proteomics RPPA Core.

#### Single-cell combinatorial indexing assay for transposase-accessible chromatin sequencing

Library preparation and sequencing: Nuclei for single-cell ATAC library preparations were performed by mincing tissue on ice followed by suspension in a Liberase reaction buffer and processed according to manufacturer’s protocols. Isolated nuclei were then distributed to a 96-well plate with 1,000 nuclei per well and brought up to 8 µL with water and 4µL 2× TD buffer (Nextera XT Kit, Illumina Inc. FC-131–1024).

For sciATAC preparations 1 µL of 2.5 µM indexed tagmentation complexes prepared as in Sinnamon et al. 2019 (Sinnamon et al. 2019). Tagmentation was performed at 55°C. For s3ATAC preparations 1 µL of 2.5 µM indexed s3 tagmentation complexes (ScaleBio Custom Order) designed using the specifications from the publication of the s3 technology (Mulqueen et al. 2021). Tagmentation was performed at 42°C. For both workflows tagmentation was carried out for 10 minutes on an Eppendorf ThermoMixer shaking at 300 rcf and then placed on ice. Nuclei were pooled and then sorted to 12 nuclei per well using DAPI fluorescence as previously described1 into wells of a 96-well plate containing 9 µL 1× TD buffer. 1 µL of 0.1% SDS was then added per well, the plate was sealed and spun down and then incubated at 55°C for 10 minutes and then placed on ice.

For sciATAC, 25 µL of NPM (Nextera XT Kit, Illumina Inc) was added per well along with 0.5uL 100X SYBR Green I (Thermo Scientific S7563) and 1 µL of indexed forward primer at 2.5 µM, 1 µL of indexed reverse primer at 2.5 µM, and ultrapure water up to 50 µL. For s3ATAC, 4 µL of NPM (Nextera XT Kit, Illumina Inc) per well was subsequently added to perform gap-fill on tagmented genomic DNA, with an incubation at 72°C for 10 minutes. 1.5 µL of 1uM A14-LNA-ME oligo was then added to supply the template for adapter switching (Mulqueen et al. 2021). Reactions were then denatured at 98°C for 30 seconds, followed by 10 cycles of 98°C for 10 seconds, 59°C for 20 seconds and 72°C for 10 seconds and then placed on ice. After adapter switching 1% (v/v) Triton-X 100 in ultrapure H2O (Sigma 93426) was added to quench persisting SDS. 2.5uL indexed i7 primer at 10 µM, 2.5 µL indexed i5 primer at 10 µM, 3 µL of ultrapure H2O, and 25 µL of NEBNext Q5U 2X Master mix (New England Biolabs M0597S), and 0.5uL 100X SYBR Green I (Thermo Scientific S7563) was then added to bring the reaction to 50 µL.

For both protocols, real time PCR was performed on a BioRad CFX, measuring SYBR fluorescence every cycle using the following program: 98°C for 30 seconds; 16–18 cycles of 98°C for 10 seconds, 55°C for 20 seconds, 72°C for 30 seconds, fluorescent reading, 72°C for 10 seconds. After fluorescence reaches an exponential growth, the samples were held at 72°C for another 30 seconds then stored at 4°C. 25 µL of each well was then pooled and cleaned up using a Qiaquick PCR purification column following manufacturer’s instructions (Qiagen 28106). Libraries were then quantified using an Agilent Tapestation 4150 D5000 tape (Agilent 5067–5592) and then sequenced on an Illumina NextSeq 2000 instrument with the following cycle conditions for sciATAC: Read 1: 50, Index 1: 8, 23 dark, 10, Index 2: 8, 27 dark, 10, Read 2: 50; and for s3ATAC: Read 1: 89, Index 1: 10, Index 2: 10, Read 2: 129.

#### Multiplex immunohistochemistry

Multiple immunohistochemistry (mIHC) was performed as previously described (Banik et al. 2020) and following (https://www.protocols.io/view/mihc-staining-ohsu-coussens-39-lab-sop-3i6gkhe.). Briefly, FFPE human tissues were sectioned at 5 microns and mounted on positively charged slides (Tanner Adhesive Slides, Mercedes Medical, TNR WHT45AD). Slides were baked overnight at 55 °C (Robbin Scientific, Model 1000) followed by an additional baking for 30 minutes at 58-60 °C. Slides were deparaffinized in xylene and graded ethanol (Xylene 2 x 5 min, 100% EtOH 2 x 2 min, 95% EtOH 2 x 2 min, 70% EtOH 2 x 2 min, 50% EtOH 1 x 2 min, diH2O 2 x 2 min), counterstained in Hematoxylin (Dako S3301), heat-mediated antigen retrieval in pH 6.0 Citra solution (BioGenex, Fremont, California, USA), blocking in Dako Dual Endogenous Enzyme Block (S2003, Dako, Santa Clara, California, USA), and protein blocking with 5% normal goat serum and 2.5% bovine serum albumin (BSA) in tris-buffered saline-0.1% Tween (TBST).

Slides were incubated with primary antibodies (**Supplemental Table 4**) for 1 hour at room temperature or 16– 17 hours at 4 °C. Primary antibody was washed off in TBST, and either anti-rat, anti-mouse, or anti-rabbit Histofine Simple Stain MAX PO horseradish peroxidase-conjugated polymer (Nichirei Biosciences, Tokyo, Japan) was applied for 30 min at room temperature, followed by AEC chromogen (Vector Laboratories, Burlingame, California, USA).

Images were acquired using the Aperio ImageScope AT (Leica Biosystems) at ×20 magnification. Following acquisition, coverslips were gently removed in 1×TBST while agitating. Removal of AEC and HRP inactivation was accomplished by incubating the slides in 0.6% fresh H2O2 in methanol for 15 minutes. AEC removal and stripping of antibodies was accomplished by ethanol gradient incubation and heat-mediated antigen retrieval such as described above between cycles. After washing and protein blocking, samples were subjected to the next round of staining. Protocol available at: https://www.protocols.io/view/mihc-staining-ohsu-coussens-39-lab-sop-3i6gkhe.

#### Cyclic immunofluorescence

Cyclic Immunofluorescence Microscopy was performed as previously described (Eng et al. 2020) and following (https://dx.doi.org/10.17504/protocols.io.23vggn6). FFPE human tissues were sectioned at 5 microns and mounted on positively charged slides (Tanner Adhesive Slides, Mercedes Medical, TNR WHT45AD). Slides were baked overnight at 55 °C (Robbin Scientific, Model 1000) and an additional 30 min at 65 °C, (Clinical Scientific Equipment NO. 100). Tissues were deparaffinized and hydrated through xylenes and graded ethanol (EtOH) as follows: xylenes (3 × 5 min), 100% EtOH (2 × 5 min), 95% EtOH (2 × 2 min), 70% EtOH (2 × 2 min), and distilled and deionized water (ddH2O, 2 × 5 min). Two-step antigen retrieval was performed in a Decloaking Chamber (Biocare Medical, Pacheco, CA), as previously described, using the following settings: setpoint 1 (SP1), 125 °C, 30 s; SP2: 90 °C, 30 s; SP limit: 10 °C variation. Following this two-step antigen retrieval, the tissues were washed in two brief changes of ddH2O (∼2 s) and then washed once for 5 min in 1x phosphate-buffered saline (PBS), pH 7.4 (Fisher, BP39920).

#### Primary antibody staining

Primary antibodies were diluted in 5% NGS and 1% BSA in 1x PBS (**Supplemental Table 4**) and applied overnight at 4 °C in a humidified chamber, covered with plastic coverslips (IHC World, IW-2601). Following overnight incubation, tissues were washed 3 × 10 min in 1x PBS and coverslipped.

#### Fluorescence Microscopy

Fluorescently stained slides were scanned on the Zeiss AxioScan.Z1 (Zeiss, Germany) with a Colibri 7/ Lumencor SpectraX-IR light source (Zeiss). Filter cubes used for image collection included DAPI (Zeiss 96 HE), Alexa Fluor 488 (AF488, Zeiss 38 HE), AF555 (Zeiss 43 HE), AF647 (Zeiss 50), and AF750 (Chroma 49007 ET Cy7). Individual slide exposure time was determined to ensure sufficient dynamic range without saturation. Full tissue scans were taken with the 20x objective (Plan-Apochromat 0.8NA WD = 0.55, Zeiss).

#### Fluorescence Signal Quenching

Following scanning, slides were soaked in a glass Coplin jar of 1x PBS until the glass coverslip could be removed without agitation, typically 10-30 minutes. Slides were placed in 10ml of freshly prepared quenching solution (20 mM sodium hydroxide (NaOH) and 3% hydrogen peroxide (H2O2) in 1x PBS) under incandescent light, for 30 minutes. Slides were removed from the chamber with forceps and washed times for 2 min in 1x PBS. The previous steps, starting with applying primary antibodies, were repeated for each round.

### Analysis Methods

#### Transcriptome sequencing

Gene Quantification: Paired-end reads were quality trimmed using Trim Galore v0.4.3. Transcripts per million (TPMs) were quantified using Kallisto v0.43.1 with annotation (GENCODE v24) (Bray et al. 2016). Gene level expression was aggregated by summing abundance of all transcripts to the corresponding gene.

Scaling/Background Cohort: Expression z-scores were computed by scaling each gene relative to a background cohort containing an additional 32 HR+ metastatic breast cancer samples along with the 13 HTAN case samples (45 total). Z-scores were calculated by scaling and mean centering the log2(TPM) of each gene.

#### Gene Expression Signatures

##### Molecular Subtype Signature

Molecular subtypes were classified using the PAM50 subtype gene signature (Parker et al. 2009). A background cohort consisting of 120 breast cancer samples including the 13 samples our cohort was mean centered and used for classifying subtypes. Spearman correlations were computed between the patient samples and the pre-defined centroids of PAM50 subtypes (Parker et al. 2009). Subtypes were assigned to each sample using the highest Spearman correlation to the subtype centroid.

##### G1 Arrest Scores

G1 arrest scores were computed from bulk RNA-seq using a previously defined gene signature of cell cycle arrest by CDK4/6i (Hafner et al. 2019). Briefly, the change in expression z-score for 87 genes during CDK4/6i was averaged across each pair of pre- and post-treatment samples (n = 7) and multiplied by negative one. A positive score indicates G1 arrest while a negative score indicates G1 entry.

##### T cell Inflamed GEP Scores

Using bulk RNA-seq T cell inflamed GEP scores were computed for each sample by taking the weighted sum of a previously defined set of 18 adaptive immune genes related to IFN-gamma response, antigen presentation, chemokine expression, and cytotoxicity(Ayers et al. 2017; Cristescu et al. 2018).

#### VIPER Regulatory Network

Regulator activity was inferred with the VIPER algorithm using a TCGA BRCA ARACNe network (Alvarez et al. 2016; Lachmann et al. 2016). Briefly, VIPER infers the activity of each regulon based on the expression of its downstream targets as defined by the ARACNe network. Gene expression data used for VIPER was mean centered and scaled using a background cohort of 45 HR+ samples.

#### Pathway enrichment analysis

The gene set variation analysis (GSVA) algorithm was used to infer pathway activity for the 50 MSigDB Cancer Hallmark Pathways. Pathway enrichment scores were also computed with GSVA using additional curated pathways relevant to CDK4/6i adaptation from Reactome, KEGG, and BioCarta (Hänzelmann, Castelo, and Guinney 2013; Liberzon et al. 2015; Jassal et al. 2020; Kanehisa et al. 2016). These pathways include gene sets related to cell cycle regulation, mTOR activation, and JAK/STAT signaling. GSVA was performed on the log2 transformed gene expression matrix of 45 HR+ samples. The GSVA algorithm was ran using a Gaussian kernel for cumulative density function estimation and a normalized enrichment statistic of the random walk deviations. Immune cell activity was also inferred from bulk RNA-seq with the GSVA algorithm using 16 pre-defined gene sets of immune cell types (Tamborero et al. 2018). Three additional immune cell signatures were also included for GSVA to compute transcriptional enrichment scores representing chemokine activity, antigen presentation, and T cell inflammation (Coppola et al. 2011; Prabhakaran et al. 2017; Wang et al. 2019; Cristescu et al. 2018).

### Clustering analysis

To identify subgroups of patients associated with malignant cell and TIME adaptations, hierarchical clustering was performed using GSVA enrichment scores from seven paired pre- and post-treatment RNA-seq samples integrated with twenty-three paired samples from the external RNA-seq dataset(Park et al. 2023). Malignant cell high (MC-high) and low (MC-low) groups were identified from hierarchical clustering using the ten malignant cell pathways from Figure 2b and selected using the cutree (k = 2) method from the stats library. Similarly, TIME high (TIME-high) and low (TIME-low) groups were identified from clustering eight immune-related gensets from MSigDB Cancer Hallmark Pathways and selected using cutree (k = 2).

#### Reverse phase protein arrays

RPPA data was first normalized for protein loading for all samples within a set followed by debatching across sets to TCGA RPPA breast cancer samples using replicate-based normalization (RBN) (Akbani et al. 2014). Prior to plotting, abundance values were z-scored by scaling and mean centering relative to a background cohort consisting of the 12 cohort samples with available RPPA data plus an additional 13 HR+ metastatic breast cancer samples. These protein z-scores were used for all downstream analysis. Protein pathway activity scores were developed using previously described predictors (Johnson et al. 2022; Labrie et al. 2019). Pathway scores were computed for each sample by taking the summation of each protein predictor in a defined pathway set, times their assigned weight (positive or negative), divided by the total number of predictors in that set (**Supplemental Table 5**).

#### Single-cell combinatorial indexing assay for transposase-accessible chromatin sequencing

Sequence reads were demultiplexed using unidex (https://github.com/adeylab/unidex) matching index reads to the PCR indexes, and then for s3ATAC, the first 8 bp of Read 2 to the Tn5 index, and removing bases 9-29 from Read 2 which includes the Tn5 Mosaic End recognition sequence. Sequence data alignment and processing: FASTQ files for each biopsy were separately aligned to the GRCh38/hg38 (GCA_000001405.15) via BWA-MEM (v0.7.17-r1198-dirty) (H. Li 2013). We used the hg38 analysis set, which contains features specifically for next-generation sequencing alignment such as masking of genomic repeat arrays and a contig to redirect reads obtained from Epstein-Barr virus. Aligned reads were filtered with the scitools (Sinnamon et al. 2019) function ‘bam-rmdup’, which removes poorly mapped reads (quality score < 10) and collapses PCR duplicated reads on a per-barcode basis. BAM files were then split into single BAM files for each tumor and condition (pre-treatment and on-progression) as appropriate through scitools ‘bam-split’, and each BAM file was indexed with SAMtools (v1.10) (Danecek et al. 2021).

##### ArchR object generation and quality control

All single cell ATAC-seq work post-alignment was performed with the R package ArchR (v1.0.2) (Granja et al. 2021). “Arrow” files, the fundamental data storage file for ArchR projects, were generated for each biopsy’s BAM file through the ArchR function createArrowFiles(). Barcodes that did not meet a minimum of 3.5 log10 unique fragments or a lenient TSS enrichment cutoff of 1.5 were removed. To eliminate doublets, we ran addDoubletScores() and filterDoublets() with default settings on each Arrow file. With the exception of the pre-treatment biopsy for 9-3, which had too few barcodes for robust doublet calling, each sample had fewer than 100 doublets identified and removed. ArchR calculates per-barcode TSS enrichment and fragment counts during Arrow file generation; after our filtering was completed, patient 9-14 (Bx1 and Bx2) and 9-15 (Bx2) datasets exceeded 6.5 median TSS enrichment and 19k median fragments. Patient 9-3 sample pair were comparatively lower in quality, achieving 2.1 & 2.8 median TSS enrichment and 19k & 9.9k median fragments, respectively. Furthermore, while all on-progression datasets retained at least 1.8k passing barcodes, pre-treatment datasets for 9-3 and 9-14 had fewer than 500 each. We therefore prioritized barcode count over additional quality filters and retained a permissive TSS enrichment cutoff (1.5) for all downstream analysis.

##### Dimensionality reduction and clustering of scATAC data

We reduced the high dimensionality of our single cell ATAC data with the iterative Latent Semantic Indexing (LSI) implementation provided by ArchR. In the standard ArchR workflow, genome-wide ATAC signal is split into 500 bp tiles for each barcode; the most accessible of these tiles are then used for the initial LSI. Preliminary clusters identified from the first round of LSI are used to identify highly variable tiles that are used for the next LSI iteration. To perform iterative LSI, we combined all scATAC-seq datasets and ran addIterativeLSI() on the top 100,000 variable tiles. Four rounds of LSI were performed at steadily increasing resolution (0.5, 1, 1.5, and 2) to help minimize batch effects. Finally, clusters of single cells were identified from the reduced dimensions via addClusters() and visualized in a 2D embedding through addUMAP() with minDist set to 0.3.

##### Cell typing of single cell ATAC clusters

To identify cell types in our single cell ATAC-seq data, we first identified marker genes for each cluster by providing the ArchR gene score matrix to getMarkerFeatures(). ArchR gene scores are a metric of overall chromatin accessibility for genes based on the proximity of ATAC signal within and around the gene body (Granja et al. 2021). Manual examination of significantly enriched genes (FDR < 0.05, log2 fold change > 0.5) for each cluster revealed known cell type marker genes from which we made an initial classification. We further verified marker genes by manually inspecting ATAC-seq reads for each gene in genome browser view via plotBrowserTrack(). Cluster 7 was classified as endothelial based on enriched markers VWF and PECAM1 (CD31); cluster 8 was labeled immune from enrichment of PTPRC (CD45); cluster 9 was identified as fibroblast from enriched mesenchymal markers LUM, PDGFRA/B, and collagen-encoding genes. Hepatocytes (clusters 4-5) were labeled based on enriched markers ALB, TTR, and CRP. No marker genes were found for cluster 6 due to low cell count (n = 29), but we identified it as cholangiocyte based on 1) manual inspection of ATAC signal surrounding cholangiocyte markers SOX9, CFTR, KRT7 (CK7), and KRT19 (CK19), and 2) by merit of cluster 6 barcodes originating solely from liver biopsy datasets. Epithelial/tumor clusters (clusters 1, 11, and 13-15) were labeled based on their marker gene enrichment (CDH1, EPCAM, or keratins) and by merit of their barcode composition being specific to an individual patient. Clusters 10 and 12 were additionally labeled as epithelial/tumor due to evident proximity to epithelial/tumor clusters in UMAP space and corresponding patient specificity.

##### Gene set module scores

We calculated module scores for gene sets through the ArchR function addModuleScores(). Given a set of genes, this function uses the ArchR gene score matrix to generate a quantitative measurement of chromatin accessibility for the gene group. MSigDB (Subramanian et al. 2005; Liberzon et al. 2015) “Hallmark” gene sets used for the module score analysis were obtained using the R package msigdbr (v7.5.1) (https://igordot.github.io/msigdbr/). Modules corresponding to Reactome (Matthews et al. 2009) pathways were obtained from “C2” MSigDB curated gene sets (Liberzon et al. 2011). Only tumor cells phenotyped as “tumor” were included in statistical testing.

#### Multiplex immunohistochemistry

Primary image processing steps were performed using the Galaxy-MCMICRO Environment (GalaxyME) (Creason et al. 2022). For each biopsy, single-channel SVS images of the whole slide for each protein assayed were aligned to a reference Hematoxylin image using PALOM (version 2022.9.1). Registration quality was validated using Avivator. Mesmer (Deepcell version 0.12.3) was used to perform nuclear and whole-cell segmentation on the resulting registered ome.tiff image using Hematoxylin, pan-cytokeratin, CD45, and CD68 marker channels. Nuclear and whole-cell masks were merged, and lone nuclei that could not be paired with a corresponding whole-cell mask were dilated by three pixels to estimate whole-cell boundaries. MCMICRO’s quantification module was used to extract single cell mean marker intensities and morphological features from the refined cell mask and registered image. The quantified cell feature table was converted to AnnData format using Scimap. The single-cell intensities were log-transformed and scaled between [0-1] using Sci-kit Learn’s MinMaxScaler function. Gates were selected for each marker based on the distribution of rescaled intensity values across cells in the biopsies and validated visually in the image using the Vitessce viewer (Keller et al. 2021). Cell phenotypes were called hierarchically using the assigned gates, and according to the phenotype map (**Supplemental Table 3**).

#### Cyclic immunofluorescence

Acquired images were stitched and registered across cycles based on the DAPI features using Zen Slidescan (v2.3). Each registered scene image underwent processing using a GalaxyME pipeline. Whole-cell masks were generated using Mesmer (Deepcell version 0.12.3). DAPI was used as a nuclear marker, and a maximum intensity projection of cytokeratin (CK19, CK17, CK14, CK8, CK7, CK5), CD45, and CD68 channels was used as a membrane marker. Feature extraction and intensity scaling was performed as described for mIHC. Non-hierarchical gates were selected for all markers in the panel, and downstream analyses were performed using the resulting marker-positivity calls. Gating and validation were performed as described for mIHC.

### Spatial analysis

#### Cell density calculation

Cell densities are reported as the number of cells of a given cell type divided by the area of the tissue in mm^2^. For both imaging assays, cell centroid coordinates were converted from pixels to micrometers using the physical resolution of the platform (cycIF = 0.325 µm/pixel, mIHC = 0.5µm/pixel). Tissue area was calculated by generating a hull (alpha parameter = 0.05) using the Python package Alphashape (version 1.3.1) (https://doi.org/10.5281/zenodo.4697576) around the centroids of all segmented cells in the biopsy. The area of the hull was converted from µm^2^ to mm^2^.

#### Ripley’s K of proliferating tumor cells

Cytokeratin-positive tumor cells were classified as proliferative based on positivity for the cell-cycle markers Ki67, PCNA, and pHH3. The R package spatialTIME (version 1.3.3.3) (Creed et al. 2021) was used to calculate univariate Ripley’s K metrics for proliferating tumor cells for each whole slide image within a radius of 50µm. Radii between 0-100µm were tested for this analysis, and 50µm was selected to maximize inter-sample variation in the degree of clustering. This resulted in a single value per biopsy that describes the distribution of proliferating tumor cells as being randomly distributed, clustered, or dispersed in space relative to a sample-specific permuted random point process (edge_correction = ‘translation’, num_permutations = 100).

#### Grid-based spatial heterogeneity

Fishnet grids of each biopsy, with a tile size of 0.01mm^2^, were generated using the Python package GeoPandas (version 0.13.2) (http://doi.org/10.5281/zenodo.3946761). Tiles with fewer than 20 cells total were excluded from downstream analysis. Per-tile cell density distributions were calculated to assess spatial heterogeneity across the tissues.

#### Minimum distance calculations

The *spatial_distance* function from the Python package Scimap (version 1.0.0) was used to compute minimum distances between cell types of interest.

#### SpatialScore calculation

The SpatialScore first reported in (Phillips et al. 2021), is defined as the ratio of the distance between a helper T cell and its nearest tumor cell to the distance between a helper T cell and its nearest regulatory T cell, reported for all helper T cells in each biopsy. The Minimum distances that comprise the SpatialScore numerator and denominator were calculated using Scimap as described above.

### Statistical Testing

All Mann-Whitney tests were performed in R using the wilcox.test function from the stats library. Two-sided tests were performed on GSVA pathway enrichment scores to validate findings in the external RNA-seq dataset(Park et al. 2023) and were adjusted for multiple testing using the false discovery rate (FDR) method. One-sided Mann-Whitney tests were performed on per-cell open chromatin accessibility enrichment scores from sciATAC-seq to evaluate concordance of pathway enrichment from bulk RNA-seq. For each tumor sample, all cells within that sample were tested against the remaining cells from all other samples using the “greater” hypothesis method and p-values were adjusted for multiple testing using FDR. Samples with an FDR < 0.01 were considered to be statistically significant. Only cells phenotyped as tumor were included for testing. The number of tumor cells in each sample can be seen in **Extended Data** Figure 5b. Two-sided Mann-Whitney tests were used between patient tumor pairs for all single-cell cycIF and mIHC data. Pairs with a p-value of < 0.01 were considered statistically significant.

### Visualization

All heatmaps and oncoplots containing bulk omics data (RNA-seq, RPPA, targeted DNA-seq) were created using the ComplexHeatmap R package (Gu, Eils, and Schlesner 2016). All box plots, violin plots, and dot plots for sciATAC-seq and multiplex tissue imaging (mIHC, cycIF) were created using the ggplot2 R package (Wickham 2009). CycIF and mIHC image visualizations were captured using Avivator (Manz et al. 2022). All diagrams (Figure 1a, Figure 3d, and Figure 5a) were created with BioRender.com.

## Extended Data Figure Legends

**Extended Data** Figure 1**: Genomic Landscape of HR+ Metastatic Breast Cancer on CDK4/6i Therapy.** Oncoplot of somatic gene alterations detected using high depth targeted DNA-seq from patient tumors before and after CDK4/6i. All genes altered in at least one patient biopsy are included in the plot with rows ordered highest to lowest by frequency within the cohort. Patient tumors are annotated by therapy (CDK4/6i and ERi), initial response, and PAM50 classification.

**Extended Data** Figure 2**: Gene Set Enrichment Activity of HR+ Metastatic Breast Cancer on CDK4/6i Therapy.** Heatmap of gene set enrichment for patient tumors before and after CDK4/6i. Gene sets include the 50 MSigDB Cancer Hallmark pathways and select curated pathways relevant to CDK4/6i adaptation from Reactome, KEGG, and BioCarta(Hänzelmann, Castelo, and Guinney 2013; Liberzon et al. 2015). Pathways (rows) are color annotated by category. Patient tumors (columns) are annotated by therapy (CDK4/6i and ERi), initial response, and PAM50 classification.

**Extended Data** Figure 3**: Gene Set Enrichment Activity of Malignant Cell Pathways During CDK4/6i Therapy in External Cohort of Metastatic Breast Cancers.** Heatmap describes the change in pathway enrichment during CDK4/6i therapy across 23 patient cases with paired baseline (BL) and progressive disease (PD) samples from Park et al. (2023)(Park et al. 2023). Columns are annotated with transcriptional G1 arrest scores representing the extent to which CDK4/6i prevents transcription for cell cycle progression. Pathways with an average change in activity greater than 0.5 between G1 positive and negative HTAN tumors were validated in positive and negative tumors from the external cohort (FDR < 0.01, two-sided Mann-Whitney test). Pathways (rows) are grouped by category. Biopsy pairs (columns) are split by G1 arrest phenotype.

**Extended Data** Figure 4**: Transcriptional profiling of multiple cohorts reveals distinct subgroups of malignant cell and TIME adaptations to CDK4/6i therapy.** Heatmap describes change in GSVA pathway enrichment during lines of CDK4/6i therapy in HTAN and Park et al. datasets. Hierarchical clustering of HTAN and Park et al. cases revealed two distinct clusters with either increasing or decreasing activation of malignant cell pathways associated with CDK4/6i adaptation (**Figure 2B; Extended Data** Figure 6). Similarly, hierarchical clustering of immune-related pathways from MSigDB hallmarks collection(Liberzon et al. 2015) also revealed two clusters of TIME-related changes. Clusters were annotated as either MC-high/TIME-High, MC-high/TIME-Low, MC-low/TIME-High, or MC-low/TIME-Low. Pathways (rows) are grouped by category. Biopsy pairs (columns) are split by cluster assignment.

**Extended Data** Figure 5**: Master Regulator Profiles of HR+ Metastatic Breast Cancer on CDK4/6i Therapy.** Heatmap of regulator activity (z-scores) for patient tumors before and after CDK4/6i. Regulator activity was inferred using the Viper algorithm. The top 25% regulators with the highest variance across the cohort were included in the heatmap. Patient tumors (columns) are annotated by therapy (CDK4/6i and ERi), initial response, and PAM50 classification.

**Extended Data** Figure 6**: Distribution of Per-cell Chromatin Accessibility Pathway Enrichment from sciATAC-seq.** Violin plots represent the distribution of per-cell module chromatin accessibility enrichment across patient tumors for the 50 MSigDB Cancer Hallmark pathways and select curated pathways relevant to CDK4/6i adaptation from Reactome, KEGG, and BioCarta(Hänzelmann, Castelo, and Guinney 2013; Liberzon et al. 2015). For all violin plots, the mean value is shown as a point, and the first (25%) and third quartiles (75%) are represented as vertical lines within each distribution. Only cells phenotyped as tumor are included in each plot. Red asterisks indicate statistically significant enrichment in each tumor compared to all other tumor cells (FDR < 0.01, one-sided Mann-Whitney test).

**Extended Data** Figure 7**: sciATAC-seq Sample Counts and Cell Type Composition. a)** UMAP coordinates from sciATAC-seq color annotated by patient tumors (top) and inferred cell type (bottom). **b)** Distribution of cell counts by inferred cell type across patient tumors.

**Extended Data** Figure 8**: Transcriptional activity of immune pathways and gene sets between metastatic tumor sites from pre-treatment and on-progression biopsies in HTAN and Park et al. 2023 datasets. a)** PCA plot of 59 biopsies colored by metastatic site from HTAN and Park using immune pathways and cell types quantified with GSVA. **b)** Distribution of GSVA activity scores by tissue type for immune pathways and cell types using samples from HTAN and Park. Tissue types with a significant difference (FDR < 0.05) in activation from a two-sided Mann-Whitney (one vs all) for a particular gene set are indicated with an asterisk (*) above each plot. Only tissue sites with at least five samples were included for testing.

**Extended Data** Figure 9**: Immune cell activity of HTAN and Park cases during CDK4/6i. a)** Change in enrichment for 16 immune cell types sets(Tamborero et al. 2018) during lines of CDK4/6i therapy in HTAN and Park et al. datasets. Tumors are annotated by change in T cell inflamed GEP score and TIME cluster. **b)** Relative enrichment for 16 immune cell types sets in pre-treatment and on-progression HTAN biopsies. Each biopsy is annotated by T cell inflamed GEP score.

**Extended Data** Figure 10**: Proteomic Pathway Activity Profiles of HR+ Metastatic Breast Cancer on CDK4/6i Therapy.** Heatmap of proteomic pathway signatures for patient tumors before and after CDK4/6i. Pathway scores were calculated using previously defined protein sets(Johnson et al. 2022; Labrie et al. 2019). Patient tumors (columns) are annotated by therapy (CDK4/6i and ERi), initial response, and PAM50 classification.

## References

1. Akbani, Rehan, Patrick Kwok Shing Ng, Henrica M. J. Werner, Maria Shahmoradgoli, Fan Zhang, Zhenlin Ju, Wenbin Liu, et al. 2014. “A Pan-Cancer Proteomic Perspective on The Cancer Genome Atlas.” Nature Communications 5 (1): 3887.

2. Al-Qasem, Abeer J., Carla L. Alves, Sidse Ehmsen, Martina Tuttolomondo, Mikkel G. Terp, Lene E. Johansen, Henriette Vever, et al. 2022. “Co-Targeting CDK2 and CDK4/6 Overcomes Resistance to Aromatase and CDK4/6 Inhibitors in ER+ Breast Cancer.” Npj Precision Oncology 6 (1): 68.

3. Alvarez, Mariano J., Yao Shen, Federico M. Giorgi, Alexander Lachmann, B. Belinda Ding, B. Hilda Ye, and Andrea Califano. 2016. “Functional Characterization of Somatic Mutations in Cancer Using Network-Based Inference of Protein Activity.” Nature Genetics 48 (8): 838–47.

4. Ayers, Mark, Jared Lunceford, Michael Nebozhyn, Erin Murphy, Andrey Loboda, David R. Kaufman, Andrew Albright, et al. 2017. “IFN-γ-Related MRNA Profile Predicts Clinical Response to PD-1 Blockade.” The Journal of Clinical Investigation 127 (8): 2930–40.

5. Banik, Grace, Courtney B. Betts, Shannon M. Liudahl, Shamilene Sivagnanam, Rie Kawashima, Tiziana Cotechini, William Larson, et al. 2020. “High-Dimensional Multiplexed Immunohistochemical Characterization of Immune Contexture in Human Cancers.” Methods in Enzymology 635: 1–20.

6. Boutilier, Ava J., and Sherine F. Elsawa. 2021. “Macrophage Polarization States in the Tumor Microenvironment.” International Journal of Molecular Sciences 22 (13): 6995.

7. Bray, Nicolas L., Harold Pimentel, Páll Melsted, and Lior Pachter. 2016. “Near-Optimal Probabilistic RNA-Seq Quantification.” Nature Biotechnology 34 (5): 525–27.

8. Caldon, C. Elizabeth, C. Marcelo Sergio, Jian Kang, Anita Muthukaruppan, Marijke N. Boersma, Andrew Stone, Jane Barraclough, et al. 2012. “Cyclin E2 Overexpression Is Associated with Endocrine Resistance but Not Insensitivity to CDK2 Inhibition in Human Breast Cancer Cells.” Molecular Cancer Therapeutics 11 (7): 1488–99.

9. Ciołczyk-Wierzbicka, Dorota, Dorota Gil, Marta Zarzycka, and Piotr Laidler. 2020. “MTOR Inhibitor Everolimus Reduces Invasiveness of Melanoma Cells.” Human Cell 33 (1): 88–97.

10. Condorelli, R., L. Spring, J. O’Shaughnessy, L. Lacroix, C. Bailleux, V. Scott, J. Dubois, et al. 2018. “Polyclonal RB1 Mutations and Acquired Resistance to CDK 4/6 Inhibitors in Patients with Metastatic Breast Cancer.” Annals of Oncology 29 (3): 640–45.

11. Coppola, Domenico, Michael Nebozhyn, Farah Khalil, Hongyue Dai, Timothy Yeatman, Andrey Loboda, and James J. Mulé. 2011. “Unique Ectopic Lymph Node-like Structures Present in Human Primary Colorectal Carcinoma Are Identified by Immune Gene Array Profiling.” The American Journal of Pathology 179 (1): 37–45.

12. Creason, Allison L., Cameron Watson, Qiang Gu, Daniel Persson, Luke Sargent, Yu-An Chen, Jia-Ren Lin, et al. 2022. “A Web-Based Software Resource for Interactive Analysis of Multiplex Tissue Imaging Datasets.” BioRxiv. 10.1101/2022.08.18.504436.

13. Creed, Jordan H., Christopher M. Wilson, Alex C. Soupir, Christelle M. Colin-Leitzinger, Gregory J. Kimmel, Oscar E. Ospina, Nicholas H. Chakiryan, et al. 2021. “SpatialTIME and ITIME: R Package and Shiny Application for Visualization and Analysis of Immunofluorescence Data.” *Bioinformatics (Oxford*, England) 37 (23): 4584–86.

14. Cristescu, Razvan, Robin Mogg, Mark Ayers, Andrew Albright, Erin Murphy, Jennifer Yearley, Xinwei Sher, et al. 2018. “Pan-Tumor Genomic Biomarkers for PD-1 Checkpoint Blockade-Based Immunotherapy.” Science (New York, N.Y.) 362 (6411): eaar3593.

15. Cristofanilli, Massimo, Angela DeMichele, Carla Giorgetti, Nicholas C. Turner, Dennis J. Slamon, Seock-Ah Im, Norikazu Masuda, et al. 2018. “Predictors of Prolonged Benefit from Palbociclib plus Fulvestrant in Women with Endocrine-Resistant Hormone Receptor–Positive/Human Epidermal Growth Factor Receptor 2–Negative Metastatic Breast Cancer in PALOMA-3.” European Journal of Cancer (Oxford, England: 1990) 104 (November): 21–31.

16. Crozier, Lisa, Reece Foy, Brandon L. Mouery, Robert H. Whitaker, Andrea Corno, Christos Spanos, Tony Ly, Jeanette Gowen Cook, and Adrian T. Saurin. 2022. “CDK4/6 Inhibitors Induce Replication Stress to Cause Long-Term Cell Cycle Withdrawal.” The EMBO Journal 41 (6): e108599.

17. Danecek, Petr, James K. Bonfield, Jennifer Liddle, John Marshall, Valeriu Ohan, Martin O. Pollard, Andrew Whitwham, et al. 2021. “Twelve Years of SAMtools and BCFtools.” GigaScience 10 (2). 10.1093/gigascience/giab008.

18. De Angelis, Carmine, Xiaoyong Fu, Maria Letizia Cataldo, Agostina Nardone, Resel Pereira, Jamunarani Veeraraghavan, Sarmistha Nanda, et al. 2021. “Activation of the IFN Signaling Pathway Is Associated with Resistance to CDK4/6 Inhibitors and Immune Checkpoint Activation in ER-Positive Breast Cancer.” Clinical Cancer Research: An Official Journal of the American Association for Cancer Research 27 (17): 4870–82.

19. Dean, J. L., C. Thangavel, A. K. McClendon, C. A. Reed, and E. S. Knudsen. 2010. “Therapeutic CDK4/6 Inhibition in Breast Cancer: Key Mechanisms of Response and Failure.” Oncogene 29 (28): 4018–32.

20. Deng, Jiehui, Eric S. Wang, Russell W. Jenkins, Shuai Li, Ruben Dries, Kathleen Yates, Sandeep Chhabra, et al. 2018. “CDK4/6 Inhibition Augments Antitumor Immunity by Enhancing T-Cell Activation.” Cancer Discovery 8 (2): 216–33.

21. Eng, Jennifer, Guillaume Thibault, Shiuh-Wen Luoh, Joe W. Gray, Young Hwan Chang, and Koei Chin. 2020. “Cyclic Multiplexed-Immunofluorescence (CmIF), a Highly Multiplexed Method for Single-Cell Analysis.” *Methods in Molecular Biology (Clifton*, N.J.) 2055: 521–62.

22. Esbona, Karla, Yanyao Yi, Sandeep Saha, Menggang Yu, Rachel R. Van Doorn, Matthew W. Conklin, Douglas S. Graham, et al. 2018. “The Presence of Cyclooxygenase 2, Tumor-Associated Macrophages, and Collagen Alignment as Prognostic Markers for Invasive Breast Carcinoma Patients.” The American Journal of Pathology 188 (3): 559–73.

23. Fallah, Yassi, Diane M. Demas, Lu Jin, Wei He, and Ayesha N. Shajahan-Haq. 2021. “Targeting WEE1 Inhibits Growth of Breast Cancer Cells That Are Resistant to Endocrine Therapy and CDK4/6 Inhibitors.” Frontiers in Oncology 11 (July): 681530.

24. Fendl, B., A. S. Berghoff, M. Preusser, and B. Maier. 2023. “Macrophage and Monocyte Subsets as New Therapeutic Targets in Cancer Immunotherapy.” ESMO Open 8 (1): 100776.

25. Finn, Richard S., Yuan Liu, Zhou Zhu, Miguel Martin, Hope S. Rugo, Véronique Diéras, Seock-Ah Im, et al. 2020. “Biomarker Analyses of Response to Cyclin-Dependent Kinase 4/6 Inhibition and Endocrine Therapy in Women with Treatment-Naïve Metastatic Breast Cancer.” Clinical Cancer Research: An Official Journal of the American Association for Cancer Research. American Association for Cancer Research (AACR).

26. Finn, Richard S., Miguel Martin, Hope S. Rugo, Stephen E. Jones, Seock-Ah Im, Karen A. Gelmon, Nadia Harbeck, et al. 2016. “PALOMA-2: Primary Results from a Phase III Trial of Palbociclib (P) with Letrozole (L) Compared with Letrozole Alone in Postmenopausal Women with ER+/HER2– Advanced Breast Cancer (ABC).” Journal of Clinical Oncology: Official Journal of the American Society of Clinical Oncology 34 (15_suppl): 507–507.

27. Fu, Tong, Lei-Jie Dai, Song-Yang Wu, Yi Xiao, Ding Ma, Yi-Zhou Jiang, and Zhi-Ming Shao. 2021. “Spatial Architecture of the Immune Microenvironment Orchestrates Tumor Immunity and Therapeutic Response.” Journal of Hematology & Oncology 14 (1): 98.

28. Goel, Shom, Molly J. DeCristo, April C. Watt, Haley BrinJones, Jaclyn Sceneay, Ben B. Li, Naveed Khan, et al. 2017. “CDK4/6 Inhibition Triggers Anti-Tumour Immunity.” Nature 548 (7668): 471–75.

29. Goetz, Matthew P., Masakazu Toi, Suzanne Klise, Martin Frenzel, Nawel Bourayou, and Angelo Di Leo. 2015. “MONARCH 3: A Randomized Phase III Study of Anastrozole or Letrozole plus Abemaciclib, a CDK4/6 Inhibitor, or Placebo in First-Line Treatment of Women with HR+, HER2-Locoregionally Recurrent or Metastatic Breast Cancer (MBC).” Journal of Clinical Oncology: Official Journal of the American Society of Clinical Oncology 33 (15_suppl): TPS624–TPS624.

30. Goswami, Kuntal Kanti, Anamika Bose, and Rathindranath Baral. 2021. “Macrophages in Tumor: An Inflammatory Perspective.” Clinical Immunology (Orlando, Fla.) 232 (108875): 108875.

31. Granja, Jeffrey M., M. Ryan Corces, Sarah E. Pierce, S. Tansu Bagdatli, Hani Choudhry, Howard Y. Chang, and William J. Greenleaf. 2021. “ArchR Is a Scalable Software Package for Integrative Single-Cell Chromatin Accessibility Analysis.” Nature Genetics 53 (3): 403–11.

32. Griffiths, Jason I., Jinfeng Chen, Patrick A. Cosgrove, Anne O’Dea, Priyanka Sharma, Cynthia Ma, Meghna Trivedi, et al. 2021. “Serial Single-Cell Genomics Reveals Convergent Subclonal Evolution of Resistance as Early-Stage Breast Cancer Patients Progress on Endocrine plus CDK4/6 Therapy.” Nature Cancer 2 (6): 658–71.

33. Gu, Zuguang, Roland Eils, and Matthias Schlesner. 2016. “Complex Heatmaps Reveal Patterns and Correlations in Multidimensional Genomic Data.” *Bioinformatics (Oxford*, England) 32 (18): 2847–49.

34. Hafner, Marc, Caitlin E. Mills, Kartik Subramanian, Chen Chen, Mirra Chung, Sarah A. Boswell, Robert A. Everley, et al. 2019. “Multiomics Profiling Establishes the Polypharmacology of FDA-Approved CDK4/6 Inhibitors and the Potential for Differential Clinical Activity.” Cell Chemical Biology 26 (8): 1067–1080.e8.

35. Hänzelmann, Sonja, Robert Castelo, and Justin Guinney. 2013. “GSVA: Gene Set Variation Analysis for Microarray and RNA-Seq Data.” BMC Bioinformatics 14 (1): 7.

36. Herrera-Abreu, Maria Teresa, Marta Palafox, Uzma Asghar, Martín A. Rivas, Rosalind J. Cutts, Isaac Garcia-Murillas, Alex Pearson, et al. 2016. “Early Adaptation and Acquired Resistance to CDK4/6 Inhibition in Estrogen Receptor-Positive Breast Cancer.” Cancer Research 76 (8): 2301–13.

37. Janni, Wolfgang, Howard A. Burris, Kimberly L. Blackwell, Lowell L. Hart, Arlene Chan, Arnd Nusch, Olga Nikolaevna Burdaeva, et al. 2017. “First-Line Ribociclib plus Letrozole for Postmenopausal Women with Hormone Receptor-Positive (HR+), HER2-Negative (HER2-) Advanced Breast Cancer (ABC): MONALEESA-2 Safety Results.” Journal of Clinical Oncology: Official Journal of the American Society of Clinical Oncology 35 (15_suppl): 1047–1047.

38. Jassal, Bijay, Lisa Matthews, Guilherme Viteri, Chuqiao Gong, Pascual Lorente, Antonio Fabregat, Konstantinos Sidiropoulos, et al. 2020. “The Reactome Pathway Knowledgebase.” Nucleic Acids Research 48 (D1): D498–503.

39. Johnson, Brett E., Allison L. Creason, Jayne M. Stommel, Jamie M. Keck, Swapnil Parmar, Courtney B. Betts, Aurora Blucher, et al. 2022. “An Omic and Multidimensional Spatial Atlas from Serial Biopsies of an Evolving Metastatic Breast Cancer.” *Cell Reports*. Medicine 3 (2): 100525.

40. Kanehisa, Minoru, Yoko Sato, Masayuki Kawashima, Miho Furumichi, and Mao Tanabe. 2016. “KEGG as a Reference Resource for Gene and Protein Annotation.” Nucleic Acids Research 44 (D1): D457–62.

41. Keller, Mark S., Ilan Gold, Chuck McCallum, Trevor Manz, Peter V. Kharchenko, and Nils Gehlenborg. 2021. “Vitessce: A Framework for Integrative Visualization of Multi-Modal and Spatially-Resolved Single-Cell Data.” 10.31219/osf.io/y8thv.

42. Kettner, Nicole M., Smruthi Vijayaraghavan, Merih Guray Durak, Tuyen Bui, Mehrnoosh Kohansal, Min Jin Ha, Bin Liu, et al. 2019. “Combined Inhibition of STAT3 and DNA Repair in Palbociclib-Resistant ER-Positive Breast Cancer.” Clinical Cancer Research: An Official Journal of the American Association for Cancer Research 25 (13): 3996–4013.

43. Knudsen, Erik S., Vishnu Kumarasamy, Amanda Ruiz, Jared Sivinski, Sejin Chung, Adam Grant, Paris Vail, et al. 2019. “Cell Cycle Plasticity Driven by MTOR Signaling: Integral Resistance to CDK4/6 Inhibition in Patient-Derived Models of Pancreatic Cancer.” Oncogene 38 (18): 3355–70.

44. Kravtsov, Dmitriy S., Amy K. Erbe, Paul M. Sondel, and Alexander L. Rakhmilevich. 2022. “Roles of CD4+ T Cells as Mediators of Antitumor Immunity.” Frontiers in Immunology 13 (September): 972021.

45. Labrie, Marilyne, Tae-Beom Kim, Zhenlin Ju, Sanghoon Lee, Wei Zhao, Yong Fang, Yiling Lu, et al. 2019. “Adaptive Responses in a PARP Inhibitor Window of Opportunity Trial Illustrate Limited Functional Interlesional Heterogeneity and Potential Combination Therapy Options.” Oncotarget 10 (37): 3533–46.

46. Lachmann, Alexander, Federico M. Giorgi, Gonzalo Lopez, and Andrea Califano. 2016. “ARACNe-AP: Gene Network Reverse Engineering through Adaptive Partitioning Inference of Mutual Information.” *Bioinformatics (Oxford*, England) 32 (14): 2233–35.

47. Larionova, Irina, Gulnara Tuguzbaeva, Anastasia Ponomaryova, Marina Stakheyeva, Nadezhda Cherdyntseva, Valentin Pavlov, Evgeniy Choinzonov, and Julia Kzhyshkowska. 2020. “Tumor-Associated Macrophages in Human Breast, Colorectal, Lung, Ovarian and Prostate Cancers.” Frontiers in Oncology 10 (October): 566511.

48. Li, Chunxiao, Ping Jiang, Shuhua Wei, Xiaofei Xu, and Junjie Wang. 2020. “Regulatory T Cells in Tumor Microenvironment: New Mechanisms, Potential Therapeutic Strategies and Future Prospects.” Molecular Cancer 19 (1): 116.

49. Li, Heng. 2013. “Aligning Sequence Reads, Clone Sequences and Assembly Contigs with BWA-MEM.” 10.48550/ARXIV.1303.3997.

50. Li, Jing, Yi He, Jing Hao, Ling Ni, and Chen Dong. 2018. “High Levels of Eomes Promote Exhaustion of Anti-Tumor CD8+ T Cells.” Frontiers in Immunology 9 (December): 2981.

51. Li, Zhiqiang, Pedram Razavi, Qing Li, Weiyi Toy, Bo Liu, Christina Ping, Wilson Hsieh, et al. 2018. “Loss of the FAT1 Tumor Suppressor Promotes Resistance to CDK4/6 Inhibitors via the Hippo Pathway.” Cancer Cell 34 (6): 893–905.e8.

52. Liberzon, Arthur, Chet Birger, Helga Thorvaldsdóttir, Mahmoud Ghandi, Jill P. Mesirov, and Pablo Tamayo. 2015. “The Molecular Signatures Database (MSigDB) Hallmark Gene Set Collection.” Cell Systems 1 (6): 417–25.

53. Liberzon, Arthur, Aravind Subramanian, Reid Pinchback, Helga Thorvaldsdóttir, Pablo Tamayo, and Jill P. Mesirov. 2011. “Molecular Signatures Database (MSigDB) 3.0.” *Bioinformatics (Oxford*, England) 27 (12): 1739–40.

54. Lin, Elaine Y., Jiu-Feng Li, Leoid Gnatovskiy, Yan Deng, Liyin Zhu, Dustin A. Grzesik, Hong Qian, Xiao-Nan Xue, and Jeffrey W. Pollard. 2006. “Macrophages Regulate the Angiogenic Switch in a Mouse Model of Breast Cancer.” Cancer Research 66 (23): 11238–46.

55. Lin, Jia-Ren, Benjamin Izar, Shu Wang, Clarence Yapp, Shaolin Mei, Parin M. Shah, Sandro Santagata, and Peter K. Sorger. 2018. “Highly Multiplexed Immunofluorescence Imaging of Human Tissues and Tumors Using T-CyCIF and Conventional Optical Microscopes.” ELife 7 (July). 10.7554/eLife.31657.

56. Liu, Jinlin, Ning Zhang, Qun Li, Weiwei Zhang, Fang Ke, Qibin Leng, Hong Wang, Jinfei Chen, and Honglin Wang. 2011. “Tumor-Associated Macrophages Recruit CCR6+ Regulatory T Cells and Promote the Development of Colorectal Cancer via Enhancing CCL20 Production in Mice.” PloS One 6 (4): e19495.

57. Manz, Trevor, Ilan Gold, Nathan Heath Patterson, Chuck McCallum, Mark S. Keller, Bruce W. Herr 2nd, Katy Börner, Jeffrey M. Spraggins, and Nils Gehlenborg. 2022. “Viv: Multiscale Visualization of High-Resolution Multiplexed Bioimaging Data on the Web.” Nature Methods 19 (5): 515–16.

58. Marciscano, Ariel E., and Niroshana Anandasabapathy. 2021. “The Role of Dendritic Cells in Cancer and Anti-Tumor Immunity.” Seminars in Immunology 52 (101481): 101481.

59. Matthews, Lisa, Gopal Gopinath, Marc Gillespie, Michael Caudy, David Croft, Bernard de Bono, Phani Garapati, et al. 2009. “Reactome Knowledgebase of Human Biological Pathways and Processes.” Nucleic Acids Research 37 (Database issue): D619–22.

60. Medrek, Catharina, Fredrik Pontén, Karin Jirström, and Karin Leandersson. 2012. “The Presence of Tumor Associated Macrophages in Tumor Stroma as a Prognostic Marker for Breast Cancer Patients.” BMC Cancer 12 (1): 306.

61. Michaloglou, Chrysiis, Claire Crafter, Rasmus Siersbaek, Oona Delpuech, Jon O. Curwen, Larissa S. Carnevalli, Anna D. Staniszewska, et al. 2018. “Combined Inhibition of MTOR and CDK4/6 Is Required for Optimal Blockade of E2F Function and Long-Term Growth Inhibition in Estrogen Receptor-Positive Breast Cancer.” Molecular Cancer Therapeutics 17 (5): 908–20.

62. Mitri, Zahi I., Swapnil Parmar, Brett Johnson, Annette Kolodzie, Jamie M. Keck, Max Morris, Alexander R. Guimaraes, et al. 2018. “Implementing a Comprehensive Translational Oncology Platform: From Molecular Testing to Actionability.” Journal of Translational Medicine 16 (1): 358.

63. Mulqueen, Ryan M., Dmitry Pokholok, Brendan L. O’Connell, Casey A. Thornton, Fan Zhang, Brian J. O’Roak, Jason Link, et al. 2021. “High-Content Single-Cell Combinatorial Indexing.” Nature Biotechnology 39 (12): 1574–80.

64. Munhoz, Rodrigo Ramella, and Michael Andrew Postow. 2016. “Recent Advances in Understanding Antitumor Immunity.” F1000Research 5 (October): 2545.

65. Ni, Chao, Liu Yang, Qiuran Xu, Hongjun Yuan, Wei Wang, Wenjie Xia, Dihe Gong, Wei Zhang, and Kun Yu. 2019. “CD68- and CD163-Positive Tumor Infiltrating Macrophages in Non-Metastatic Breast Cancer: A Retrospective Study and Meta-Analysis.” Journal of Cancer 10 (19): 4463–72.

66. Noy, Roy, and Jeffrey W. Pollard. 2014. “Tumor-Associated Macrophages: From Mechanisms to Therapy.” Immunity 41 (1): 49–61.

67. Núñez Abad, Martín, Silvia Calabuig-Fariñas, Miriam Lobo de Mena, Susana Torres-Martínez, Clara García González, José Ángel García García, Vega Iranzo González-Cruz, and Carlos Camps Herrero. 2022. “Programmed Death-Ligand 1 (PD-L1) as Immunotherapy Biomarker in Breast Cancer.” Cancers 14 (2): 307.

68. Occhipinti, Giulia, Emanuela Romagnoli, Matteo Santoni, Alessia Cimadamore, Giulia Sorgentoni, Monia Cecati, Matteo Giulietti, et al. 2020. “Sequential or Concomitant Inhibition of Cyclin-Dependent Kinase 4/6 before MTOR Pathway in Hormone-Positive HER2 Negative Breast Cancer: Biological Insights and Clinical Implications.” Frontiers in Genetics 11 (April): 349.

69. O’Leary, Ben, Rosalind J. Cutts, Yuan Liu, Sarah Hrebien, Xin Huang, Kerry Fenwick, Fabrice André, et al. 2018. “The Genetic Landscape and Clonal Evolution of Breast Cancer Resistance to Palbociclib plus Fulvestrant in the PALOMA-3 Trial.” Cancer Discovery 8 (11): 1390–1403.

70. Park, Yeon Hee, Seock-Ah Im, Kyunghee Park, Ji Wen, Kyung-Hun Lee, Yoon-La Choi, Won-Chul Lee, et al. 2023. “Longitudinal Multi-Omics Study of Palbociclib Resistance in HR-Positive/HER2-Negative Metastatic Breast Cancer.” Genome Medicine 15 (1): 55.

71. Parker, Joel S., Michael Mullins, Maggie C. U. Cheang, Samuel Leung, David Voduc, Tammi Vickery, Sherri Davies, et al. 2009. “Supervised Risk Predictor of Breast Cancer Based on Intrinsic Subtypes.” Journal of Clinical Oncology: Official Journal of the American Society of Clinical Oncology 27 (8): 1160–67.

72. Pernas, Sonia, Sara M. Tolaney, Eric P. Winer, and Shom Goel. 2018. “CDK4/6 Inhibition in Breast Cancer: Current Practice and Future Directions.” Therapeutic Advances in Medical Oncology 10 (July): 1758835918786451.

73. Phillips, Darci, Magdalena Matusiak, Belén Rivero Gutierrez, Salil S. Bhate, Graham L. Barlow, Sizun Jiang, Janos Demeter, et al. 2021. “Immune Cell Topography Predicts Response to PD-1 Blockade in Cutaneous T Cell Lymphoma.” Nature Communications 12 (1): 6726.

74. Poh, Ashleigh R., and Matthias Ernst. 2018. “Targeting Macrophages in Cancer: From Bench to Bedside.” Frontiers in Oncology 8 (March): 49.

75. Prabhakaran, Sangeetha, Victoria T. Rizk, Zhenjun Ma, Chia-Ho Cheng, Anders E. Berglund, Dominico Coppola, Farah Khalil, James J. Mulé, and Hatem H. Soliman. 2017. “Evaluation of Invasive Breast Cancer Samples Using a 12-Chemokine Gene Expression Score: Correlation with Clinical Outcomes.” Breast Cancer Research: BCR 19 (1): 71.

76. Pu, Yunzhou, and Qing Ji. 2022. “Tumor-Associated Macrophages Regulate PD-1/PD-L1 Immunosuppression.” Frontiers in Immunology 13 (May): 874589.

77. Qadir, Abdul S., Paolo Ceppi, Sonia Brockway, Calvin Law, Liang Mu, Nikolai N. Khodarev, Jung Kim, et al. 2017. “CD95/Fas Increases Stemness in Cancer Cells by Inducing a STAT1-Dependent Type I Interferon Response.” Cell Reports 18 (10): 2373–86.

78. Raskov, Hans, Adile Orhan, Jan Pravsgaard Christensen, and Ismail Gögenur. 2021. “Cytotoxic CD8+ T Cells in Cancer and Cancer Immunotherapy.” British Journal of Cancer 124 (2): 359–67.

79. Romano, Gabriele, Pei-Ling Chen, Ping Song, Jennifer L. McQuade, Roger J. Liang, Mingguang Liu, Whijae Roh, et al. 2018. “A Preexisting Rare PIK3CAE545K Subpopulation Confers Clinical Resistance to MEK plus CDK4/6 Inhibition in NRAS Melanoma and Is Dependent on S6K1 Signaling.” Cancer Discovery 8 (5): 556–67.

80. Rozenblatt-Rosen, Orit, Aviv Regev, Philipp Oberdoerffer, Tal Nawy, Anna Hupalowska, Jennifer E. Rood, Orr Ashenberg, et al. 2020. “The Human Tumor Atlas Network: Charting Tumor Transitions across Space and Time at Single-Cell Resolution.” Cell 181 (2): 236–49.

81. Sautès-Fridman, Catherine, Florent Petitprez, Julien Calderaro, and Wolf Herman Fridman. 2019. “Tertiary Lymphoid Structures in the Era of Cancer Immunotherapy.” Nature Reviews. Cancer 19 (6): 307–25.

82. Sawant, Deepali V., Hiroshi Yano, Maria Chikina, Qianxia Zhang, Mengting Liao, Chang Liu, Derrick J. Callahan, et al. 2019. “Adaptive Plasticity of IL-10+ and IL-35+ Treg Cells Cooperatively Promotes Tumor T Cell Exhaustion.” Nature Immunology 20 (6): 724–35.

83. Scheidemann, Erin R., and Ayesha N. Shajahan-Haq. 2021. “Resistance to CDK4/6 Inhibitors in Estrogen Receptor-Positive Breast Cancer.” International Journal of Molecular Sciences 22 (22): 12292.

84. Schmidt, Angelika, Nina Oberle, and Peter H. Krammer. 2012. “Molecular Mechanisms of Treg-Mediated T Cell Suppression.” Frontiers in Immunology 3 (March): 51.

85. Scirocchi, Fabio, Simone Scagnoli, Andrea Botticelli, Alessandra Di Filippo, Chiara Napoletano, Ilaria Grazia Zizzari, Lidia Strigari, et al. 2022. “Immune Effects of CDK4/6 Inhibitors in Patients with HR+/HER2-Metastatic Breast Cancer: Relief from Immunosuppression Is Associated with Clinical Response.” EBioMedicine 79 (104010): 104010.

86. Sinnamon, John R., Kristof A. Torkenczy, Michael W. Linhoff, Sarah A. Vitak, Ryan M. Mulqueen, Hannah A. Pliner, Cole Trapnell, Frank J. Steemers, Gail Mandel, and Andrew C. Adey. 2019. “The Accessible Chromatin Landscape of the Murine Hippocampus at Single-Cell Resolution.” Genome Research 29 (5): 857–69.

87. Spranger, Stefani. 2016. “Mechanisms of Tumor Escape in the Context of the T-Cell-Inflamed and the Non-T-Cell-Inflamed Tumor Microenvironment.” International Immunology 28 (8): 383–91.

88. Subramanian, Aravind, Pablo Tamayo, Vamsi K. Mootha, Sayan Mukherjee, Benjamin L. Ebert, Michael A. Gillette, Amanda Paulovich, et al. 2005. “Gene Set Enrichment Analysis: A Knowledge-Based Approach for Interpreting Genome-Wide Expression Profiles.” Proceedings of the National Academy of Sciences of the United States of America 102 (43): 15545–50.

89. Tamborero, David, Carlota Rubio-Perez, Ferran Muiños, Radhakrishnan Sabarinathan, Josep M. Piulats, Aura Muntasell, Rodrigo Dienstmann, Nuria Lopez-Bigas, and Abel Gonzalez-Perez. 2018. “A Pan-Cancer Landscape of Interactions between Solid Tumors and Infiltrating Immune Cell Populations.” Clinical Cancer Research: An Official Journal of the American Association for Cancer Research 24 (15): 3717– 28.

90. Teh, Jessica L. F., and Andrew E. Aplin. 2019. “Arrested Developments: CDK4/6 Inhibitor Resistance and Alterations in the Tumor Immune Microenvironment.” Clinical Cancer Research: An Official Journal of the American Association for Cancer Research 25 (3): 921–27.

91. Tiainen, Satu, Ritva Tumelius, Kirsi Rilla, Kirsi Hämäläinen, Markku Tammi, Raija Tammi, Veli-Matti Kosma, Sanna Oikari, and Päivi Auvinen. 2015. “High Numbers of Macrophages, Especially M2-like (CD163-Positive), Correlate with Hyaluronan Accumulation and Poor Outcome in Breast Cancer.” Histopathology 66 (6): 873–83.

92. Tibes, Raoul, Yihua Qiu, Yiling Lu, Bryan Hennessy, Michael Andreeff, Gordon B. Mills, and Steven M. Kornblau. 2006. “Reverse Phase Protein Array: Validation of a Novel Proteomic Technology and Utility for Analysis of Primary Leukemia Specimens and Hematopoietic Stem Cells.” Molecular Cancer Therapeutics 5 (10): 2512–21.

93. Trujillo, Jonathan A., Randy F. Sweis, Riyue Bao, and Jason J. Luke. 2018. “T Cell-Inflamed versus Non-T Cell-Inflamed Tumors: A Conceptual Framework for Cancer Immunotherapy Drug Development and Combination Therapy Selection.” Cancer Immunology Research 6 (9): 990–1000.

94. Tu, Daoyuan, Jin Dou, Mingkao Wang, Haiwen Zhuang, and Xiaoyu Zhang. 2021. “M2 Macrophages Contribute to Cell Proliferation and Migration of Breast Cancer.” Cell Biology International 45 (4): 831– 38.

95. Ubhi, Tajinder, and Grant W. Brown. 2019. “Exploiting DNA Replication Stress for Cancer Treatment.” Cancer Research 79 (8): 1730–39.

96. Villarino, Alejandro V., Yuka Kanno, John R. Ferdinand, and John J. O’Shea. 2015. “Mechanisms of Jak/STAT Signaling in Immunity and Disease.” The Journal of Immunology 194 (1): 21–27.

97. Waks, Adrienne G., and Eric P. Winer. 2019. “Breast Cancer Treatment: A Review.” JAMA: The Journal of the American Medical Association 321 (3): 288–300.

98. Wander, Seth A., Ofir Cohen, Xueqian Gong, Gabriela N. Johnson, Jorge E. Buendia-Buendia, Maxwell R. Lloyd, Dewey Kim, et al. 2020. “The Genomic Landscape of Intrinsic and Acquired Resistance to Cyclin-Dependent Kinase 4/6 Inhibitors in Patients with Hormone Receptor-Positive Metastatic Breast Cancer.” Cancer Discovery 10 (8): 1174–93.

99. Wang, Shixiang, Zaoke He, Xuan Wang, Huimin Li, and Xue-Song Liu. 2019. “Antigen Presentation and Tumor Immunogenicity in Cancer Immunotherapy Response Prediction.” ELife 8 (November). 10.7554/eLife.49020.

100. Wickham, Hadley. 2009. Ggplot2. New York, NY: Springer New York.

101. Xu, Xia-Qing, Xiao-Hui Pan, Ting-Ting Wang, Jian Wang, Bo Yang, Qiao-Jun He, and Ling Ding. 2021. “Intrinsic and Acquired Resistance to CDK4/6 Inhibitors and Potential Overcoming Strategies.” Acta Pharmacologica Sinica 42 (2): 171–78.

102. Yang, C., Z. Li, T. Bhatt, M. Dickler, D. Giri, M. Scaltriti, J. Baselga, N. Rosen, and S. Chandarlapaty. 2017. “Acquired CDK6 Amplification Promotes Breast Cancer Resistance to CDK4/6 Inhibitors and Loss of ER Signaling and Dependence.” Oncogene 36 (16): 2255–64.

103. Yoshida, Akihiro, Yiwen Bu, Shuo Qie, John Wrangle, E. Ramsay Camp, E. Starr Hazard, Gary Hardiman, Renée de Leeuw, Karen E. Knudsen, and J. Alan Diehl. 2019. “SLC36A1-MTORC1 Signaling Drives Acquired Resistance to CDK4/6 Inhibitors.” Science Advances 5 (9): eaax6352.

104. Zaidi, M. Raza. 2019. “The Interferon-Gamma Paradox in Cancer.” Journal of Interferon & Cytokine Research: The Official Journal of the International Society for Interferon and Cytokine Research 39 (1): 30–38.

105. Zhang, Hao, Lin Liu, Jinbo Liu, Pengyuan Dang, Shengyun Hu, Weitang Yuan, Zhenqiang Sun, Yang Liu, and Chengzeng Wang. 2023. “Roles of Tumor-Associated Macrophages in Anti-PD-1/PD-L1 Immunotherapy for Solid Cancers.” Molecular Cancer 22 (1): 58.

106. Zhang, Xiaohui, Yuanyuan Zeng, Qiuxia Qu, Jianjie Zhu, Zeyi Liu, Weiwei Ning, Hui Zeng, et al. 2017. “PD-L1 Induced by IFN-γ from Tumor-Associated Macrophages via the JAK/STAT3 and PI3K/AKT Signaling Pathways Promoted Progression of Lung Cancer.” International Journal of Clinical Oncology 22 (6): 1026–33.

107. Zhu, Zhou, Nicholas C. Turner, Sherene Loi, Fabrice André, Miguel Martin, Véronique Diéras, Karen A. Gelmon, et al. 2022. “Comparative Biomarker Analysis of PALOMA-2/3 Trials for Palbociclib.” Npj Precision Oncology 6 (1): 56.

108. Zou, Zhilin, Tao Tao, Hongmei Li, and Xiao Zhu. 2020. “MTOR Signaling Pathway and MTOR Inhibitors in Cancer: Progress and Challenges.” Cell & Bioscience 10 (1): 31.

109. Zou, Zijuan, Hongfen Lin, Mengsen Li, and Bo Lin. 2023. “Tumor-Associated Macrophage Polarization in the Inflammatory Tumor Microenvironment.” Frontiers in Oncology 13 (February): 1103149.

