## Extended Data Figures for "Longitudinal and multimodal auditing of tumor adaptation to CDK4/6 inhibitors in HR+ metastatic breast cancers"

### **Extended Data Figure 1:** Genomic Landscape of HR+ Metastatic Breast Cancer on CDK4/6i Therapy

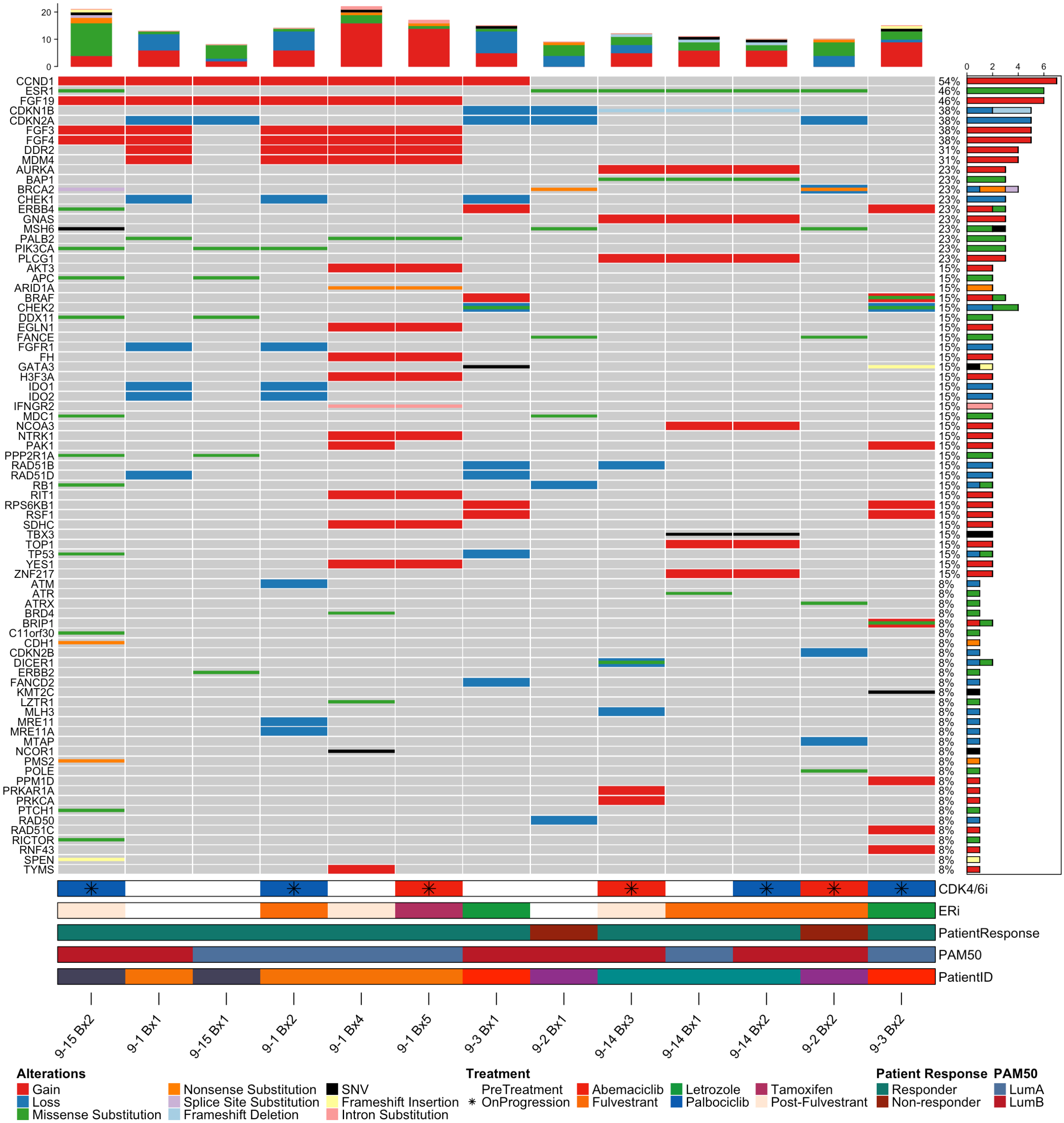

**Extended Data Figure 2:** Gene Set Enrichment Activity of HR+ Metastatic Breast Cancer on CDK4/6i Therapy

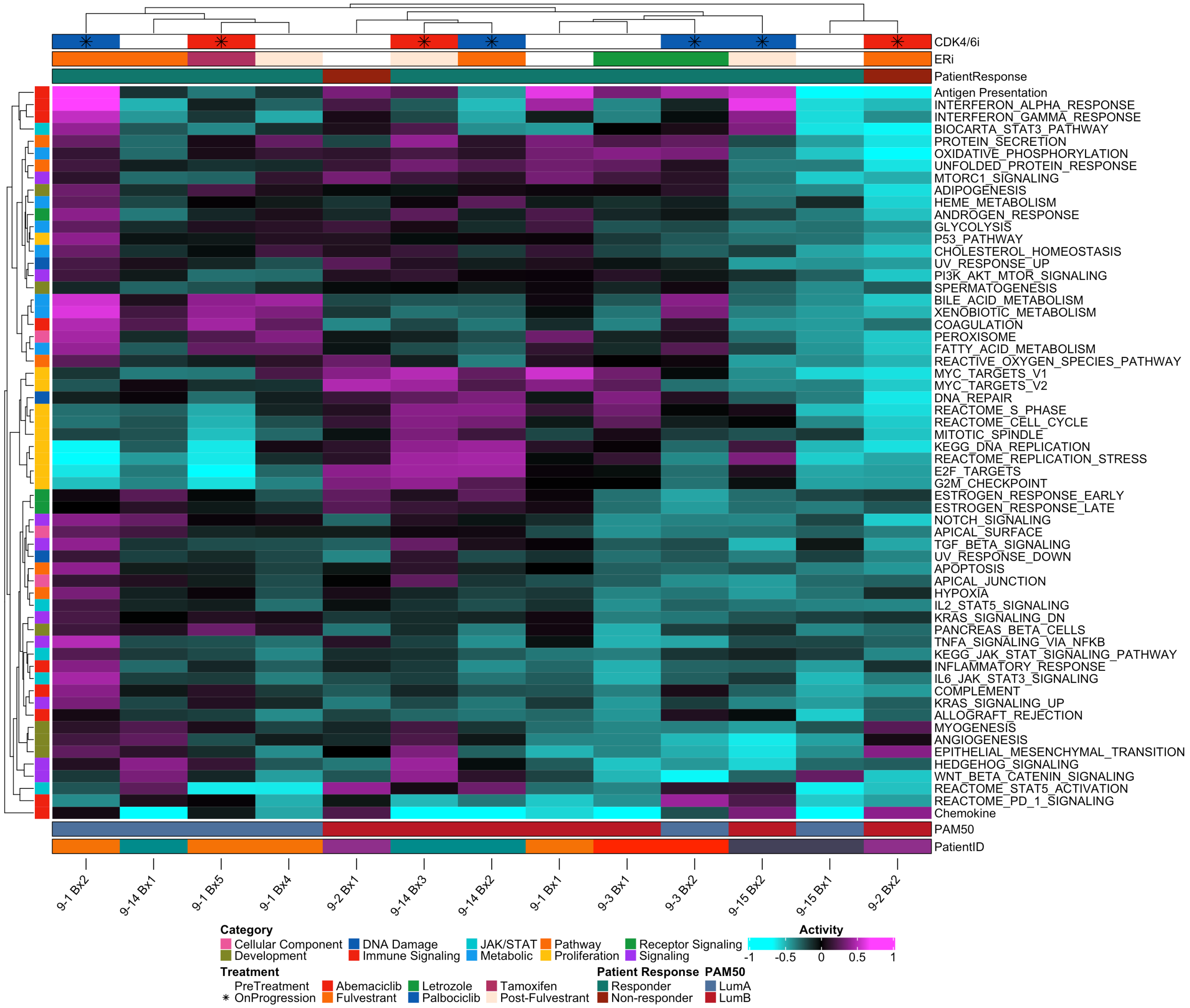

**Extended Data Figure 3:** Gene Set Enrichment Activity of Malignant Cell Pathways During CDK4/6i in External Cohort of HR+ Metastatic Breast Cancers

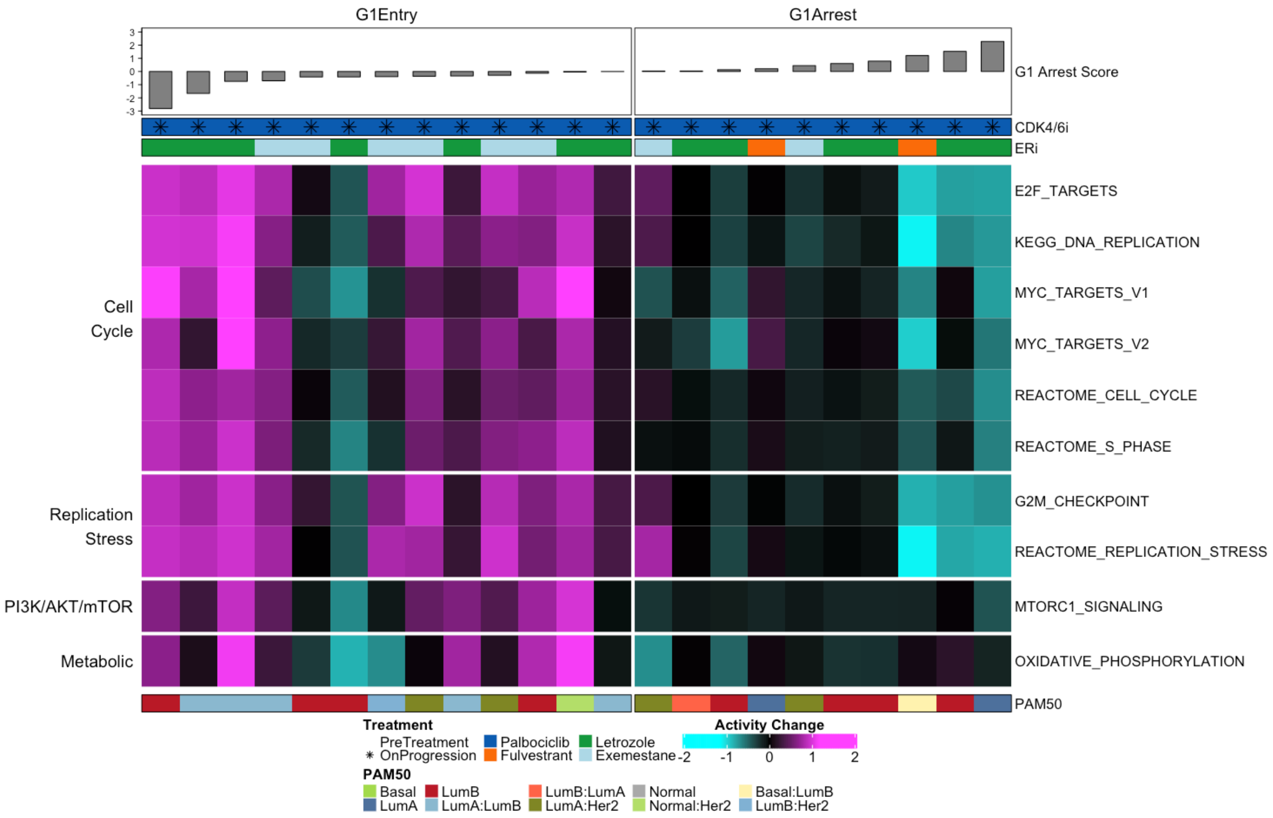

**Extended Data Figure 4:** Transcriptional profiling of multiple cohorts reveals distinct subgroups of malignant cell and TIME adaptations to CDK4/6i therapy

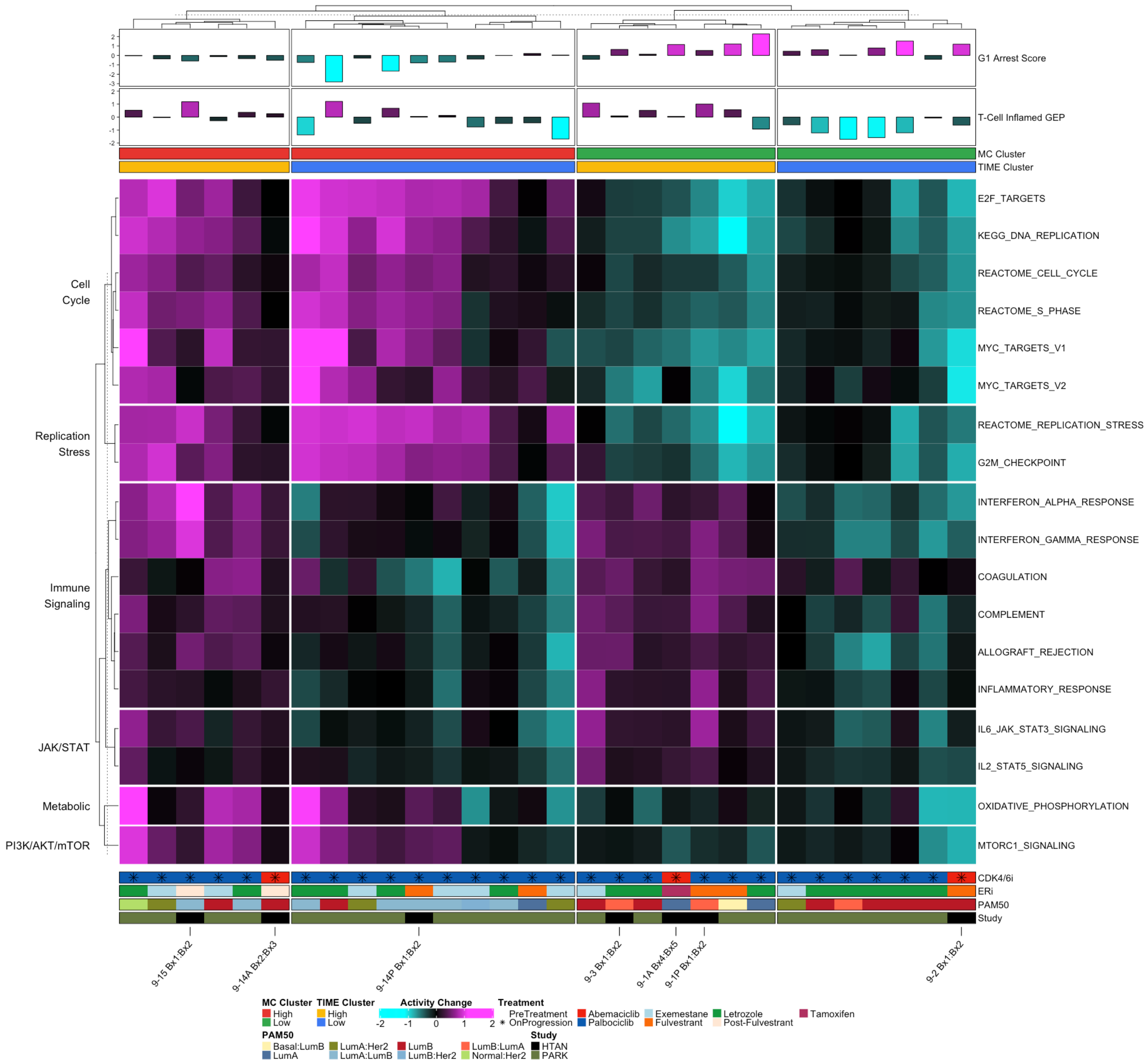

**Extended Data Figure 5:** Master Regulator Profiles of HR+ Metastatic Breast Cancer Cohort

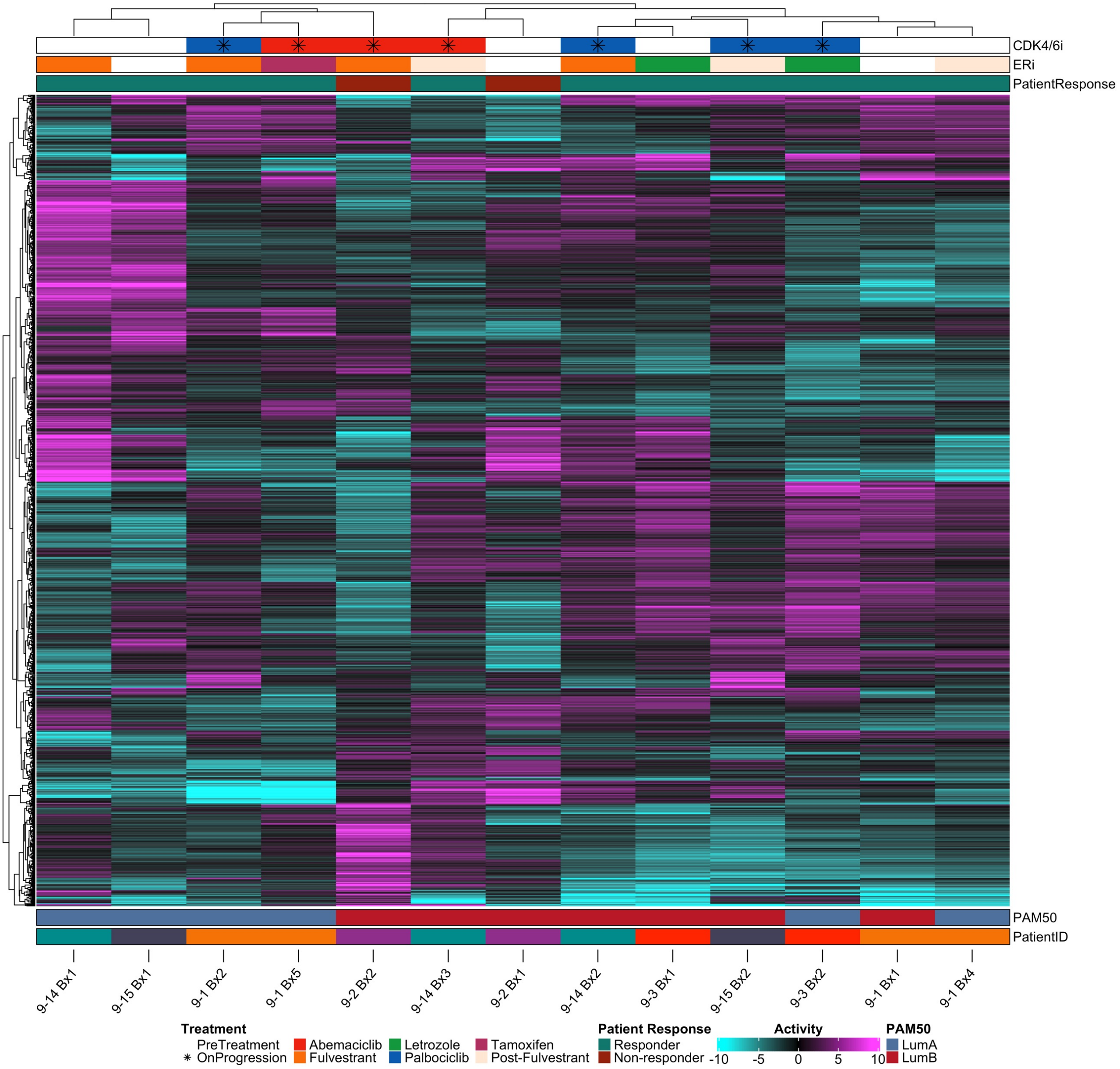

**Extended Data Figure 6:** Distribution of Per-cell Chromatin Accessibility Pathway Enrichment from sciATAC-seq

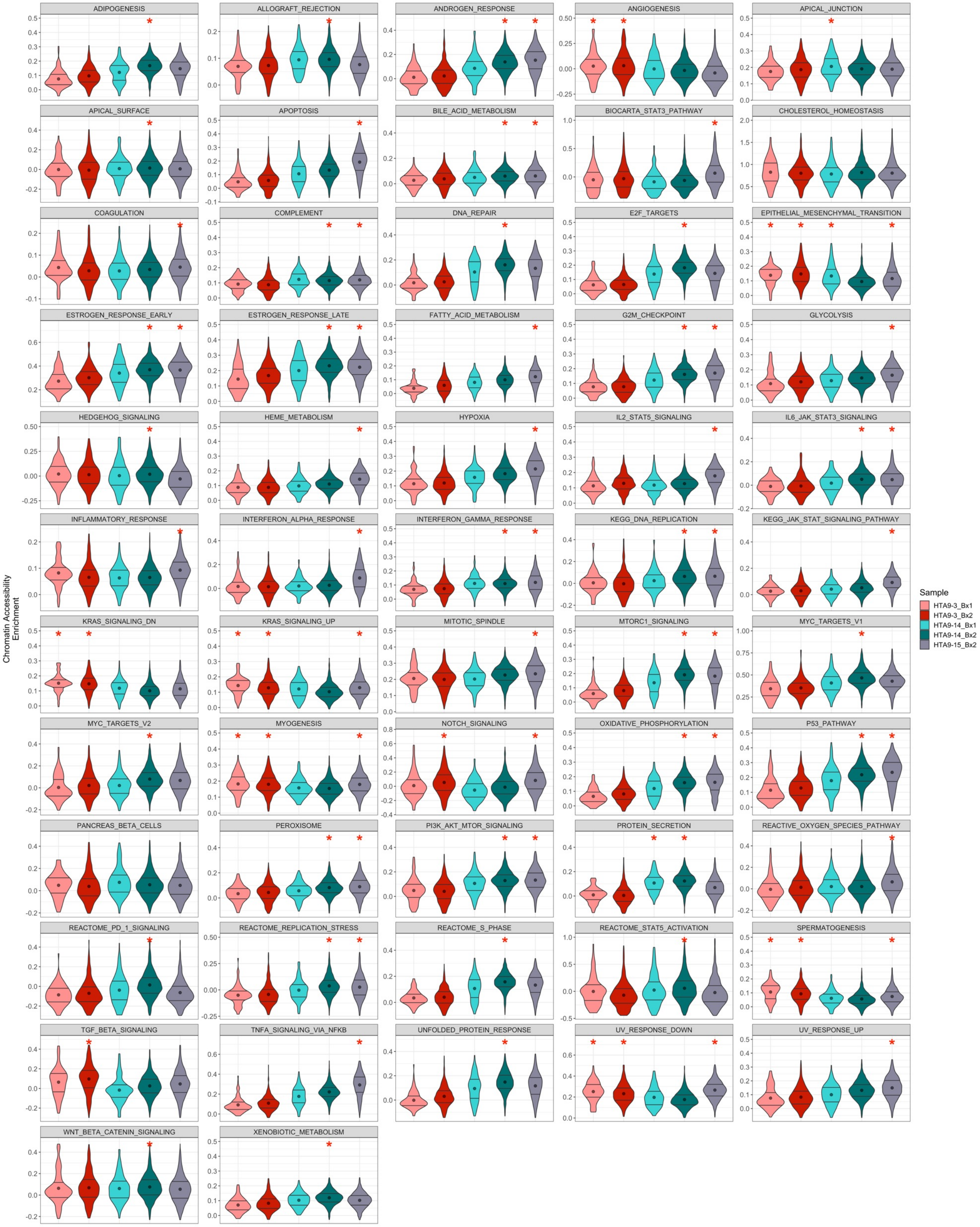

**Extended Data Figure 7:** sciATAC-seq Sample Counts and Cell Type Composition

a

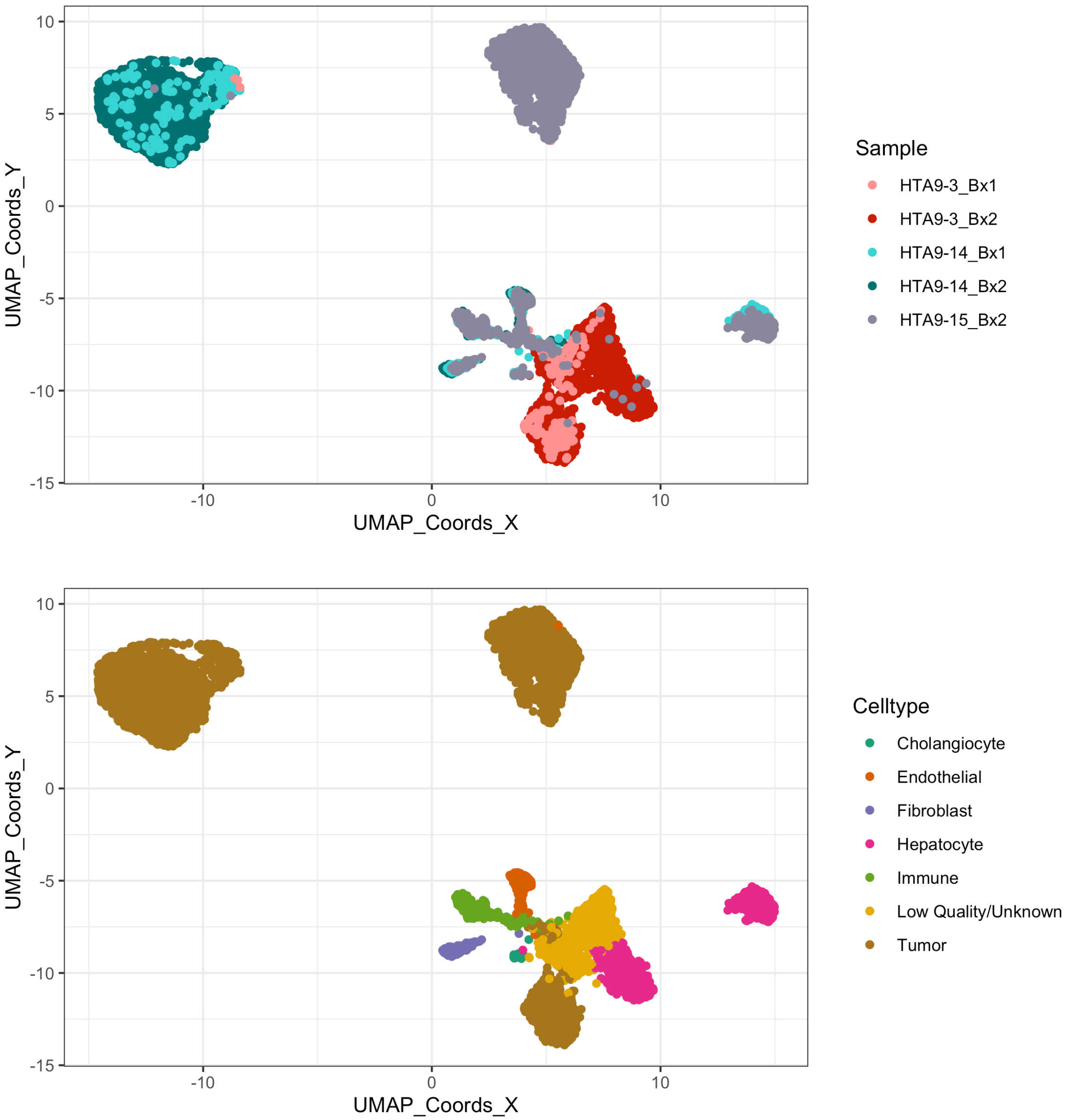


b

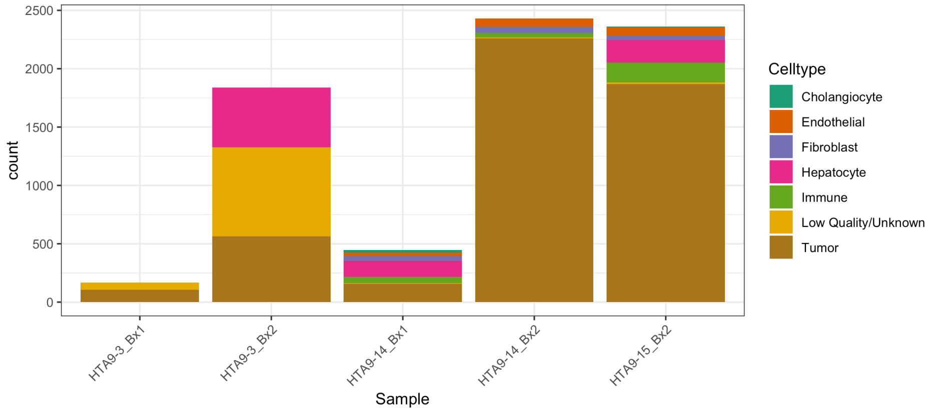

**Extended Data Fig. 8: Immune cell activity across metastatic tumor sites in HTAN and Park et al. 2023 datasets**

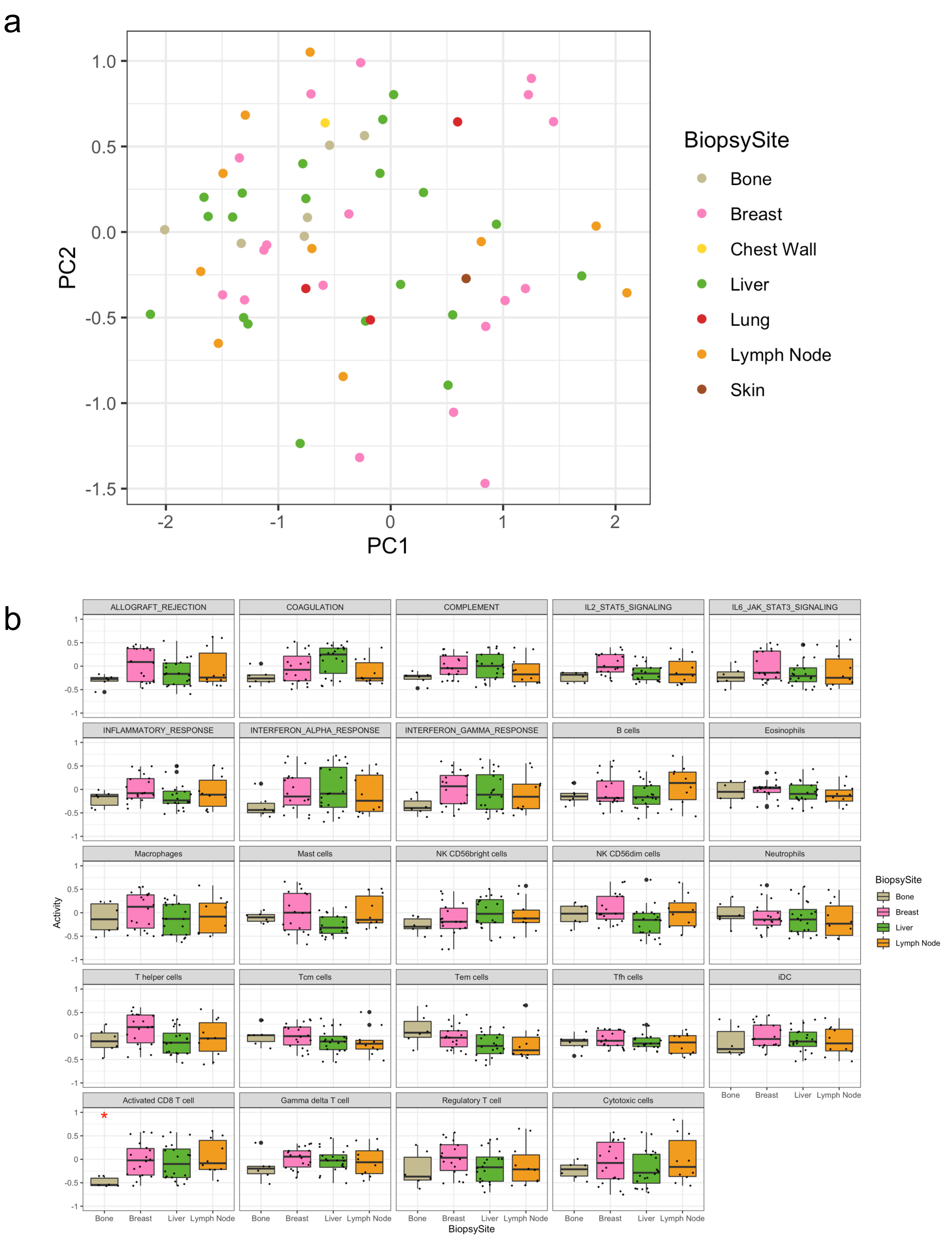

**Extended Data Figure 9:** Immune cell activity of HTAN and Park cases during CDK4/6i

a

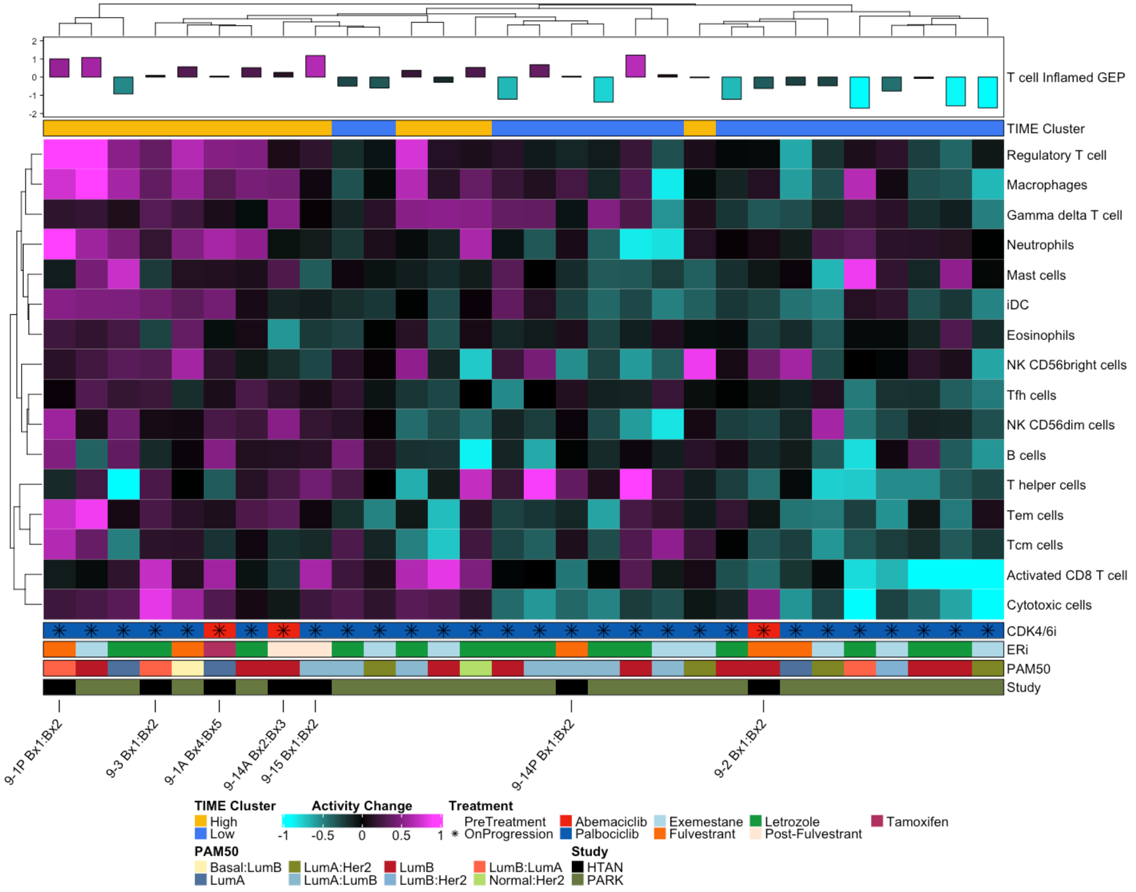


b

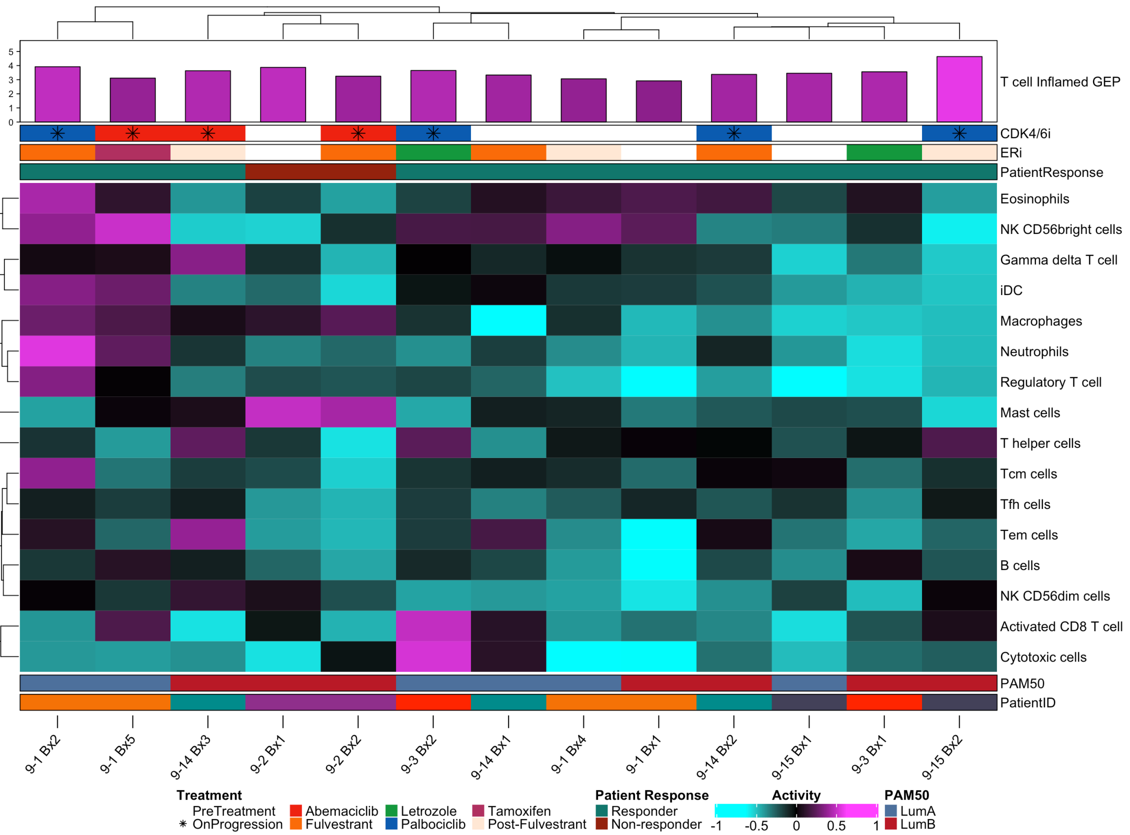

**Extended Data Figure 10:** Proteomic Pathway Activity Profiles of HR+ Metastatic Breast Cancer Cohort

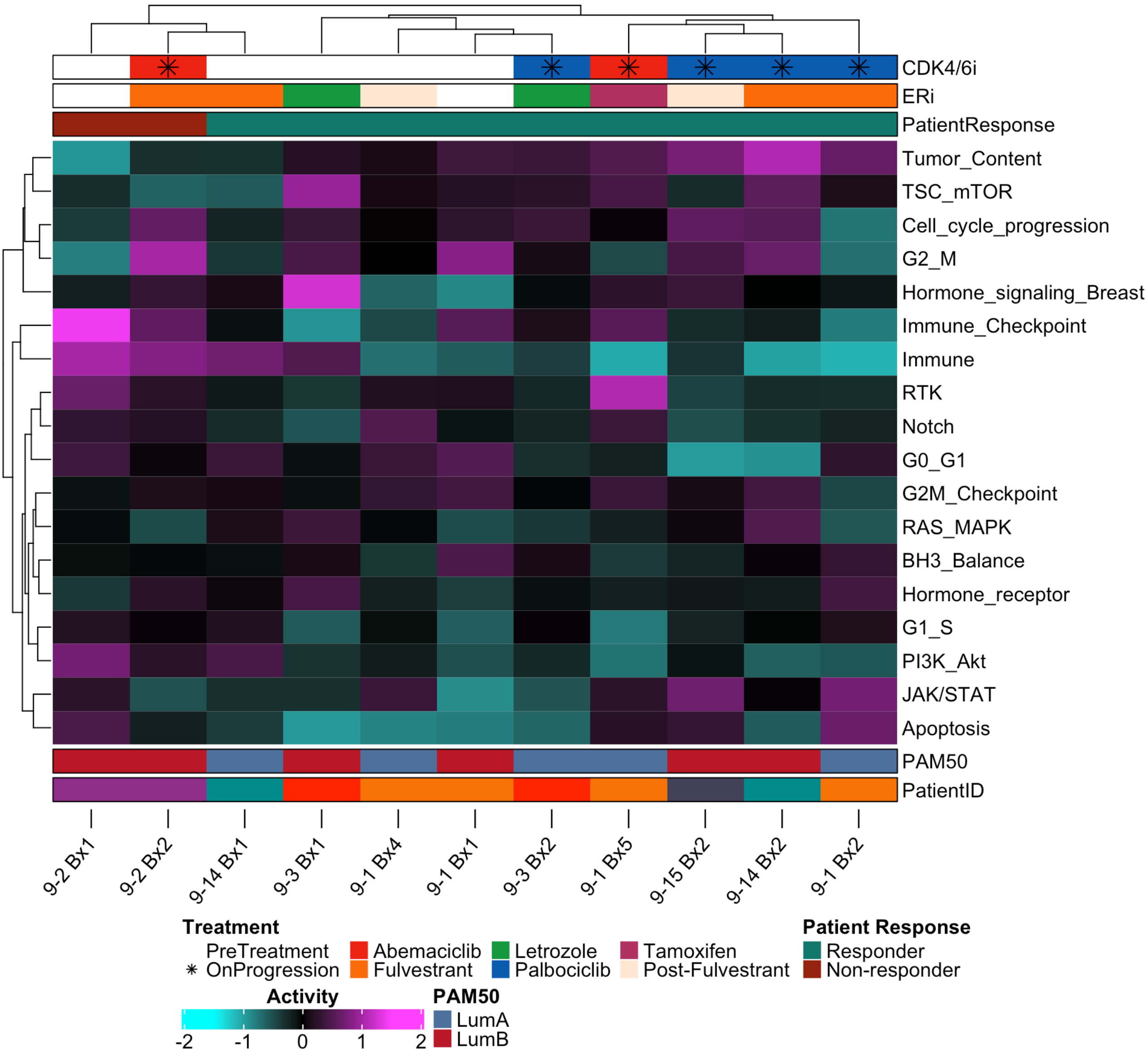
