## Supplemental Tables and Figures for "Longitudinal and multimodal auditing of tumor adaptation to CDK4/6 inhibitors in HR+ metastatic breast cancers"

**Supplemental Figure S1.**

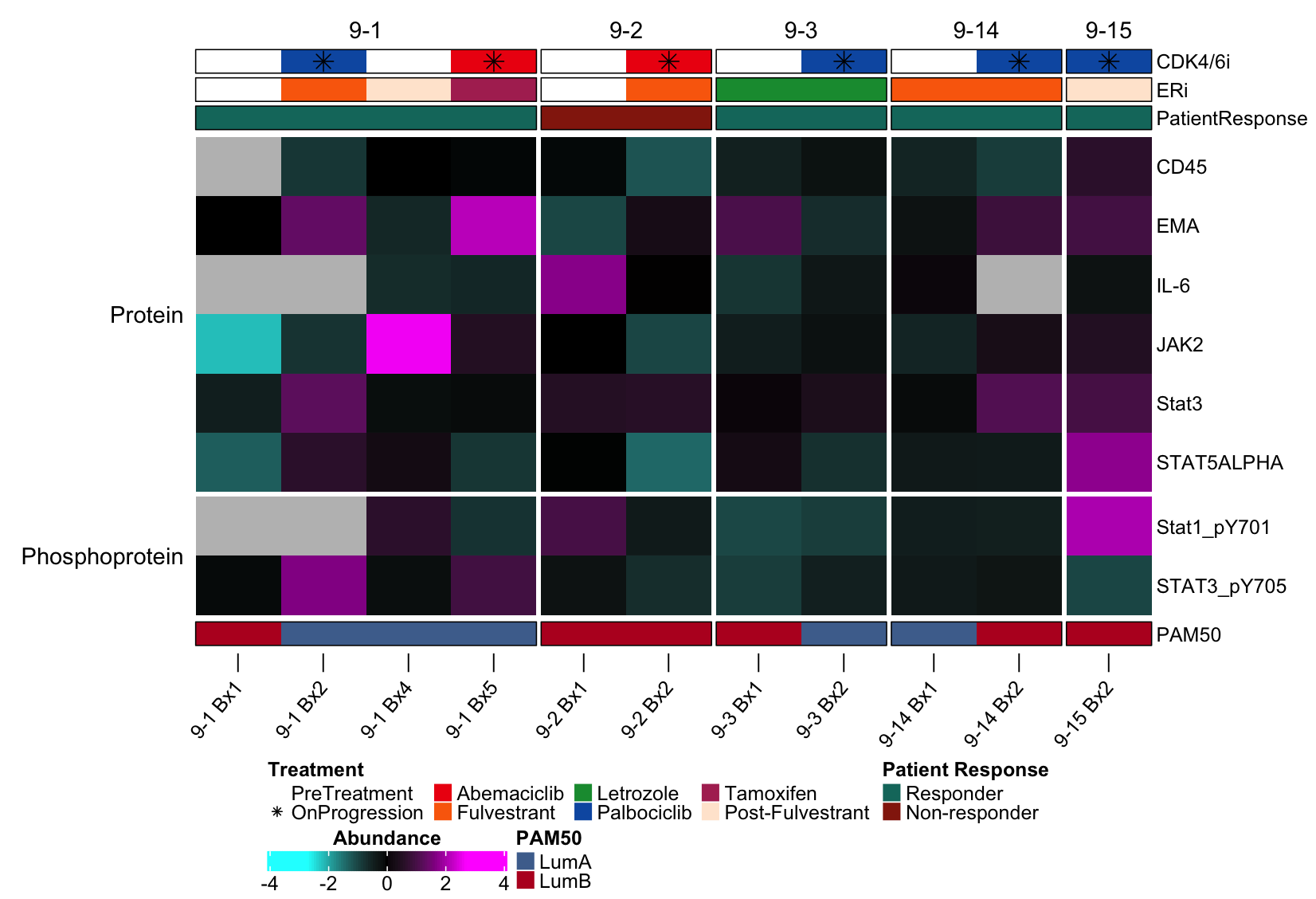

**Supplemental Figure S1**. Heatmap of protein and phosphoprotein changes during CDK4/6i therapy and progression. Heatmap depicts the relative change of markers during therapy using four measurements protein abundance and phosphoprotein levels. Relevant Immune and Jak/Stat were selected for inclusion. Each value is computed by first scaling and mean centering the data relative to a background HR+ cohort and for each pair subtracting the pre-treatment biopsy from the on-progression biopsy.

**Supplemental Figure S2.**

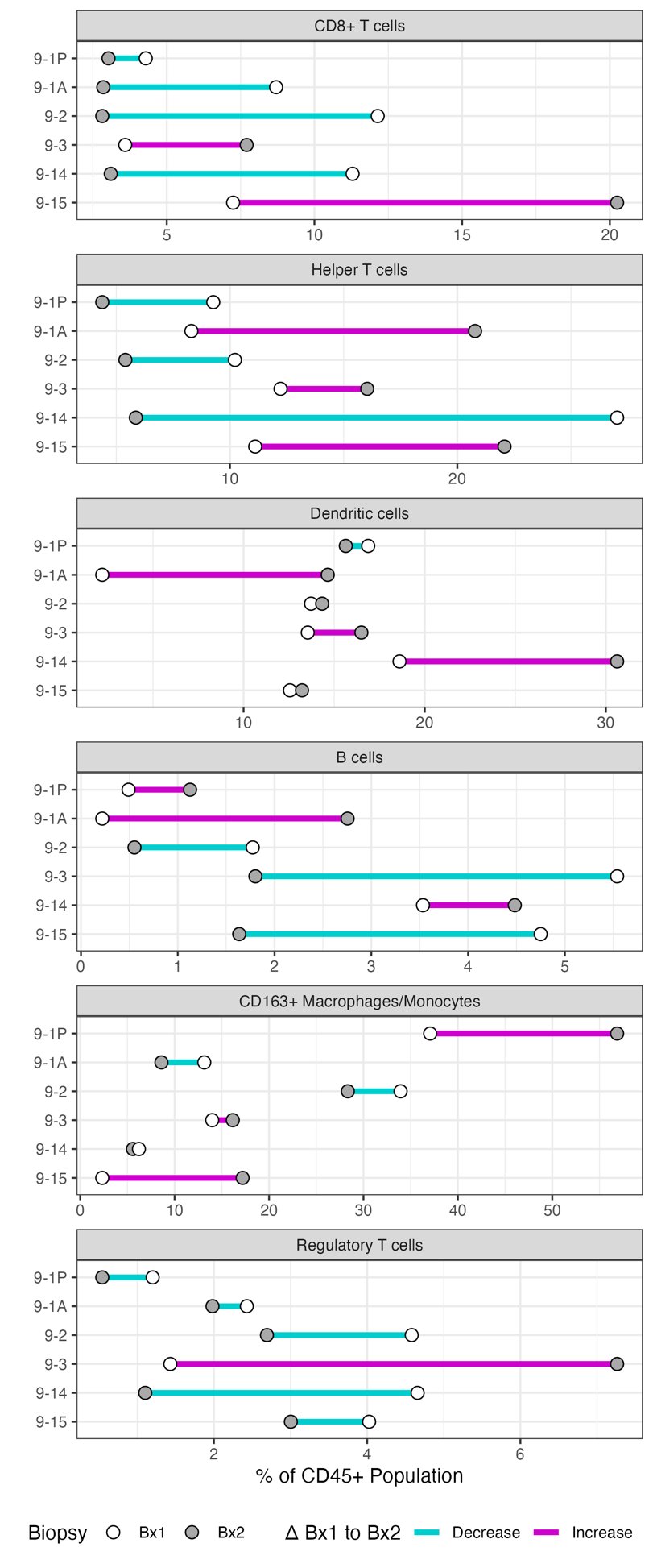

**Supplemental Figure S2**. Cell proportions reported for all CD45^+^ immune cells and individual immune cell type populations across pre-treatment (Bx1, white) and on-progression (Bx2, dark grey) biopsies for all biopsy pairs using mIHC. Line color connecting the pre-treatment and on-progression biopsy points represents direction of change from Bx1 to Bx2.

**Supplemental Table 1.**

|  | AR | BCL2 | ER | GATA3 | HER2 | Ki67 | PDL1 | PR |
| --- | --- | --- | --- | --- | --- | --- | --- | --- |
| 9-1 Bx1 | NA | NA | Positive | NA | Negative | Test Value Reported | NA | Negative |
| 9-1 Bx2 | Positive | Positive | Positive | NA | Negative | Test Value Reported | Test Value Reported | Negative |
| 9-1 Bx4 | Positive | Test Value Reported | Positive | Positive | Negative | Test Value Reported | Negative | Negative |
| 9-1 Bx5 | Positive | Test Value Reported | Positive | Positive | Negative | Test Value Reported | Negative | Negative |
| 9-2 Bx1 | Positive | Positive | Positive | NA | Negative | Test Value Reported | Negative | Positive |
| 9-2 Bx2 | Positive | Positive | Positive | NA | Negative | Test Value Reported | Negative | Positive |
| 9-3 Bx1 | Positive | Positive | Positive | Positive | Equivocal | Test Value Reported | Negative | Negative |
| 9-3 Bx2 | Positive | Positive | Positive | Positive | Negative | Test Value Reported | Negative | Negative |
| 9-3 Bx3 | Positive | Positive | Positive | Positive | Equivocal | Test Value Reported | Negative | Negative |
| 9-14 Bx1 | Positive | Positive | Positive | Positive | Negative | Test Value Reported | Negative | Positive |
| 9-14 Bx2 | Positive | Test Value Reported | Positive | Positive | Equivocal | Test Value Reported | Negative | Positive |
| 9-15 Bx1 | NA | NA | Positive | NA | Negative | Test Value Reported | Negative | Negative |
| 9-15 Bx2 | Positive | Test Value Reported | Positive | Positive | Negative | Test Value Reported | Negative | Negative |
| 9-15 Bx3 | Positive | Test Value Reported | Positive | Positive | Negative | Test Value Reported | NA | NA |

**Supplemental Table 2.**

| **Pathway** | **Avg-G1Arrest** | **Avg-G1Entry** | **FDR** |
| --- | --- | --- | --- |
| E2F_TARGETS | -0.26695 | 0.62027 | 0.00121 |
| G2M_CHECKPOINT | -0.25742 | 0.56652 | 0.00121 |
| REACTOME_CELL_CYCLE | -0.17712 | 0.40951 | 0.00157 |
| KEGG_DNA_REPLICATION | -0.30767 | 0.54574 | 0.00160 |
| MTORC1_SIGNALING | -0.16128 | 0.34945 | 0.00172 |
| MYC_TARGETS_V2 | -0.24419 | 0.44421 | 0.00191 |
| REACTOME_REPLICATION_STRESS | -0.23513 | 0.57390 | 0.00216 |
| MYC_TARGETS_V1 | -0.24354 | 0.41072 | 0.00809 |
| REACTOME_S_PHASE | -0.17034 | 0.40201 | 0.01344 |
| OXIDATIVE_PHOSPHORYLATION | -0.15416 | 0.24188 | 0.07686 |

**Supplemental Table 3.**

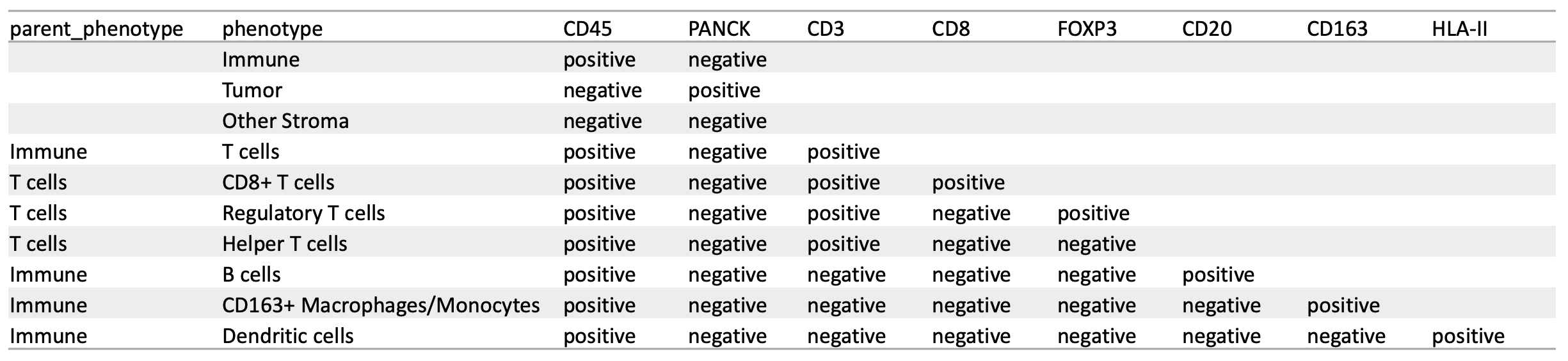

**Supplemental Table 4.**

| Target Name | Antibody Role | Antibody Name | RRID identifier | Fluorophore | Clone | Lot | Vendor | Catalog Number | Assay |
| --- | --- | --- | --- | --- | --- | --- | --- | --- | --- |
| CK5 |  |  |  | 488 | EP1601Y |  | abcam | ab19894 | cycIF |
| CK7 |  | Anti-Cytokeratin 7 |  | 488 | EPR1618Y |  | abcam | ab203434 | cycIF |
| CK8 |  | Anti-Cytokeratin 8 | AB_2864346 | 488 | EP1628Y |  | abcam | ab192467 | cycIF |
| CK14 |  | Anti-Cytokeratin 14 | AB_306091 | 555 | LL002 |  | abcam | ab7800 | cycIF |
| CK17 |  | Anti-Cytokeratin 17 | AB_2889195 | 488 | EP1623 |  | abcam | ab185032 | cycIF |
| CK19 |  | anti-Cytokeratin 19 | AB_439773 | 750 | A53-B/A2 |  | BioLegend | 628502 | cycIF |
| PanCK |  | Pan Cytokeratin | AB_1834350 | 488 | AE1/AE3 |  | Thermofisher | 53-9003-82 | cycIF |
| MUC1 |  | Anti-MUC1 |  | 488 | EPR1023 |  | abcam | ab196443 | cycIF |
| BMP2 |  | Anti-BMP2 |  | 750 | EPR20807 |  | abcam | ab225898 | cycIF |
| TFF1 |  | Anti-Estrogen Inducible Protein pS2 | | 488 | EPR3972 |  | abcam | ab200799 | cycIF |
| TUBB3 |  | Anti-beta III Tubulin |  | 488 | 2G-10 |  | abcam | ab195879 | cycIF |
| SYP |  | Anti-Synaptophysin | AB_2864347 | 555 | YE269 |  | abcam | ab206870 | cycIF |
| CGA |  | Anti-Chromogranin A | | 750 | EP1030Y |  | abcam | ab215276 | cycIF |
| GATA3 |  | Anti-GATA3 | AB_2889199 | 555 | EPR16651 |  | abcam | ab210672 | cycIF |
| Ecad |  | Anti-E Cadherin | AB_2910587 | 750 | EP700Y |  | abcam | ab201499 | cycIF |
| CD44 |  | Anti-CD44 | AB_2889192 | 750 | EPR1013Y |  | abcam | ab194988 | cycIF |
| ER |  | Anti-Estrogen Receptor alpha | AB_2728817 | 647 | EPR4097 |  | abcam | ab205851 | cycIF |
| PR |  | Anti-Progesterone Receptor | | 647 | SP2 |  | abcam | ab267524 | cycIF |
| AR |  | Anti-Androgen Receptor | AB_2922447 | 555 | EPR1535(2) |  | abcam | ab275124 | cycIF |
| EGFR |  | EGF Receptor | AB_10694337 | 555 | D38B1 |  | CST | 5108 | cycIF |
| HER2 |  | Neu (3B5) | AB_627996 | 555 | 3B5 |  | SCBT | sc-33684 | cycIF |
| pERK |  | Phospho-p44/42 MAPK (Erk1/2) (Thr202/Tyr204) | AB_2798131 | 647 | Erk1/2 |  | CST | 13148 | cycIF |
| pAkt |  | Phospho-Akt (Ser473) | AB_916029 | 647 | EPR9701(B) |  | CST | 4075 | cycIF |
| pMYC |  |  |  | 647 | 12B4 |  | Sears Lab | NA | cycIF |
| CD31 |  | Anti-CD31 | AB_2857973 | 647 | EPR3094 |  | abcam | ab218582 | cycIF |
| CAV1 |  |  |  | 488 | EPR15554 |  | abcam | ab225380 | cycIF |
| aSMA |  | alpha-Actin | AB_262054 | 488 | 1A4 |  | Santa Cruz | sc-32251 | cycIF |
| Vimentin | | Vimentin | AB_10829352 | 488 | D21H3 |  | CST | 9854 | cycIF |
| CD90 |  | Anti-CD90 / Thy1 | AB_2889264 | 555 | EPR3133 |  | abcam | ab202511 | cycIF |
| COL(IV) |  | Collagen IV Monoclonal | AB_10853027 | 647 | 1042 |  | Thermofisher | 51-9871-82 | cycIF |
| COL(I) |  | Recombinant Anti-Collagen I | AB_2909621 | 750 | EPR7785 |  | abcam | ab215969 | cycIF |
| H3K27 |  | Tri-Methyl-Histone H3 (Lys27) | AB_2797612 | 488 | C36B11 |  | CST | 5499 | cycIF |
| H3K4 |  | Tri-Methyl-Histone H3 (Lys4) | AB_2797779 | 647 | C42D8 |  | CST | 11960 | cycIF |
| CD45 |  | Anti-CD45 |  | 647 | EP322Y |  | abcam | ab200317 | cycIF |
| CD3 |  | Anti-CD31 | AB_2857973 | 750 | EPR3094 |  | abcam | ab218582 | cycIF |
| CD20 |  | Anti-CD20 |  | 750 | EP459Y |  | abcam | ab198941 | cycIF |
| CD8 |  | Anti-CD8 alpha |  | 555 | C8/468 |  | abcam | ab213017 | cycIF |
| CD4 |  | Anti-CD4 | AB_2923526 | 547 | EPR6855 |  | abcam | ab196147 | cycIF |
| CD68 |  | anti-CD68 | AB_2616797 | 555 | KP1 |  | BioLegend | 916104 | cycIF |
| CSF1R |  | Anti-CSF-1-R |  | 750 | SP211 |  | abcam | ab240265 | cycIF |
| PD1 |  |  |  | 647 | EPR4877(2) |  | abcam | ab2011825 | cycIF |
| FoxP3 |  | Purified anti-human FOXP3 | AB_430881 | 750 | 206D |  | BioLegend | 320102 | cycIF |
| PDL1 |  | Anti-PD-L1 | AB_2728794 | 555 | 28-8 |  | abcam | ab213358 | cycIF |
| GRNZB |  | Anti-Granzyme B | AB_2910576 | 750 | EPR20129 |  | abcam | ab219803 | cycIF |
| PDPN |  | Purified anti-Podoplanin | AB_2565820 | 555 | D2-40 |  | BioLegend | 916606 | cycIF |
| HIF1a |  | Anti-HIF-1 alpha | AB_2923033 | 647 | EP1215Y |  | abcam | ab190569 | cycIF |
| Glut1 |  | Anti-Glucose Transporter GLUT1 | AB_2714026 | 555 | EPR3915 |  | abcam | ab195359 | cycIF |
| CoxIV |  | COX IV | AB_11178794 | 555 | 3E11 |  | CST | 8693 | cycIF |
| pS6RP |  | Phospho-S6 Ribosomal Protein (Ser235/236) | AB_10693792 | 555 | D57 |  | CST | 3985 | cycIF |
| Ki67 |  | Ki67 | AB_2728830 | 647 | D3B5 |  | CST | 12075 | cycIF |
| PCNA |  | PCNA | AB_11178664 | 488 | PC10 |  | CST | 8580 | cycIF |
| pHH3 |  | Anti-Histone H3, phospho (Ser10) | AB_10695860 | 488 | D2C8 |  | CST | 3465 | cycIF |
| pRB |  | Anti-Rb |  | 647 | EPR17732 |  | abcam | ab215947 | cycIF |
| CCND1 |  | Anti-Cyclin D1 | AB_2890216 | 555 | ERP2241 |  | abcam | ab203448 | cycIF |
| p63 |  | Anti-p63 |  | 647 | EPR5701 |  | abcam | ab246728 | cycIF |
| 53BP1 |  | Anti-53BP1 |  | 750 | EPR2172(2) |  | abcam | ab222232 | cycIF |
| MSH6 |  | Anti-MSH6 | AB_2889204 | 647 | EPR3945 |  | abcam | ab198334 | cycIF |
| Rad51 |  | Anti-Rad51 |  | 750 | EPR4030(3) |  | abcam | ab221796 | cycIF |
| gH2AX |  | Anti-gamma H2A.X (phospho S139) | | 647 | EP854(2)Y |  | abcam | ab195189 | cycIF |
| pRPA2 |  | RPA [p Ser33] | AB_10002227 | 750 | Poly |  | Novus | NB100-544 | cycIF |
| LaminAC | | Anti-Lamin A/C | AB_10743057 | 750 | 4C11 |  | Sigma | SAB4200236 | cycIF |
| LaminB1 | | Anti-Lamin B1 | AB_2728786 | 488 | EPR8985(B) |  | abcam | ab194106 | cycIF |
| LaminB2 | | Anti-Lamin B2 | AB_2889288 | 555 | EPR9701(B) |  | abcam | ab200427 | cycIF |
| BcI2 |  | Bcl-2 | AB_2799997 | 647 | 124 |  | CST | 82655 | cycIF |
| CC3 |  | Cleaved Caspase-3 (Asp175) | AB_2797708 | 555 | D3E9 |  | CST | 9604 | cycIF |
| cPARP |  | Cleaved PARP (Asp214) | AB_10830735 | 555 | D64E10 |  | CST | 6894 | cycIF |
| DSDNA |  | anti DS DNA |  |  | DSD/958 |  | Abcam | ab215896 | mIHC |
| PD1 |  | anti PD-1 |  |  | NAT105 |  | Abcam | ab52587 | mIHC |
| PDL1 |  | anti PD-L1 |  |  | E1L3N |  | Cell Signaling Technology | 13684 | mIHC |
| HLAII |  | anti HLA II |  |  | WR18 |  | LS Bio | LS-B10162 | mIHC |
| DCLAMP | | anti DCLAMP |  |  | 1010E1.01 |  | Novus Biological | DDX0191P-100 | mIHC |
| CD3 |  | anti CD3 |  |  | SP7 |  | Thermo Fisher | MA1-90582 | mIHC |
| CD45 |  | anti CD45 |  |  | H130 |  | Invitrogen | 14045982 | mIHC |
| CD8 |  | anti CD8 |  |  | C8/144B |  | Invitrogen | MA513473 | mIHC |
| CCR2 |  | anti CCR2 |  |  | 48607 |  | Abcam | ab176390 | mIHC |
| GRZB |  | anti Granzyme B |  |  | EP230 |  | Sigma Aldrich | 262R | mIHC |
| CD20 |  | anti CD20 |  |  | SP32 |  | Abcam | ab64088 | mIHC |
| CD68 |  | anti CD68 |  |  | PG-M1 |  | Abcam | ab783 | mIHC |
| TBET |  | anti T-bet |  |  | D6N8B |  | Cell signaling technology | 13232 | mIHC |
| CD66B |  | anti CD66B |  |  | G10F5 |  | eBioscience | BDB555723 | mIHC |
| CD11C |  | anti CD11c |  |  | EP1347Y |  | Abcam | ab52632 | mIHC |
| CD163 |  | anti CD163 |  |  | 10D6 |  | Thermo Scientific | MA5-11458 | mIHC |
| CD169 |  | anti CD169 |  |  | HSn 7D2 |  | Novus Biological | NB600-534 | mIHC |
| EOMES |  | anti EOMES |  |  | polyclonal |  | Atlas antibodies | HPA028896 | mIHC |
| EOMES |  | anti EOMES |  |  | polyclonal |  | EMD Millipore | AB2283 | mIHC |
| FOXP3 |  | anti FOXP3 |  |  | 236A/E7 |  | Invitrogen | 14477782 | mIHC |
| CD11B |  | anti CD11b |  |  | EPR1334 |  | Abcam | ab133357 | mIHC |
| KI67 |  | anti KI67 |  |  | SP6 |  | Abcam | ab15580 | mIHC |
| PANCK |  | anti pan cytokeratin |  |  | AE1/AE3 |  | Abcam | ab27988 | mIHC |
| ASMA |  | anti alpha SMA |  |  | 1A4 |  | Dako | IS611 | mIHC |
| CD320NKP46 | | anti CD3/anti CD30/anti NKp46 | |  | SP7/SP32/195314 |  | Thermo/Abcam/R&D Bio | MA1-90582/ab64088/MAB1850500 | mIHC |
| CD4 |  | anti CD4 |  |  | SP35 |  | Abcam | ab227709 | mIHC |
| CD56 |  | anti CD56 |  |  | 123C3 |  | Santa Cruz | sc-7326 | mIHC |
| CSF1R |  | anti CSF1R |  |  | SP211 |  | Abcam | ab183316 | mIHC |
| DCSIGN |  | anti DCSIGN |  |  | DC-28 |  | Santa Cruz | sc-65740 | mIHC |
| GATA3 |  | anti GATA3 |  |  | L50-823 |  | BioCare Medical | CM405A | mIHC |
| ICOS |  | anti ICOS |  |  | SP98 |  | LifeSpan Bio | LS-C210350 | mIHC |
| IDO |  | anti IDO |  |  | 1F8.2 |  | EMD Millipore | MAB10009 | mIHC |
| IL10 |  | anti IL10 |  |  | LS-B7411 |  | LifeSpan Bio | LS-B7411 | mIHC |
| NUCLEI |  | hematoxylin |  |  |  |  | Dako | S330130-2 | mIHC |
| TRYPTASE | | anti Tryptase |  |  | AA1 |  | Abcam | ab2378 | mIHC |

**Supplemental Table 5.**

| Pathway | Predictor | Weight | Count |
| --- | --- | --- | --- |
| Apoptosis | CASPASE7CLEAVEDD198 | 1 | 1 |
| Apoptosis | CASPASE8 | 1 | 1 |
| AR | AR | 1 | 1 |
| AR | AR | 1 | 1 |
| BCL2 | BCL2 | 1 | 1 |
| BH3_Balance | BCL2 | -1 | 1 |
| BH3_Balance | BAK | 1 | 1 |
| BH3_Balance | BAX | 1 | 1 |
| BH3_Balance | BID | 1 | 1 |
| BH3_Balance | BIM | 1 | 1 |
| BH3_Balance | Mcl-1 | -1 | 1 |
| BH3_Balance | BAD_pS112 | -1 | 1 |
| BH3_Balance | BCLXL | -1 | 1 |
| BH3_Balance | CIAP | -1 | 1 |
| Cell_cycle_progression | CYCLINB1 | 1 | 1 |
| Cell_cycle_progression | PLK1 | 1 | 1 |
| Cell_cycle_progression | CDK1_pT14 | 1 | 1 |
| Cell_cycle_progression | CHK1 | 1 | 1 |
| Cell_cycle_progression | cdc25C | 1 | 1 |
| Cell_cycle_progression | RB_pS807S811 | 1 | 1 |
| Cell_cycle_progression | P21 | -1 | 1 |
| Cell_cycle_progression | P27_pT198 | -1 | 1 |
| Cell_cycle_progression | CYCLIND1 | -1 | 1 |
| G2M_Checkpoint | CDK1_pT14 | 1 | 1 |
| G2M_Checkpoint | Histone-H3 | 1 | 1 |
| G2M_Checkpoint | H2AX_pS140 | 1 | 1 |
| G2M_Checkpoint | ATM_pS1981 | 1 | 1 |
| G2M_Checkpoint | ATR_pS428 | 1 | 1 |
| G2M_Checkpoint | CDK1_pY15 | 1 | 1 |
| G2M_Checkpoint | CHK1_pS296 | 1 | 1 |
| G2M_Checkpoint | CHK2_pT68 | 1 | 1 |
| G2M_Checkpoint | Wee1_pS642 | 1 | 1 |
| G2M_Checkpoint | CHK1_pS345 | 1 | 1 |
| G2M_Checkpoint | H2AX_pS139 | 1 | 1 |
| G2M_Checkpoint | RPA32_pS4_S8 | 1 | 1 |
| G0_G1 | CYCLINB1 | -1 | 1 |
| G0_G1 | P21 | 1 | 1 |
| G0_G1 | P27_pT198 | 1 | 1 |
| G0_G1 | CYCLIND1 | 1 | 1 |
| G0_G1 | BRD4 | -1 | 1 |
| G1_S | BRD4 | 1 | 1 |
| G1_S | CYCLINE1 | 1 | 1 |
| G2_M | CYCLINB1 | 1 | 1 |
| G2_M | PLK1 | 1 | 1 |
| G2_M | CDK1_pT14 | 1 | 1 |
| G2_M | cdc25C | 1 | 1 |
| G2_M | RB_pS807S811 | 1 | 1 |
| Hormone_receptor | AR | 1 | 1 |
| Hormone_receptor | ERALPHA | 1 | 1 |
| Hormone_receptor | ERALPHA_pS118 | 1 | 1 |
| Hormone_receptor | PR | 1 | 1 |
| Hormone_signaling_Breast | BCL2 | 1 | 1 |
| Hormone_signaling_Breast | ERALPHA_pS118 | 1 | 1 |
| Hormone_signaling_Breast | GATA3 | 1 | 1 |
| Immune | LCK | 1 | 1 |
| Immune | ZAP-70 | 1 | 1 |
| Immune_Checkpoint | B7-H4 | 1 | 1 |
| Immune_Checkpoint | PDL1 | 1 | 1 |
| Notch | Jagged1 | 1 | 1 |
| Notch | Notch3 | 1 | 1 |
| Notch | TAZ | 1 | 1 |
| Notch | YAP | 1 | 1 |
| Notch | YAP_pS127 | -1 | 1 |
| PI3K_Akt | P27_pT198 | 1 | 1 |
| PI3K_Akt | AKT_pS473 | 0.5 | 0.5 |
| PI3K_Akt | AKT_pT308 | 0.5 | 0.5 |
| PI3K_Akt | GSK3ALPHABETA_pS21S9 | 1 | 1 |
| PI3K_Akt | PRAS40_pT246 | 1 | 1 |
| PI3K_Akt | PTEN | -1 | 1 |
| RAS_MAPK | BRAF_pS445 | 1 | 1 |
| RAS_MAPK | CJUN_pS73 | 1 | 1 |
| RAS_MAPK | CRAF_pS338 | 1 | 1 |
| RAS_MAPK | MEK1_pS217S221 | 1 | 1 |
| RAS_MAPK | P38_pT180Y182 | 1 | 1 |
| RAS_MAPK | P38MAPK | 1 | 1 |
| RAS_MAPK | P90RSK_pT359S363 | 1 | 1 |
| RAS_MAPK | YB1_pS102 | 1 | 1 |
| RAS_MAPK | MAPK_pT202Y204 | 1 | 1 |
| RTK | CMET_pY1235 | 1 | 1 |
| RTK | EGFR_pY1173 | 1 | 1 |
| RTK | HER2_pY1248 | 1 | 1 |
| RTK | HER3_pY1289 | 1 | 1 |
| RTK | IGF1R_pY1135Y1136 | 1 | 1 |
| RTK | IRS1 | 1 | 1 |
| RTK | SRC_pY527 | 1 | 1 |
| RTK | SHP2_pY542 | 1 | 1 |
| RTK | SHC_pY317 | 1 | 1 |
| RTK | SRC_pY416 | 1 | 1 |
| TSC_mTOR | RB_pS807S811 | 1 | 1 |
| TSC_mTOR | MTOR_pS2448 | 1 | 1 |
| TSC_mTOR | P70S6K_pT389 | 1 | 1 |
| TSC_mTOR | RICTOR_pT1135 | 1 | 1 |
| TSC_mTOR | S6_pS235S236 | 0.5 | 0.5 |
| TSC_mTOR | S6_pS240S244 | 0.5 | 0.5 |
| Tumor_Content | LCK | -1 | 1 |
| Tumor_Content | BETACATENIN | 1 | 1 |
| Tumor_Content | CLAUDIN7 | 1 | 1 |
| Tumor_Content | ECADHERIN | 1 | 1 |
| Tumor_Content | RBM15 | 1 | 1 |
| Tumor_Content | EPPK1 | 1 | 1 |
| Tumor_Content | CAVEOLIN1 | -1 | 1 |
| Tumor_Content | COLLAGENVI | -1 | 1 |
| Tumor_Content | MMP2 | -1 | 1 |
| Tumor_Content | PAI1 | -1 | 1 |
| HER2 | HER2 | 1 | 1 |
| HER2_pY1248 | HER2_pY1248 | 1 | 1 |
| HER3 | HER3 | 1 | 1 |
| HER3_pY1289 | HER3_pY1289 | 1 | 1 |
| GATA3 | GATA3 | 1 | 1 |
| Cyclind1 | CYCLIND1 | 1 | 1 |
| ERALPHA | ERALPHA | 1 | 1 |
| ERALPHA_pS118 | ERALPHA_pS118 | 1 | 1 |
| STAT3PY705 | STAT3_pY705 | 1 | 1 |
| JAK/STAT | STAT3_pY705 | 1 | 1 |
| JAK/STAT | EMA | 1 | 1 |
| JAK/STAT | IL-6 | 1 | 1 |
| JAK/STAT | JAK2 | 1 | 1 |
| JAK/STAT | Stat1_pY701 | 1 | 1 |
| JAK/STAT | STAT5ALPHA | 1 | 1 |
